# MLL1 regulates cytokine-driven cell migration and metastasis

**DOI:** 10.1101/2022.10.18.512715

**Authors:** Praful R. Nair, Ludmila Danilova, Estibaliz Gómez-de-Mariscal, Dongjoo Kim, Rong Fan, Arrate Muñoz-Barrutia, Elana J. Fertig, Denis Wirtz

## Abstract

Cell migration is a critical requirement for cancer metastasis. Cytokine production and its role in cancer cell migration have been traditionally associated with immune cells in the tumor microenvironment. MLL1 is a histone methyltransferase that controls 3D cell migration via the secretion of cytokines, IL-6 and TGF-β1, by the cancer cells themselves. *In vivo*, MLL1 depletion reduced metastatic burden and prolonged survival. MLL1 exerts its effects with its scaffold protein, Menin. Mechanistically, the MLL1-Menin interaction controls actin filament assembly via the IL-6/pSTAT3/Arp3 axis and acto-myosin contractility via the TGF-β1/Gli2/ROCK1/2/pMLC2 axis, which regulate dynamic protrusion generation and 3D cell migration. MLL1 also regulates cell proliferation via mitosis-based and cell cycle-related pathways. Combining an MLL1-Menin inhibitor with Paclitaxel, a standard chemotherapeutic, abrogated tumor growth and metastasis in a syngeneic model. These results highlight the potential of targeting the MLL1 in metastasis prevention and its potential to be combined with currently administered chemotherapeutics.

**Statement of Significance:** We identify MLL1 as being vital to metastasis, which causes the vast majority of cancer-related deaths. MLL1 controls cell migration, a requirement for metastasis, by regulating the secretion of cytokines. MLL1 inhibition lowers metastatic burden independent of its impact on primary tumor growth, highlighting its anti-metastatic potential in TNBC.

## Introduction

Cell migration, particularly in three-dimensional (3D) environments, is a critical requirement for metastasis - the spread of cancer cells from a primary tumor to distant sites (*1*). Despite causing the vast majority of cancer-related deaths, targeting metastasis remains challenging clinically (*2*). Cancer cell migration can be triggered and maintained by elevated levels of cytokines, thus playing a vital role in cancer metastasis (*3, 4, 13–22, 5–12*). In solid tumors, these cytokines are presumed to be predominantly secreted by immune cells in the tumor microenvironment, although recent studies have shed light on the production of these cytokines by cancer cells themselves (*3, 4, 6, 9, 23–27*). While the signaling cascades that regulate cell migration have been studied extensively (*28*), the epigenetic regulation of cell migration is still poorly understood (*29*). Despite identifying some epigenetic regulators of cancer cell migration, there is currently no known epigenetic regulator of cytokine-driven cell migration (Fig. S1a). This is clinically significant as inhibiting epigenetic drivers of cancer-cell migration present potential new targets and treatment venues targeting metastasis specifically. An epigenetic factor that is strongly correlated with cell migration-related genes in The Cancer Genome Atlas (TCGA) database is Mixed-Lineage Leukemia 1 (MLL1 or MLL, gene name *KMT2A*). MLL1 is a histone methyltransferase, which along with its scaffold protein Menin (*30*), is integral to methylation at the histone 3 lysine 4 (H3K4) site (*30–32*). One of these marks, trimethylation of H3K4 (H3K4me3), is a well-established mark for activation of transcription (*33*). MLL1 has not yet been implicated in cell migration or metastasis.

Here, we show that the MLL1-Menin interaction is essential for the migration of cancer cells and that the disruption of this interaction impairs cell migration and metastasis. In 3D *in vitro* models of migration, MLL1 depletion decreases migration by reducing protrusion generation via downregulation of the cytokine IL-6 and downstream proteins that activate actin filament assembly via an IL-6/STAT3/Arp2/3 pathway. Pharmacological inhibitors permit to finely tune the inhibition of MLL1-Menin interaction and reveal a non-linear mechanism of regulation of cell migration. A deep inhibition of the MLL1-Menin interaction also leads to reduced myosin II-based contractility via a TGF-β1/GLI2/ROCK1/2 pathway. These studies reveal that MLL1 plays a two-pronged role in cell migration by controlling both actin filament assembly and myosin-based contractility. Our *in vivo* studies support these mechanisms of cell migration and reveal that mice bearing orthotopic breast MLL1-depleted tumors grow significantly more slowly and induce lower metastatic burden than control tumors, leading to a significantly longer survival. Reduced lung metastatic burden was observed even after accounting for the difference in the growth rate of primary tumors. Finally, in a syngeneic orthotopic mouse TNBC model, concurrent administration of the MLL1-Menin inhibitor and Paclitaxel led to a near-arrest of tumor growth and the extremely low metastatic burden. Overall, our studies show that MLL1 is an epigenetic regulator of 3D cancer cell migration, which exerts its effects by controlling the production of the key cytokines IL-6 and TGF-β1. Our *in vivo* studies demonstrate a potential anti-cancer therapy that can reduce metastatic burden via two separate molecular mechanisms.

## Results

### TCGA analysis suggests MLL1 is an epigenetic regulator of cell migration

Triple negative breast cancer (TNBC) represents 15-20% of all breast cancer cases (*34*), and is an aggressive subtype with a poor prognosis (*35, 36*) and a high incidence of metastases (*37–39*). Patients with TNBC also relapse more frequently than other breast-cancer subtypes (*39*), and despite administration of optimal chemotherapy regimens, only 30% of metastatic TNBC patients show >5-year survival (*40*), hinting at the inadequacy of current standard-of-care treatments. Additionally, targeting metastasis is a formidable challenge and there is a dearth of anti-metastatic therapies in the clinic (*41*). Hence, TNBC was chosen as the key focus and testbed for this study due to its propensity for metastatic spread.

To identify potential epigenetic regulators of cell migration in TNBC, correlation coefficients of cell migration-related genes (*ROCK1, ROCK2, LIMK1, RAC1, RHOA, CDC42, MYLK, WASF3, ACTR2,* and *ACTR3*) were calculated from the PanCanAtlas (TCGA) basal breast cancer dataset for 715 epigenetic genes (*42*). These epigenetic genes were then ranked based on the summation of correlation coefficients (Fig. 1a) and the top 10 epigenetic factors (*ASXL2, REST, PPP4R2, RPS6KA3, CLOCK, USP12, KMT2A/*MLL1*, RAD54L2, PBRM1,* and *MYSM1*) were examined as potential candidates. Only three of these genes (*ASXL2, KMT2A/*MLL1, and *PBRM1*) were histone modifiers or chromatin remodelers (Fig. S1b) and only *KMT2A/*MLL1 has been the subject of cancer clinical trials; albeit these trials are in acute myeloid leukemia and are neither in solid tumors nor focused on metastatic prevention (Fig. S1b). Additionally, MLL1 was ranked much higher than previously reported epigenetic factors of cell migration in Fig. S1a (Fig. S1c). Together, these findings indicate that MLL1 is a potential regulator of motility in cancer cells and has high translational potential owing to ongoing clinical trials

**Figure 1.**
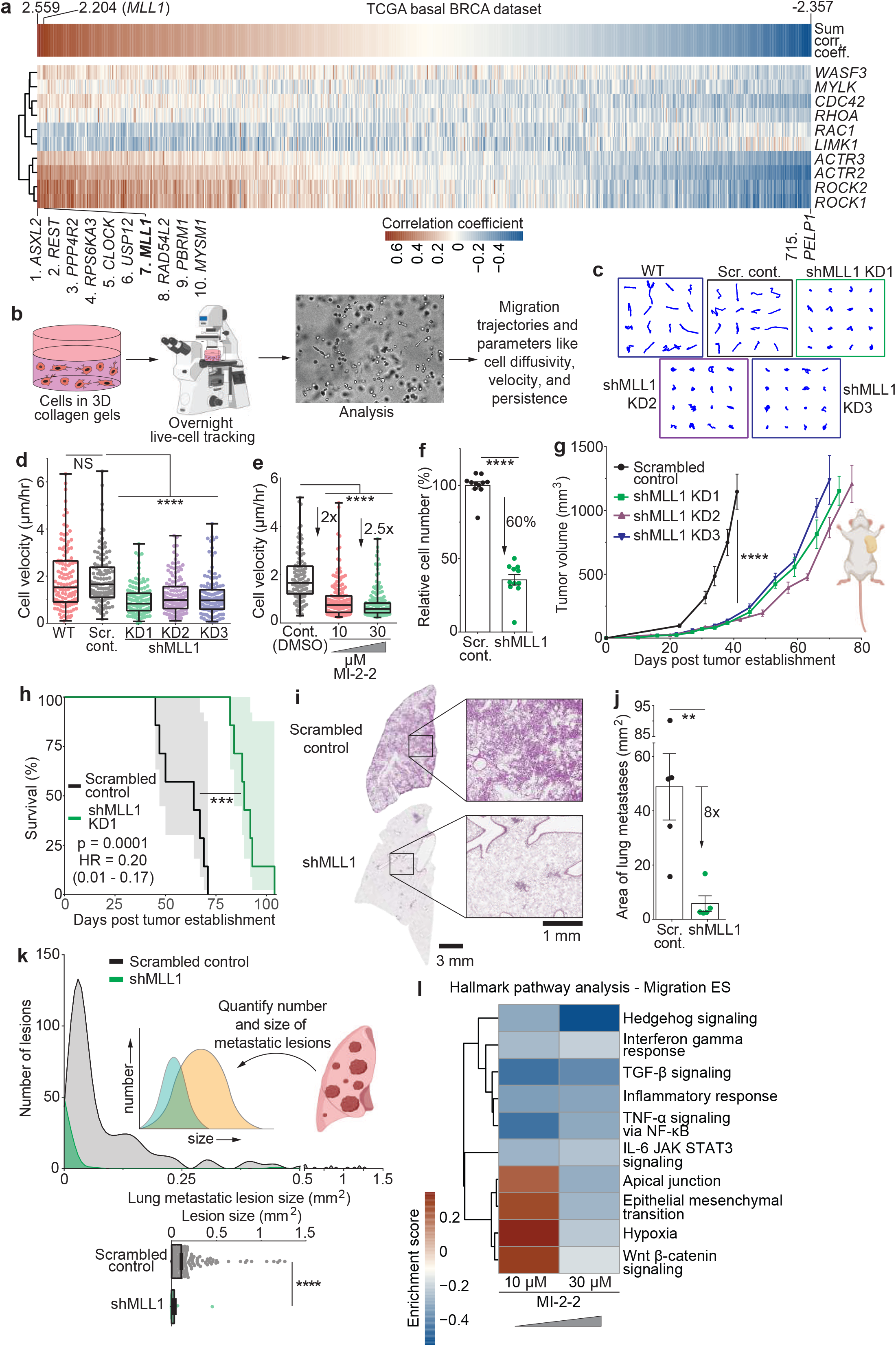
MLL1-Menin interaction regulates cancer cell migration, proliferation, and tumor progression. **(a)** Epigenetic factors in the TCGA basal breast cancer dataset were queried for correlation with genes reported to be essential for cell migration genes, including RHOA, RAC1, ROCK1, ROCK2, MYLK, CDC42, LIMK1, ACTR2,ACTR3, and WASF3. MLL1 was the seventh-highest correlated gene among the 715 epigenetic factors and had high correlation with genes involved in myosin-based contractility. **(b)** Cells were embedded in 3D collagen gels and cells were tracked overnight using live-cell phase contrast microscopy. Resulting videos were analyzed to obtain cell trajectories, velocity, and diffusivity using the APRW cell-migration model. **(c)** shMLL1 cells exhibited shorter trajectories than scrambled non-targeting shRNA control and wild type cells. **(d)** shMLL1 cells had lower velocity, compared to control and WT cells. Velocities are depicted in a box-and-whisker plot, with each dot corresponding to one cell. The upper limit of the box represents the 75^th^ percentile, lower limit represents the 25^th^ percentile, and the center line represents the median. The top and bottom whiskers stretch to the 99^th^ and 1^st^ percentile, respectively. n(cells) = 123(WT), 122(Scr. cont.), 128(shMLL1 KD1), 139(KD2), 132(KD3); one-way ANOVA (**** p < 0.0001, F (4, 639) = 26.65); two-tailed t-test (**** p(SC - sh) < 0.0001, NS p(WT - SC) = 0.8151). **(e)** Pharmacological inhibition of MLL1-Menin interaction using MI-2-2 decreased cell velocity to levels similar to shMLL1 cells. n(cells) = 112(WT), 175(10 µM), 183(30 µM); one-way ANOVA (**** p < 0.0001, F (2, 467) = 78.12); two-tailed t-test (**** p < 0.0001). **(f)** MLL1 depletion impairs cancer cell proliferation. n(wells) = 11(each condition); two-tailed t-test (**** p < 0.0001). **(g)** shMLL1 tumors grew slower compared to scrambled control tumors. n(mice) = 7(each condition); one-way ANOVA (**** p < 0.0001, F (3, 24) = 35.18). **(h)** NSG mice bearing shMLL1 tumors survive longer than mice bearing scrambled control tumors. n(mice) = 7(each condition); Log rank (Mantel–Cox) test (*** p = 0.0001, HR = 0.20, 95% C.I. = 0.01 – 0.17). **(i)** H&E staining of lungs show darker staining for scrambled control tumors indicative of higher metastatic burden compared to shMLL1 tumors. **(j)** Quantification of total metastatic burden per lung showed an increased metastatic burden in mice bearing scrambled control tumors compared to shMLL1 tumor bearing mice. n(lungs) = 5(each condition, 1 lung/mouse); two-tailed t-test (** p = 0.0091). **(k)** Histogram of lesion size of a single lung shows that decreased metastatic burden in shMLL1 lungs were due to both reduced number of lesions as well as their decreased size. n(lesions) = 367 (Scr. cont.), 61 (shMLL1); two-tailed t-test (**** p < 0.0001). **(l)** RNA-seq analysis of MLL1-Menin-inhibited cells revealed downregulation of key cell migration and metastasis-related pathways including the epithelial-mesenchymal transition, IL-6-JAK- STAT3 signaling, and TGF-β signaling pathways. Data in this figure was generated with MDA-MB-231 cells *in vivo* (g-k) or embedded in 3D collagen gels (b-f and l) except panel a (TCGA).

MLL1 is expressed in breast tumors at both the RNA (Fig. S1d, TCGA dataset via Human Protein Atlas) and the protein levels (Fig. S1e): 92% of the breast-cancer patients show medium-to-high expression of MLL1 in breast cancer tissue (IHC, stained with antibody CAB024270). MLL1 had little overall prognostic value in overall breast cancer survival (Fig. S1f; Human protein atlas, p value = 0.091, number of patients = 1075), or in later stage breast cancers (cancers that have invaded and spread to nearby lymph nodes or distant organs (*43*)); but was prognostic in earlier stage breast cancer (that have yet to spread from the original tumor site). Key motility genes *ROCK1* and *ROCK2* showed similar prognostic trends (Fig. S1g); as did breast cancer metastasis suppressor, BRMS1 (*44*) (fig. S1h, although earlier stages show better prognosis with higher BRMS1 expression as BRMS1 expression suppresses metastasis). GSEA analysis showed that MLL1 expression was positively correlated with two metastasis-related gene sets: genes associated with a poor prognosis and genes expressed in EMT mammary tumors (Fig. S2a-b). Finally, the expression of MLL1 in two metastatic TNBC cell lines (MDA-MB-231 and SUM-159) was compared against non-tumorigenic breast epithelial cells (MCF- 10A). The metastatic TNBC cells expressed higher levels of MLL1 than normal breast epithelial cells (Fig. S2c). In sum, MLL1 is prognostic in breast cancers that have yet to metastasize, and its expression is higher in breast carcinoma cells compared to non-cancer breast epithelial cells.

### MLL1-Menin interaction regulates 3D cell migration, proliferation, and tumor progression

Since MLL1 was identified as potential epigenetic regulator of cell migration, which is essential for metastasis, we next sought to determine if depleting MLL1 impacted cell migration and/or metastasis. MLL1 was depleted in human TNBC (MDA-MB-231) cells via shRNA (Fig. S2d). Cells transduced with non-targeting scrambled shRNA (scrambled control), MLL1-depleted (shMLL1) cells, and wild-type (WT) cells were embedded in 3D collagen matrices, which mimic the 3D collagen-rich environment of the stromal matrix (*45*). Embedded cells were subjected to overnight live-cell tracking (Fig. 1b) and migration-related parameters were computed using a custom code (*46*). Live-cell microscopy permits the tracking of individual cells, fully dissociating any influence of proliferation on migration. Scrambled control cells and WT cells were highly motile, as evident by longer trajectories (Fig. 1c), whereas all three MLL1 knockdowns were significantly less motile, indicating that MLL1 knockdown impaired cell migration. These observations were reflected when extracting cell velocities from cell trajectories (Fig. 1d): control cells moved twice as fast as shMLL1 cells. The persistence time of migration (a measure of how long a cell takes to change direction significantly), speed, and cell diffusivity (an intrinsic measure of motility that is independent of elapsed time) displayed similar trends (Fig. S2e-g). Additionally, Menin was depleted separately using shRNA (Fig. S2h): Menin-depleted cells displayed a loss in migration and decreased velocity similar to those displayed by MLL1-depleted cells (Fig. S2i).

MLL1 was chosen over other epigenetic candidates as there are drugs against it currently being evaluated in clinical trials (Fig. S1b). In addition to the translational potential, pharmacological inhibition also allows for a finer modulation of MLL1-Menin interaction than what is possible with a knockdown. Inhibition of the MLL1-Menin interaction (referred to interchangeably as MLL1-Menin inhibition or MLL1 inhibition) was achieved using the drug MI-2-2 (*47*), which led to decrease in cell motility similar to that observed with shMLL1 cells (Fig. 1e). Similarly, MLL1-Menin inhibition reduced cell diffusivity (Fig. S2j) and shortened cell trajectories (Fig. S2k) compared to untreated (DMSO) control cells. A dose response of the inhibition of cell migration by MI-2-2 revealed the IC50 for cell migration to be 5 µM (Fig. S2l). These observations were validated in another 3D *in vitro* system: tumor spheroids (Fig. S2m). These double-layered spheroids consist of a core of cancer cells in Matrigel surrounded by an outer collagen layer. This system allows for the analysis of: (1) invasion through a basement membrane-like material (Matrigel) and (2) migration through collagen I (in yellow). MLL1-Menin inhibition reduced the dissemination of cells from the inner core into the outer collagen-rich corona (Fig. S2n, invasive front traced in orange). These results were confirmed in four other TNBC cell lines: BT-549, SUM-149, SUM- 159 (all human TNBC), and 4T1 (mouse TNBC) (Fig. S2o), as well as in HT-1080 fibrosarcoma cells (Fig. S2p). Thus, MLL1-Menin interaction controls cell migration in a wide panel of cells.

MLL1 is a component of the COMPASS complex that incorporates other transcription factors (*48*). WDR5, a COMPASS member (*48*) and an MLL1-binding protein (*49*), is a part of chromatin-regulatory complexes, and has been shown to play a role in cancer progression (*50, 51*). Disruption of MLL1- WDR5 interaction (via OICR-9429, (*52*)) reduced cell migration (Fig. S2q) to levels observed with MLL1- Menin inhibition or MLL1 depletion. MLL1 has been implicated as a driver in a subset of leukemias, where MLL1-fusion proteins drive leukemogenesis (*53*). In MLL1-fusion leukemias, the H3K79-specific methyltransferase Dot1L, which is not part of COMPASS, is necessary for progression and maintenance in MLL1-fusion leukemias (*54–56*). Dot1L inhibitors such as EPZ-5676 have shown efficacy in MLL1-rearranged acute myeloid leukemia and is being investigated in a phase Ib/II clinical trial (NCT03724084) (*57, 58*). Treatment of breast cancer cells with Dot1L inhibitor at multiple doses did not alter cell migration (Fig. S2r). Additionally, depletion or inhibition of MLL1 in TNBC cells did not decrease the expression of Hox genes reported to be downstream of MLL1 in MLL-fusion leukemias (HOXA7, HOXA9, HOXA10, HOXA11, MEIS1, and PBX3 (*59, 60*)), whose downregulation has been shown to be a hallmark of MLL1-Menin inhibition in MLL-fusion leukemias (Fig. S3a-d). Hence, the underlying biology of MLL1-based cell migration is different from that of leukemogenesis of MLL1-fusion driven leukemias. MLL1-Menin-based regulation of cell migration was also independent of 3D cell density (20-250 cells/mm^3^, Fig. S3e) and collagen density in 3D gels (1-3 mg/ml collagen, Fig. S3f). In sum, perturbation of MLL1’s interaction with COMPASS members WDR5 and Menin reduced 3D cell migration and the mechanism of MLL1-based cell migration is distinct from MLL1’s role in MLL1-fusion leukemias.

In addition to regulating cell migration, MLL1-Menin interaction also plays a role in controlling proliferation. MLL1 depletion decreased proliferation by 60% (Fig. 1f). These *in vitro* observations were validated in orthotopic TNBC mouse models to assess the role of MLL1-Menin mediated migration on tumor progression (tumor growth and metastasis). Further validation is shown below in syngeneic mouse models. *In vivo* modeling demonstrated that MLL1-depleted tumors grew significantly more slowly (Fig. 1g and S3g) and mice bearing shMLL1 tumors exhibited increased survival (Fig. 1h and S3h). MLL1 depletion prolonged the survival (median survival time of 89 days for shMLL1 compared to 64 days for scrambled control) and had a substantially reduced hazard ratio (HR) of 0.20. Thus, depletion of MLL1 prolongs survival *in vivo*. However, survival in mouse models is primarily impacted by proliferation at the primary tumor site. To assess the impact of MLL1 on metastasis specifically, mice bearing scrambled control and shMLL1 tumors were sacrificed at a set time point, six weeks after tumor establishment (Fig. S3i). Lung sections showed decreased metastatic nodules in shMLL1 tumor bearing mice, evident in lighter hematoxylin and eosin (H&E) staining in lungs (Fig. 1i). Quantification of total lung metastatic burden showed an eight-fold decrease in metastatic burden for mice-bearing shMLL1 tumors compared to mice-bearing control tumors (Fig. 1j). The size distribution of metastatic lesions was also determined for lungs shown in Fig. 1i. Control-bearing lungs not only had many more metastatic lesions (Fig. 1k, bottom), but these lesions were much larger than shMLL1-bearing lungs (Fig. 1k indicated by a thick histogram tail for control mice). Thus, MLL1 regulates tumor growth and metastasis *in vivo*. These studies were purely mechanistic, with an emphasis on validation of the role of MLL1 in tumor progression. The translation of these findings to pre-clinical *in vivo* models (Fig. 7), will demonstrate abrogation of tumor growth and metastasis when combining MLL1 inhibition with standard-of-care treatment.

Finally, we have done an exhaustive assessment of the role of potential off-target effects in our observations and found none. Disruption of the MLL1-Menin interaction was confirmed using other newer MLL1-Menin inhibitors, MI-503 (*59*) (Fig. S3j-k) and VTP50469 (*60*) (Fig. S3l). A more potent inhibitor permits disruption of the MLL1-Menin interaction at a lower dose, minimizing potential off-target effects. Both drugs reduced cell migration to the same extent, albeit at an order of magnitude lower dose. To test for potential off-target effects of MI-2-2, a weak inhibitor of MLL1-Menin interaction, MI-nc, was used as a negative control. MI-nc has a similar structure to MI-2-2, but binds much more weakly to MLL1-Menin and does not inhibit MLL1-Menin interaction (*61*). MI-nc treatment did not impact cell migration (Fig. S4a), suggesting that inhibition of cell migration by MI-2-2 or MI-503 was due to the disruption of the MLL1-Menin interaction rather than off-target effects. In non-cancerous MCF10A breast epithelial cells (Fig. S2c), MLL1 inhibition had a markedly lower impact on cell migration (Fig. S4b) and proliferation (Fig. S4c-d) compared to TNBC cells, hinting at a potentially weaker impact of MLL1-Menin inhibition on non-cancerous cells compared to tumor cells. Additionally, decreased in proliferation following MLL1-Menin inhibition was not accompanied by a significant increase in apoptosis (Fig. S4e-f), indicating that decreased cell numbers and/or reduced cell motility after MLL1 depletion was not due to decreased cell viability. The observed pharmacological inhibition of cell migration was also readily and fully reversible upon withdrawal of drug treatment. Cells were pre-treated with MLL1 inhibitor for two days before being seeded and subjected to further MLL1 inhibition (post-treatment) (Fig. S4g). Pre-treated cells without any post-treatment showed a full recovery in cell velocity and had the same motility as untreated cells within two-three days (Fig. S4h). Both pre-treated and non-pretreated cells receiving post-treatment showed similarly low motility. Similar results were also observed in HT-1080 fibrosarcoma cells. MLL1-Menin inhibition also did not further reduce the motility of shMLL1 cells (Fig. S4i), indicating that concurrent MLL1-Menin inhibition and MLL1 knockdown could not further reduce cell migration compared to either inhibition or depletion alone. This implies that MLL1- Menin inhibitors had no effect when MLL1 was absent in cells, and – together with MI-nc results – the observed results were highly unlikely to be due to off-target effects. Thus, decreased cell migration via pharmacological inhibition was due to the disruption of MLL1-Menin interaction rather than off-target effects, and this effect was fully reversible following withdrawal of drug treatment. Administration of MLL1-Menin inhibitors was associated with low toxicity (indicated by low apoptosis rates), which is consistent with other published results that report low toxicity *in vivo*, even when administered at up to 100 mg/kg (*59*). Further discussion pertaining to the assessment of off-target effects can be found in the methods section.

In sum, MLL1-Menin interaction regulates the 3D migration of TNBC cells and drives TNBC metastasis *in vivo*.

### MLL1-Menin regulates IL-6 production and cell protrusion generation

We next sought to determine the mechanism behind MLL1-mediated cell migration. Transcriptomic analysis showed that MLL1-Menin inhibition led to changes in key migration-related gene sets (Fig. 1l). In particular, major cytokine-based cell-migration-related pathways such as IL-6/JAK/STAT3 signaling (*3*), TGF-β signaling (*62*) and TNF-α signaling via NF-κB (*14*) were downregulated following lower dose of MLL1 inhibitor (10 µM MI-2-2). Higher dose (30 µM) led to generally a much deeper repression of these pathways, as well as of other metastasis-related pathways such as epithelial mesenchymal transition (*63*), apical junction transition (*64*), hypoxia (*65, 66*), and Wnt β-catenin signaling pathways (*67*).

To determine whether MLL1-menin interaction affects cell migration via the secretion of any soluble factors such as cytokines, conditioned medium was collected from scrambled control cells embedded in 3D collagen gels (Fig. 2a). This conditioned medium was then added to shMLL1 cells in collagen gels followed by overnight live cell tracking. Addition of conditioned medium to shMLL1 cells fully rescued the motility lost during MLL1 knockdown (trajectories in Fig. 2b and cell velocities in Fig. 2c). This conditioned medium did not further increase the motility of scrambled control cells. Conditioned medium was also collected from shMLL1 embedded in 3D collagen gels. The cell density was adjusted to be higher to account for the slower proliferation of shMLL1 cells and ensure similar cell numbers to scrambled control cells. Addition of shMLL1 conditioned medium to scrambled control or shMLL1 cells did not increase cell velocity compared to fresh medium (Fig. S4j). Cells in 3D collagen gels display increased migration for increasing cell density (i.e. for increased number of cells per unit volume) (*3*). Cells at low density (LD, 10 cells/mm^3^) move significantly less compared to the standard cell density utilized in prior experiments (100 cells/mm^3^, high density - HD), but their motility can be increased by the addition of conditioned medium from HD cells. To further confirm that MLL1 mediated cell motility is mediated via secretion of soluble factors, LD cells were incubated with conditioned medium from scrambled control and shMLL1 cells (Fig. S4k). Conditioned medium was collected after three days, allowing ample time for the secretion of cytokines critical for cell motility. Addition of conditioned medium from scrambled control cells to LD cells increased their motility (red) to levels comparable to cells at HD (black), evident in both the trajectories (Fig. S4l) and cell velocities (Fig. S4m). However, conditioned medium from shMLL1 cells had no impact on the motility of cells at LD (green), indicating that conditioned medium from shMLL1 cells lacks secreted factors key for cell migration. In sum, MLL1- Menin regulates cell migration occurs via the secretion of soluble factors.

**Figure 2.**
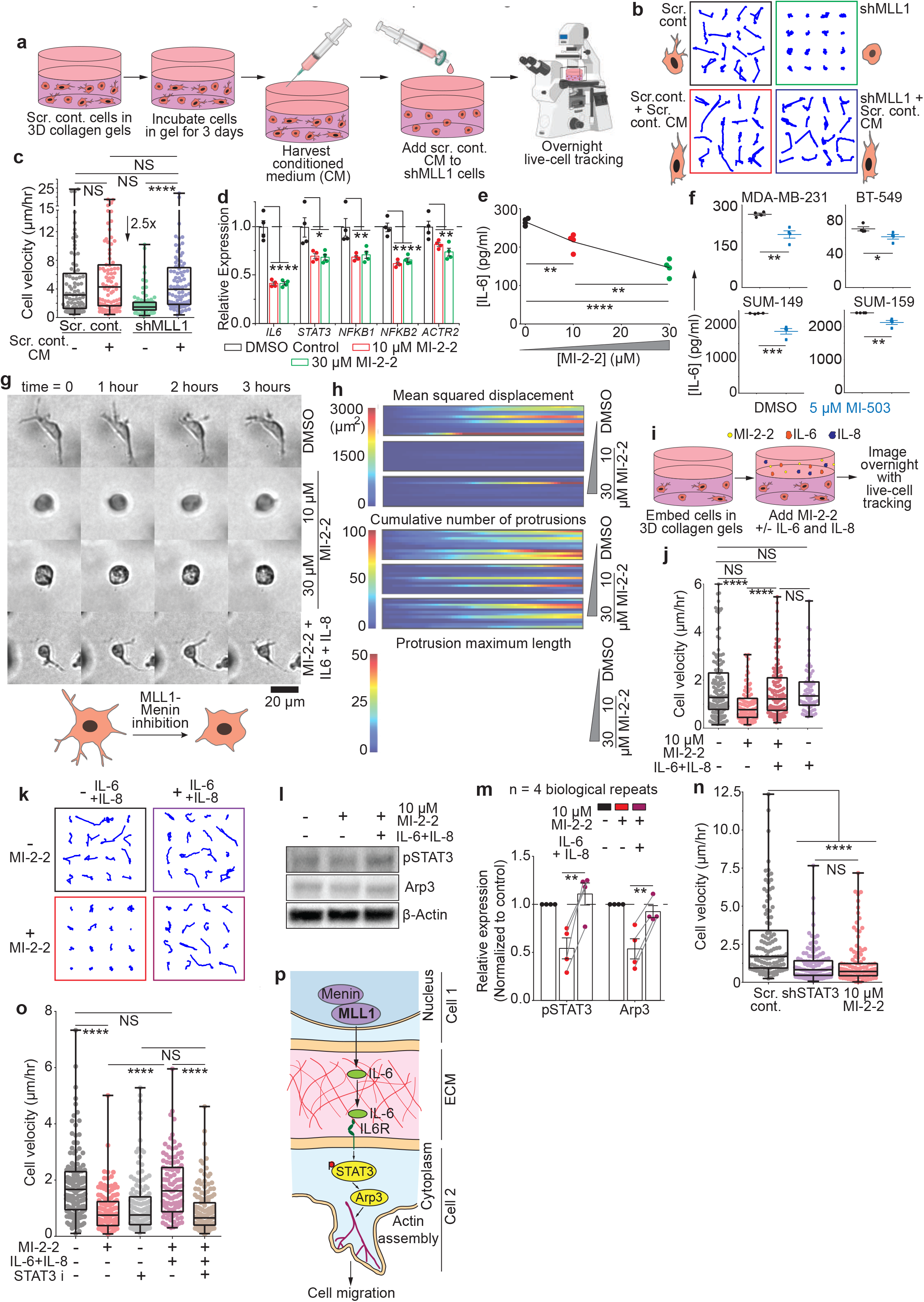
MLL1-Menin interaction regulates cell protrusion generation via IL-6/pSTAT3 signaling. **(a)** Conditioned medium was collected from scrambled control cells and added to shMLL1 or scrambled control cells. Conditioned medium from scrambled control cells fully rescued cell migration of shMLL1 cells as seen in **(b)** trajectories and **(c)** velocities. n(cells) = 81(Scr. cont.), 81(Scr. Cont.+CM), 89 (shMLL1), 85 (shMLL1+CM); one-way ANOVA (**** p < 0.0001, F (3, 332) = 13.68); two-tailed t-test (**** p(sh - sh+CM) < 0.0001, NS p(SC - sh+CM) = 0.8563, NS p(SC+CM - sh+CM) = 0.4392, NS p(SC - SC+CM) = 0.3979). **(d)** MLL1-Menin inhibition leads to downregulation of key genes in the IL- 6/JAK/STAT3 signaling pathway assessed via qRT-PCR. MLL1-Menin inhibition by n(wells) = 4(each condition); one-way ANOVA (IL6: **** p < 0.0001, F (2, 9) = 61.73; STAT3: * p = 0.0104, F (2, 9) = 7.905; NFKB1: ** p = 0.0075, F (2, 9) = 8.858; NFKB2: **** p < 0.0001, F (2, 9) = 47.67 ; ACTR2: ** p = 0.0073, F (2, 9) = 8.918). **(e)** MI-2-2 [n(wells) = 4(each condition); one-way ANOVA (**** p < 0.0001, F (2, 9) = 41.11); two-tailed t-test (** p(0 - 10) = 0.0049, **** p(0 - 30) < 0.0001, ** p(10 - 30) = 0.005)] or **(f)** MI-503 decreased IL-6 production TNBC cell lines. n(wells) = 4(each condition); two-tailed t-test (** p(MDA-MB-231) = 0.0020, * p(BT-549) = 0.0420, *** p(SUM-149) = 0.0007, ** p(SUM-159) = 0.0030,). **(g)** Phase contract images of cells treated with MLL1-Menin inhibitor show decreased generation of protrusions. Protrusion generation can be rescued by adding IL-6+IL-8 to cells treated with MLL1-Menin inhibitor, MI-2-2. Scale bar 20 µm. **(h)** A temporal heatmap of protrusion-related parameters show that the cumulative number of protrusions generated per cell and the maximum length of protrusion generated by a cell were reduced by MLL1-Menin inhibition, which coincided with a decrease in cell migration (mean squared displacement). Each row is one cell, and each block is one condition. **(i)** A rescue of cell motility was attempted by supplementing cells with IL-6/8 on top of MLL1 inhibition by 10 µM MI-2-2. IL-6/8 supplementation fully rescued cell motility despite continuing inhibition of the MLL1-Menin interaction, evident in **(j)** velocity and **(k)** trajectories. n(cells) = 135(Untreat.), 82(IL), 98(MI), 120 (IL+MI); one-way ANOVA (**** p < 0.0001, F (3, 431) = 9.233); two-tailed t-test (**** p(MI – MI+IL) < 0.0001, **** p(Untreat. – MI) < 0.0001, NS p(Untreat. – MI+IL) = 0.2905, NS p(MI+IL – IL) = 0.8257, NS p(Untreat. – IL) = 0.4676). **(l)** Western blotting on pSTAT3 (downstream of IL-6/8) and Arp2/3 (involved in actin assembly downstream of STAT3) shows inhibition and rescue following MLL1 inhibition and IL-6/8 supplementation, respectively. **(m)** Quantification of four biological repeats of IL- 6/8 rescue western blots emphasize that pSTAT3 and Arp3 levels are significantly reduced and subsequently restored. n(western blots) = 4 (biological replicates); paired two-tailed t-test (** p(pSTAT3) = 0.0056, ** p(Arp3) = 0.0098). **(n)** STAT3 knockdown reduces cell migration to levels observed with MLL1-Menin inhibition. n(cells) = 110(Scr. cont.), 121(shSTAT3), 121(MI-2-2); one-way ANOVA (**** p < 0.0001, F (2, 349) = 28.75); two-tailed t-test (**** p(SC – shSTAT3) < 0.0001, NS p(shSTAT3 – MI-2-2) = 0.6118, **** p(SC – MI-2-2) < 0.0001). **(o)** Motility rescued via IL-6/8 supplementation was lost by inhibiting STAT3 (using S3I-201), which lies downstream of IL-6/8. n(cells) = 146(Untreat.), 134(MI), 146 (S3I), 107(MI+IL), 147(MI+IL+S3I); one-way ANOVA (**** p < 0.0001, F (4, 675) = 30.68); two-tailed t-test (**** p(Untreat. - MI) < 0.0001, **** p(MI – MI+IL) < 0.0001, **** p(MI+IL – MI+IL+S3I) < 0.0001, NS p(Untreat. – MI+IL) = 0.9059, NS p(S3I - MI+IL+S3I) = 0.0535). **(p)** Schematic illustration of MLL1-Menin-based regulation of cell migration. MLL1-Menin interaction controls the production of IL-6, which binds to the IL-6 receptor and leads to phosphorylation of STAT3. pSTAT3 drives actin filament assembly via Arp2/3. Data in this figure was generated with MDA-MB-231 cells embedded in 3D collagen gels except panel f (MDA-MB-231, BT549, SUM-149, and SUM-159).

MLL1-inhibited cells showed downregulation of key members involved in the IL-6-STAT3-actin filament assembly pathway, including *IL-6, STAT3, and ACTR2* (Fig. 2d and S4n). The same IL-6-based actin assembly pathway was also downregulated in all three shMLL1 clones (Fig. S4o). Secretion of IL-6 was inhibited by MLL1-Menin inhibition in a dose-dependent manner (Fig. 2e), with IL-6 levels being halved. MI-503 also decreased IL-6 production in a range of TNBC cells (MDA-MB-231, BT-549, SUM- 149, and SUM-159) (Fig. 2f). Secretion of a wider panel of cytokines was tested using a multiplex cytokine assay. A multiplexed antibody barcode microarray chip was prepared against IL-6 and IL-8, which has also been shown to be essential for 3D cell migration (*3*). IL-6 production was decreased by MLL1-Menin inhibition as well as by MLL1 depletion (Fig. S4p). In contrast to IL-6, levels of IL-8 were only marginally reduced, mostly at high drug dose or in MLL1-depleted cells (Fig. S4q).

Cells produce dendritic protrusions to move effectively inside 3D collagen matrices, and hence, cell migration is tightly connected to cell morphology (*28, 68, 69*): more protrusions correlate with more effective 3D cell migration (*70, 71*). Cells embedded in 3D collagen gels produced large and dynamic protrusions, which was abrogated by MLL1-Menin inhibition (Fig. 2g, shown in phase contrast images over the course of 3 hours). MLL1-Menin inhibited cells supplemented with IL-6/8 displayed large protrusions, indicating that IL-6/8 supplementation rescued generation of cellular protrusions. A machine-learning algorithm was used to further analyze cell and protrusion morphology from phase-contrast images recorded during overnight live-cell tracking. MLL1-Menin inhibition reduced the number of cells that generated protrusions, particularly for lower dosage of MLL1 inhibitor (Figs. S4r and S5a). Decreased cell migration, measured as reduced mean squared displacements, was accompanied by a decrease in the number and maximum length of protrusions generated of cells (Fig. 2h). Thus MLL1- Menin regulated cell migration is accompanied by a reduction in cytokine (IL-6) secretion and protrusion generation, including size and number of protrusions.

### MLL1-Menin regulates actin filament assembly via an IL-6-pSTAT3-Arp3 axis

Exogenous IL-6 was supplemented to rescue cell motility following impairment of cell migration by MLL1-Menin inhibition. Since IL-8 levels were marginally reduced (Fig. S4q), it was also added to avoid being a bottleneck following IL-6 supplementation. Cells were embedded in collagen gels and treated with MI-2-2 as described previously (Fig. 2i). Rescue wells were supplemented with recombinant human IL-6 and IL-8 at reported concentrations of these cytokines in 3D collagen gels (*3*). MLL1- inhibited cells supplemented with IL-6 and IL-8 (referred to as IL-6/8, maroon, Fig. 2j) displayed increased cell motility compared to treated cells (red) and had velocities comparable to untreated control (black). Hence, cell motility lost by MLL1-Menin inhibition can be fully rescued by supplementation of IL-6/8. Addition of IL-6/8 to untreated cells (purple) did not further increase their velocity, indicating that IL-6/8-mediated cell migration reaches a plateau, just as observed with conditioned medium having minimal effect on scrambled control cells. As a positive control for cytokine supplementation, addition of IL-6/8 to LD cells increased their motility (Fig. S5b). These changes in cell velocity were reflected in cell trajectories (Fig. 2k), with supplementation of IL-6/8 lengthening cell trajectories despite continuing MLL1-inhibition. Cell motility lost by MI-503 could also be rescued by the supplementation of IL-6/8 (Fig. S5c). Thus, IL-6 lies downstream of MLL1-Menin interaction and mediates MLL1-based cell migration.

We then assessed the effect of MLL1-Menin inhibition and IL-6/8 supplementation on pSTAT3, a key transcription factor downstream of IL-6 (*72*). Levels of pSTAT3 and Arp3, which is involved in actin filament assembly (*73*), were halved with MLL1-Menin inhibition (Fig. 2l, left lane *vs*. middle lane), and supplementation of IL-6/8 increased levels of both proteins (Fig. 2l, right lane; quantified in Fig. 2m) back to levels observed for untreated cells. Arp3 is essential for protrusion generation (*3*) and decreased Arp3 is consistent with fewer and smaller protrusions generated by cells following MLL1- Menin inhibition (Figs. 2g-h, S4r, and S5a). Further, supplementation of IL-6/8 to MLL1-inhibited cells rescued protrusion generation (Fig. 2g). These decreases may partly explain the observed decrease in motility after MLL1-Menin inhibition. To verify that STAT3 was indeed responsible for cell migration, we depleted STAT3 via shRNA (Fig. S5d) and subjected the shSTAT3 cells to our overnight 3D cell migration assay (Fig. 2n). STAT3 depletion reduced cell velocity to levels observed with MLL1 inhibition, indicating that STAT3 plays a key role in cell migration. The same pathway involving *JAK2, STAT3, WASF3,* and *ARP2/3* was also responsible for reduced cell migration in BT549 TNBC cells following MLL1-Menin inhibition (Fig. S5e). To further confirm that increased STAT3 was responsible for increased cell motility downstream of IL-6/8 supplementation, we sought to determine if rescued motility (Fig. 2j) could be lost by inhibiting STAT3 (Fig. 2o). Cell motility was decreased with STAT3 inhibition (STAT3 i, grey) alone. Rescued movement in cells treated with MI-2-2+IL-6/8 (maroon) was lost by adding STAT3 inhibition on top of the rescue (brown, MI-2-2+IL-6/8+STAT3 i), leading to similar cell motility as either MLL1 inhibition (red) or STAT3 inhibition (grey) alone. Thus, STAT3 lies downstream of IL-6 and regulates IL-6-mediated cell migration.

Taken together, our results establish that IL-6 lies downstream of MLL1-Menin interaction and that STAT3 lies downstream of IL-6 (Fig. 2p). Inhibition of MLL1-Menin interaction inhibits the production of IL-6, leading to lower pSTAT3, Arp2/3, and protrusion generation. Together these results suggest that MLL1-Menin interaction controls 3D cell motility by regulating STAT3/Arp2/3-based actin filament assembly and associated protrusion generation via IL-6 production.

### MLL1-Menin interaction regulates TGF-β1-mediated myosin-II contractility

In addition to providing a translational aspect, the utilization of inhibitors also permits the modulation of the MLL1-Menin interaction at a much finer level that protein knockdowns. Increasing the concentration of MLL1-Menin inhibitors allows for a gradual ‘titration’ of cell migration (Figs. S2l and S3k). Given that the MLL1/IL-6/STAT3/Arp2/3-based actin filament assembly is perturbed by MLL1 inhibitors at low dosages, we next sought to determine if additional migration-based pathways were also controlled by the same MLL1-Menin interaction. Cells treated with a higher (30 µM) dose of MI-2-2 displayed the same extent of migration inhibition; however, supplementation of IL-6/8 did not rescue cell motility (Fig. 3a-b, dark green vs light green). Supplementation of IL-6/8 at three times the regular concentration (3x IL-6/8) also failed to rescue motility (Fig. S5f). Hence, a ‘deeper’ inhibition of the MLL1-Menin interaction affects cell migration by pathways other than IL-6/STAT3/Arp2/3-based actin filament assembly.

**Figure 3.**
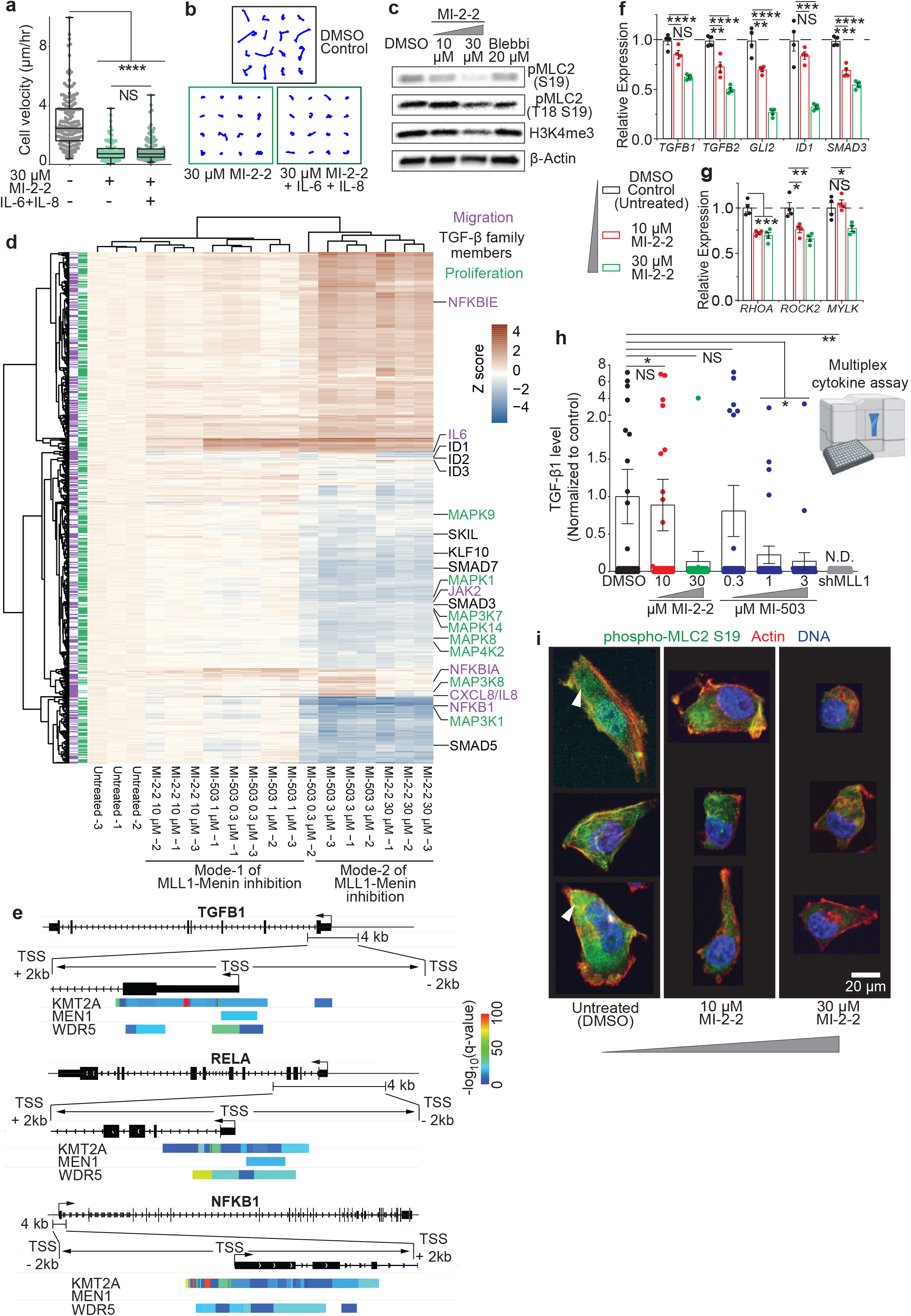
MLL1-Menin-based regulation of TGF-β1-mediated cell migration is mechanistically non-linear. **(a)** In contrast to 10 µM MI-2-2 treatment, supplementation of IL-6/8 did not rescue cell migration in 30 µM MI-2-2 treated cells, shown by cell velocities [n(cells) = 164(Untreat.), 85(MI), 121(MI+IL); one-way ANOVA (**** p < 0.0001, F (2, 367) = 106.1); two-tailed t-test (NS p(ML – MI+IL) = 0.6957, **** p(Untreat. - MI) < 0.0001, **** p(Untreat. – MI+IL) < 0.0001)] and **(b)** trajectories. **(c)** Phospho-MLC (both S19 and T18-S19) as well as H3K4me3 (right) were downregulated by MI-2-2 treatment. Blebbistatin treated cells were the positive control and actin was the loading control. **(d)** Heatmap of gene expression values for the genes involved in Hallmark cell migration (in purple) or proliferation (in green) gene sets shows two distinct gene expression patterns corresponding to low (mode-1) and high (mode-2) MLL1-Menin inhibitor dosage. TGF-β family members (labeled in black) were downregulated with deep MLL1-Menin inhibition. **(e)** ChIP-Atlas (an online ChIP-seq database)-based analysis reveals that MLL1, Menin, and WRD5 bind to the promoter region of TGFB1. COMPASS members also bound to NFKB1 and RELA promoter sequences. **(f)** qRT-PCR analysis shows that TGF-β pathway members are downregulated with MI-2-2. n(wells) = 4(each condition); one-way ANOVA (TGFB1: **** p < 0.0001, F (2, 9) = 30.35; TGFB2: **** p < 0.0001, F (2, 9) = 48.36; GLI2: **** p < 0.0001, F (2, 9) = 72.79; ID1: **** p < 0.0001, F (2, 9) = 31.00; SMAD3: **** p < 0.0001, F (2, 9) = 78.95). two-tailed t-test (TGFB1: NS p(Untreat. - 10) = 0.0506, **** p(Untreat. - 30) < 0.0001; TGFB2: ** p(Untreat. - 10) = 0.0039, **** p(Untreat. - 30) < 0.0001; GLI2: ** p(Untreat. - 10) = 0.0063, **** p(Untreat. - 30) < 0.0001; ID1: NS p(Untreat. - 10) = 0.2106, *** p(Untreat. - 30) < 0.0006; SMAD3: *** p(Untreat. - 10) = 0.0003, **** p(Untreat. - 30) < 0.0001). **(g)** Genes involved in regulating myosin contractility were also downregulated. n(wells) = 4(each condition); one-way ANOVA (RHOA: *** p = 0.0002, F (2, 9) = 26.95; ROCK2: *** p = 0.001, F (2, 9) = 16.47; MYLK: ** p = 0.0041, F (2, 9) = 10.79). two-tailed t-test (ROCK2: * p(Untreat. - 10) = 0.0131, ** p(Untreat. - 30) = 0.0022; MYLK: NS p(Untreat. - 10) = 0.4789, * p(Untreat. - 30) = 0.0157). **(h)** Multiplex cytokine analysis showed that TGF-β1 levels were unaffected with low dose MLL1 inhibition but were reduced with high dosage. N.D. - not detected. n(chambers) = 30(each condition); one-way ANOVA (** p = 0.0070, F (6, 203) = 3.054); two-tailed t-test (NS p(Untreat. - 10) = 0.8204, * p(Untreat. - 30) = 0.0289, NS p(Untreat. – 0.3) = 0.7004, * p(Untreat. – 1) = 0.0457, * p(Untreat. – 3) = 0.0269, ** p(Untreat. - sh) = 0.0076). **(i)** Immuno-fluorescence microscopy of MI-2-2 treated cells showed that MLL1-Menin inhibition reduces cell size, increases cell roundedness, and disrupts the actin cytoskeleton (red). 30 µM MI-2-2 treatment also reduces phospho-MLC2 (green), while 10 µM MI-2-2 treated cells still show pMLC2. Data in this figure was generated with MDA-MB-231 cells embedded in collagen gels except panels e (ChIP Atlas) and i (MDA-MB-231 cells in 2D).

Cell migration requires both actin filament assembly (regulated here by the IL-6/STAT3/Arp2/3 axis) and acto-myosin contractility (*74, 75*). Acto-myosin-based contractility is regulated via phosphorylation of myosin II (*74*), and levels of mono (S19) as well as di (T18 S19) phospho-myosin light chain II (pMLC2) were selectively reduced at a deep MLL1-Menin inhibition. pMLC2 (S19) levels showed a drastic reduction upon increasing inhibition of the MLL1-Menin inhibition (Fig. 3c, quantified in Fig. S5g), and was lower than that observed with Blebbistatin, a myosin II inhibitor. ROCK1 and ROCK2, major regulators of pMLC2, were also downregulated (Fig. S5g). Increasing depth of MLL1-Menin inhibition was indicated by a dose-dependent reduction in H3K4me3 levels, consistent with increasing extent of inhibition of MLL1’s histone methyltransferase activity. These results were also validated with another MLL1-menin inhibitor, MI-503 (Fig. S5h). Additionally, MLL1-depleted cells also exhibited reduced myosin contractility, indicated by reduced pMLC2 levels (Fig. S5i). Together, these results indicate that, in addition to actin filament assembly, MLL1-Menin interaction controls myosin-based contractility.

Hierarchical clustering on a heatmap of the most significantly downregulated migration and proliferation-related genes identified two distinct sets of gene expression (Fig. 3d), one corresponding to low doses of MLL-Menin inhibitors (10 µM MI-2-2, as well as 0.3 and 1 µM MI-503, hereby referred to as <u>Mode-1</u> of MLL1-Menin inhibition) and another corresponding to high doses (30 µM MI-2-2 and 3 µM MI-503, hereby referred to as <u>Mode-2</u> of MLL1-Menin inhibition) which forms a distinct pattern of gene expression corresponding to a deep inhibition of the MLL1-Menin interaction. Notably, several members of the TGF-β family (SMAD3, SMAD5, SMAD7, ID1, ID2, ID3, SKIL, and KLF10) were differentially downregulated in Mode-2 but were unchanged in Mode-1. To assess whether TGF-β based signaling was a potential mediator of MLL1-based myosin contractility, we examined whether MLL1 (KMT2A) directly bound to the promoter of *TGFB1* using ChIP-Atlas, a ChIP-seq database (*76*). Regions flanking the transcription start site (TSS), specifically ± 2 kb of the TSS, was also checked for Menin (MEN1) and the COMPASS member WDR5 (Fig. S2q). KMT2A bound directly to the TGFB1 promoter sequence (Fig. 3e) along with MEN1 and WDR5, indicating that COMPASS (and hence MLL1) directly binds to the TGFB1 promoter to potentially activate transcription. KMT2A and WDR5 also bound to the TGFB2 promoter (Fig. S5j), while MEN1 did not show binding to this region. Additionally, COMPASS members also bound to NFKB1 and RELA promoter sequences. Along with downregulation of NFKB1 in Fig. 2d and 3d, this hints at MLL1-based regulation of NF-κB signaling, which in turn controls IL-6 production (*77, 78*). Crucially, MLL1 depletion reduces levels of both these proteins, NF-κB1 (p50 and p105) as well as RELA (Fig. S5k-l). Consistent with NF-κB being upstream of IL-6, NF-κB inhibition in the absence of IL-6/8 supplementation did decrease motility (Fig. S5m). However, inhibition of NF-κB after IL-6/8-mediated rescue had no effect on cell motility.

At the transcript level, expression of both *TGFB1* and *TGFB2* were downregulated following MLL1- Menin inhibition (Fig. 3f, heatmaps in fig. S5n). However, only *TGFB1* showed a non-linear response with MLL1-Menin inhibition, mirroring the trend observed with myosin contractility. Downstream TGF-β family members were also downregulated by MLL1 inhibition. GSEA enrichment plots show downregulation of key cell migration-related pathways for mode-2 MLL1 inhibition cells, including TGF- β signaling and epithelial-mesenchymal transition (Fig. S5o). Additionally, levels of TGF-β receptors*, TGFBR1* and *TGFBR2,* were unaffected by MLL1-menin inhibition (Fig. S5p), indicating that the downregulation of TGF-β signaling was potentially due to down regulation of secreted products. Thus, TGF-β1 (gene product of *TGFB1*) was hypothesized to be the likely regulator of pMLC2, that lay downstream of MLL1-Menin interaction. Genes related to myosin contractility such as *RHOA*, *ROCK2*, and *MYLK* were also downregulated with MLL1-Menin inhibition (Fig. 3g). Myosin light chain kinase (*MYLK*), essential for maintaining the phosphorylation of MLC2 (*79, 80*), was downregulated selectively at mode-2 but not mode-1; consistent with reduced pMLC2 levels observed exclusively with mode-2 inhibition in Fig. 3c. Similar results were also seen in shMLL1 cells (Fig. S5q) and BT549 TNBC cells (Fig. S6a). Secretion of TGF-β1, measured via a multiplex cytokine assay, was decreased by mode-2 MLL1-Menin inhibition, but were unchanged with mode-1 inhibition (Fig. 3h). shMLL1 cells, which were hypothesized to phenocopy the mode-2, or deep, MLL1 inhibition, also displayed a dramatic reduction in TGF-β1 levels; with TGF-β1 secretion not detected in shMLL1 cells. In contrast, TGF-β2 levels were largely unchanged, ever at deep MLL1 inhibition (Fig. S6b). MLL1-Menin inhibition also impacted cell morphology in addition to cell migration machinery. Untreated cells were elongated, had high levels of pMLC2 (green, pointed to by arrowheads), and featured prominent actin fibers (red, Fig. 3i, left). Mode- 2 MLL1 inhibition led to more rounded cells with virtually no pMLC2 staining (Fig. 3i, right). Mode-1 cells looked similar to mode-2 cells, but had higher levels of pMLC2 (Fig. 3i, middle). MLL1-Menin inhibition also increased cell roundedness as assessed by our machine learning algorithms (Fig. S6c).

To summarize, a deeper MLL1-Menin inhibition also reduces myosin contractility, and its effect on 3D cell migration cannot be reversed solely by IL-6/8 supplementation. Concurrently, secretion of another cytokine, TGF-β1, and the expression of TGF-β family members were also downregulated at higher dosages of MLL1-Menin inhibitor.

### TGF-β1 and IL-6/8 are necessary and sufficient for MLL1-Menin based cell migration

In addition to actin filament assembly, MLL1 also regulates myosin contractility. As a result, supplementation of exogenous IL-6 was no longer sufficient to rescue cell motility. To assess whether the reduction in motility in mode-2 was also associated with loss of TGF-β1, exogenous TGF- β1 was added to MLL1-Menin inhibited cells (Fig. S6d). Supplementation of either IL-6/8 (dark green) or TGF- β1 (brown) by itself to did not change motility (Velocities in Fig. 4a and trajectories in Fig. 4b). However, concurrent supplementation of both IL-6/8 and TGF-β1 (blue) restored cell motility to that observed with untreated control (black). This is consistent with the fact that both actin filament assembly via IL-6/8 and myosin contractility via TGF-β1 are required for cell motility. MLL1-depleted (shMLL1) cells had >90% of MLL1 depleted (Fig. S2d) and was hypothesized to behave similarly to mode-2 MLL1 inhibition, including rescue of motility via supplementation of IL-6/8 + TGF-β1. Notably, shMLL1 cells show decreased levels of IL-6 (Fig. S4p), IL-8 (Fig. S4q), and TGF-β1 (Fig. 3h). Supplementation of IL- 6/8 + TGF-β1 to shMLL1 cells (blue) restored their motility to levels observed with scrambled control cells (black, Fig. 4c). Once again, supplementation of either IL-6/8 or TGF-β1 alone did not increase motility to levels observed with scrambled control. These rescue experiments show that at mode-2 MLL1-Menin inhibition, TGF-β1 production is also decreased along with IL-6/8, and supplementation of these cytokines can fully rescue 3D cell migration.

**Figure 4.**
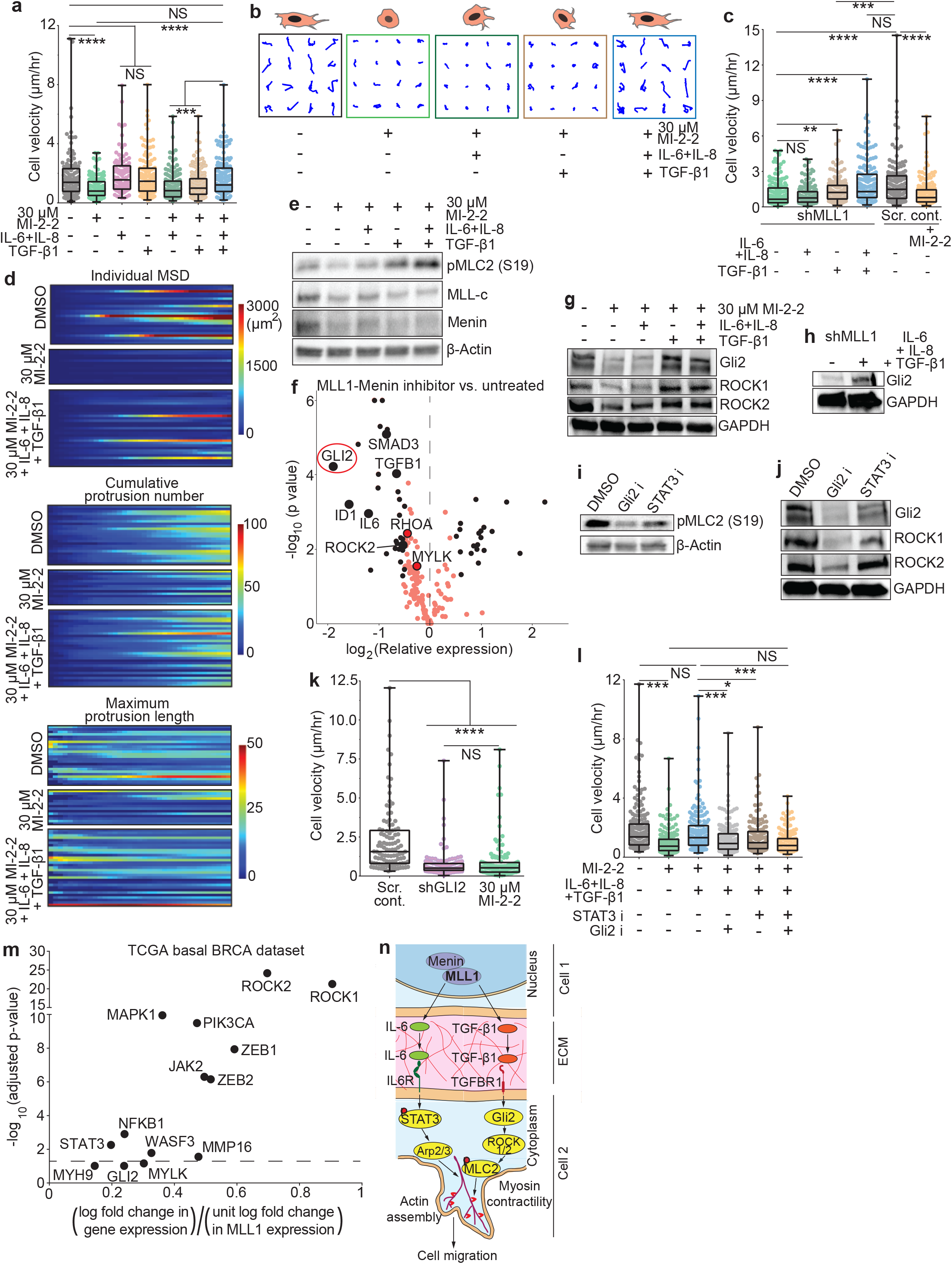
MLL1-Menin interaction regulates myosin-based contractility via a Gli2/ROCK1/2/pMLC2 axis. **(a)** Supplementation of TGF-β1 along with IL-6/8 rescued cell migration to levels observed with untreated control. Supplementation of either TGF-β1 or IL-6/8 on its own did not rescue motility. n(cells) = 119(Untreat.), 122(MI), 96(IL), 123(TGF), 162(MI+IL), 175(MI+TGF), 154(MI+IL+TGF); one-way ANOVA (**** p < 0.0001, F (6, 944) = 13.35); two-tailed t-test (**** p(MI – MI+IL+TGF) < 0.0001, NS p(Untreat. – MI+IL+TGF) = 0.4581, **** p(Untreat. - MI) < 0.0001, NS p(Untreat. - IL) = 0.5982, NS p(Untreat. - TGF) = 0.8383, **** p(MI+IL – MI+IL+TGF) < 0.0001, *** p(MI+TGF – MI+IL+TGF) = 0.0004). **(b)** TGF-β1+IL-6/8-mediated rescue of cell motility is evident in individual cell trajectories. **(c)** Supplementation of TGF-β1+IL-6/8 to MLL1-depleted cells increased their motility to levels seen with control cells. n(cells) = 172(sh), 164(sh+IL), 158(sh+TGF), 162(sh+IL+TGF), 149(SC), 178(SC+MI); one-way ANOVA (**** p < 0.0001, F (5, 977) = 19.54); two-tailed t-test (**** p(sh – sh+IL+TGF) < 0.0001, NS p(sh+IL+TGF - SC) = 0.9070, NS p(sh – sh+IL) = 0.2482, ** p(sh – sh+TGF) = 0.0073, *** p(sh+TGF - SC) = 0.0005, **** p(sh - SC) < 0.0001, **** p(SC- SC+MI) < 0.0001, NS p(sh – SC+MI) = 0.5494). **(d)** IL-6/8+TGF-β1 rescued protrusion generation in MLL1-inhibited cells. Temporal heatmap of cumulative protrusion number and maximum protrusion length show a rescue concurrent with an increase in MSD. **(e)** Levels of pMLC2 were rescued with supplementation of TGF-β1 (either by itself or concurrently with IL-6/8), indicating restoration of myosin contractility. **(f)** A volcano plot of genes assessed by PCR shows *GLI2*, a member of the TGF-β signaling pathway, as the most downregulated gene after MLL1-Menin inhibition. **(g)** Gli2, ROCK1, and ROCK2 levels were rescued by replenishment of TGF-β1, indicating that these lie downstream of both the MLL1-Menin interaction and TGF-β1. **(h)** shMLL1 cells expressed a low level of Gli2, which was rescued by supplementation of TGF-β1+IL-6/8. **(i)** Inhibition of Gli2 (using GANT-61), but not STAT3, reduced pMLC2, indicating that myosin contractility was largely affected via TGF-β1-Gli2 signaling, rather than IL-6-STAT3 signaling. **(j)** Gli2 inhibition also reduced levels of ROCK1 and ROCK2, necessary for myosin contractility, indicating that TGF-β1 and Gli2 regulated myosin contractility was mediated by ROCK1/2. **(k)** Gli2 knockdown reduces cell migration. n(cells) = 106(Scr. cont.), 126(shGLI2), 122(MI-2-2); one-way ANOVA (**** p < 0.0001, F (2, 351) = 35.65); two-tailed t-test (**** p(SC – shGLI2) < 0.0001, NS p(shGLI2 – MI-2-2) = 0.2295, **** p(SC – MI-2-2) < 0.0001). **(l)** Motility rescued by TGF-β1+IL-6/8 supplementation after MI-2-2 treatment (blue) is lost by inhibiting either Gli2 (downstream of TGF-β1, grey) or STAT3 (downstream of IL-6/8, brown). Treatment with both Gli2 and STAT3 inhibitors led to the lowest cell motility (orange), comparable to MLL1 inhibited cells (green). n(cells) = 199(Untreat.), 189(MI), 172(MI+IL+TGF), 182(MI+IL+TGF+Gli2i), 145(MI+IL+TGF+STAT3i), 159(MI+IL+TGF+Gli2i+STAT3i); one-way ANOVA (**** p < 0.0001, F (5, 1040) = 17.35); two-tailed t-test (NS p(Untreat. – MI+IL+TGF) = 0.5535, **** p(Untreat. - MI) < 0.0001, *** p(MI+IL+TGF - MI+IL+TGF+Gli2i) = 0.0002, * p(MI+IL+TGF - MI+IL+TGF+STAT3i) = 0.0169, **** p(MI+IL+TGF – MI+IL+TGF+Gli2i+STAT3i) < 0.0001, NS p(MI – MI+IL+TGF+Gli2i+STAT3i) = 0.8970). **(m)** The expression of several key genes implicated in MLL1-Menin regulated cell migration and proliferation are positively correlated with MLL1 expression in TCGA basal breast cancer dataset. ROCK1 and ROCK2 were the two most closely correlated genes. **(n)** Deep (mode-2) inhibition of MLL1-Menin interaction disrupts motility in a two-pronged manner. MLL1-Menin interaction controls the production of IL-6, which regulates motility via STAT3-Arp2/3 mediated protrusion generation. Additionally, MLL1- Menin interaction also regulates the production of TGF-β1, which controls myosin contractility via a Gli2-ROCK1/2-pMLC2 axis. Data in this figure was generated with MDA-MB-231 cells embedded in 3D collagen gels except panel m (TCGA).

The rescue of cell migration via cytokine supplementation was accompanied by an increase in actin filament assembly and myosin contractility. Protrusion analysis showed that MLL1-Menin inhibition reduced the number of protrusions and maximum protrusion length (Fig. 4d). Supplementation of IL- 6/8 + TGF-β1 rescued these parameters, with rescued cells generating similar sized and number of protrusions as untreated control. Along with protrusion generation (actin filament assembly), myosin contractility was also restored in rescued cells (Fig. 4e, quantified in Fig. S6e), with supplementation of TGF-β1 (by itself or as IL-6/8 + TGF-β1) increasing pMLC2 levels. Supplementation with IL-6/8 did not fully rescue pMLC2, demonstrating that pMLC2 lost by MLL1-menin inhibition can be regained by supplementing cells with TGF-β1. None of the supplementations affected the levels of MLL1 or Menin appreciably, consistent with the fact that both MLL1 and Menin lie upstream of these cytokines.

Thus, MLL1-Menin interaction controls 3D cell migration via the secretion of cytokines IL-6 and TGF- β1. Supplementation of exogenous IL-6/8 and TGF- β1 rescued actin filament assembly and myosin contractility, respectively, thus restoring cell motility.

### TGF-β1-based myosin contractility is mediated by a Gli2-ROCK1/2-pMLC2 axis

MLL1 regulates cell motility in part by controlling the secretion of TGF-β1. To identify the mediators of TGF-β1-mediated myosin contractility, we conducted a transcript level (PCR-based) assessment of several genes following mode-2 MLL1-Menin inhibition, which revealed *GLI2* to be the most downregulated gene (Fig. 4f, circled in red). Several members of the TGF-β family (*TGFB1*, *ID1*, *SMAD3*, *SMAD7*) as well as genes involved in myosin contractility (*RHOA*, *ROCK2*, *MYLK*) were also downregulated. Additionally, *GLI2* was the most differentially expressed gene between mode-2 *vs*. mode-1 inhibition of MLL1-Menin interaction (Fig. S6f). *GLI2* has been shown to play a role in metastasis in melanoma and breast cancers (*81, 82*), but not in cell migration itself. Critically, *GLI2* has been reported to lie downstream of both TGF-β and Hedgehog signaling (*81*). As MLL1-Menin inhibition does not induce an appreciable change in Hedgehog signaling, including in the expression levels of ligands, receptors, and intermediates (Fig. S6g-h), we reasoned that Gli2 mediates the TGF-β1-based regulation of myosin contractility.

Gli2 expression was downregulated by MLL1-Menin inhibition (Fig. 4g). Supplementation of IL-6/8 did not restore Gli2 expression, but supplementation of TGF-β1 (or IL-6/8 + TGF-β1) restored Gli2 levels, indicating that Gli2 lies downstream of TGF-β1. MLL1-depleted (shMLL1) cells also expressed low levels of Gli2, which was increased after supplementation with IL-6/8 + TGF-β1 (Fig. 4h). Treatment of cancer cells with Gli2 inhibitor (Gli2 i) drastically reduced pMLC2 levels (Fig. 4i), indicating that Gli2 is a major regulator of myosin contractility. STAT3 inhibition did not change pMLC2 levels appreciably, denoting little crosstalk between STAT3-mediated actin filament assembly and myosin contractility. Along with Gli2, levels of ROCK1 and ROCK2 were also reduced by MLL1-Menin inhibition and restored with TGF-β1 supplementation (Fig. 4g). ROCK1 and ROCK2, are serine/threonine kinases that are responsible for phosphorylating myosin light chain and are essential for maintaining pMLC2 levels (*83–85*). Direct inhibition of Gli2 also reduced levels of both ROCK1 and ROCK2 (Fig. 4j), indicating that Gli2-mediated myosin contractility was mediated by ROCK1/2. In contrast, STAT3 inhibition had little effect on ROCK1/2, which is consistent with STAT3 being a major regulator of actin filament assembly, but not myosin contractility. Thus, TGF-β1-mediated myosin contractility is regulated by the Gli2- ROCK1/2-pMLC2 axis. To verify that Gli2 is essential for cell migration, Gli2 was depleted via shRNA (Fig. S6i). shGli2 cells were non-motile and exhibited velocities similar to MLL1-Menin inhibited cells (Fig. 4k). Further, a Gli2 inhibitor (Gli2 i) was used to determine if motility rescued by cytokine supplementation could be lost by Gli2 inhibition (Fig. 4l). Mode-2 inhibited cells showed reduced motility (green compared black), which was rescued after supplementation with IL-6/8 + TGF-β1 (blue). Inhibition of STAT3 (brown) or Gli2 (grey) on top of the MLL1 inhibition and cytokine supplementation reduced cell motility again, indicating that both proteins play an important role in cell migration downstream of cytokines. Maximum inhibition of cell migration was obtained by concurrent STAT3 and Gli2 inhibition (orange, nullifying the effect of supplemented IL-6/8 and TGF-β1), which also reduced the distribution of cell velocities to that observed with MLL1 inhibition (green).

To further substantiate our elucidated mechanism of MLL1-based cell migration, basal breast cancer RNA-seq data from PanCanAtlas (TCGA) was examined to identify genes and gene sets that were significantly correlated with MLL1 expression (Fig. 4m and S6j). Top genes correlating with MLL1 expression included ROCK1 (0.9 log fold change in gene per log fold increase in MLL1 expression) and ROCK2 (0.7 log fold change) (Fig. 4m). Other genes that were implicated in MLL1-Menin-mediated cell migration and found to positively correlate with MLL1 expression included JAK2, STAT3, GLI2 and MYLK (although GLI2 and MYLK were just under the significance threshold). Epithelial-mesenchymal transition genes ZEB1 and ZEB2 were also significantly and positively correlated with MLL1 expression. Additionally, focal adhesion, ECM receptor interaction, myosin phosphorylation, and epithelial-mesenchymal transition gene sets were all significantly positively correlated with MLL1 expression in TCGA basal breast cancer dataset (Figs. S6j and S7a).

Thus, our results establish that TGF-β1 lies downstream of MLL1-Menin interaction along with IL-6 and is depleted exclusively in mode-2 MLL1-menin inhibition (such as by 30 µM MI-2-2, 3 µM MI-503, or in shMLL1 cells). Gli2 lies downstream of TGF-β1, while STAT3 lies downstream of IL-6 (Fig. 4n). Deep inhibition of the MLL1-Menin interaction inhibits the secretion of both IL-6 and TGF-β1, leading to lower pSTAT3 and Gli2 levels. Lower pSTAT3 levels lead to lesser actin filament assembly via reduced nucleator Arp3, while lower Gli2 levels correspond to decreased myosin contractility via ROCK1/2 and pMLC2. Therefore, MLL1-Menin interaction controls 3D cell motility by regulating both actin-assembly via the IL-6-STAT3-Arp3 axis and myosin contractility via the TGF-β1-Gli2-ROCK1/2-pMLC2 axis.

### MLL1-Menin inhibition reduces cell proliferation without altering cell cycle distribution

In addition to controlling cell migration, MLL1 also regulates cell proliferation. Cells depleted of MLL1 (Fig. 1f) or treated with MLL1-Menin inhibitors (MI-2-2 and MI-503: Fig. 5a, VTP50469: Fig. S7b) showed reduced cell numbers. Unlike inhibition of cell migration, which was dose-independent, quantification of cell numbers in 3D collagen matrices showed that MLL1-Menin inhibition reduces cell proliferation in a dose-dependent manner (Fig. 5a). Kinetics of cell proliferation arrest revealed that the effect of MLL1 inhibition on proliferation was evident after day two. Cell proliferation was inhibited in mode-2 leading to a near growth-arrest, while mode-1 had minimal impact (Fig. S7c-d). Proliferation affected by MLL1-Menin inhibition was also readily reversible. Cells were pre-treated with MLL1 inhibitor for two days before being seeded and subjected to further drug treatment. Pre-treated cells without further treatment showed increased cell proliferation two days after drug washout (Fig. S7e). Both pre-treated cells and non-pretreated cells receiving further drug treatment showed the lowest cell numbers. In addition to TNBC cells, these results were also validated in fibrosarcoma cells (Fig. S7f). MLL1-Menin inhibition reduced transcript levels of key markers of cellular proliferation, *KI67* and *PCNA* (Fig. 5b, heatmaps in Fig. S7g). Expression of both genes were halved following MLL1-menin inhibition. Surprisingly, MLL1-Menin inhibition did not alter the cycle distribution appreciably, and MLL1-Menin inhibited cells did not show appreciable accumulation in the G0/G1 phase (Fig. S7h). Cell-cycle distribution after mode-1 inhibition was identical to those for untreated control. Mode-2 inhibition led to a slight accumulation in G0/G1 phase and a corresponding decrease in S phase, but not enough to justify the observed arrest in proliferation. Another TNBC cell line (BT-549) showed decreased cell numbers with minimal change in apoptosis levels (Fig. S7i-j), a largely unchanged cell cycle distribution (Fig. S7k), and decrease in *KI67* levels (Fig. S6a). Thus, MLL1-Menin interaction controls cell proliferation, however, inhibition of this interaction does not change cell cycle distribution.

**Figure 5.**
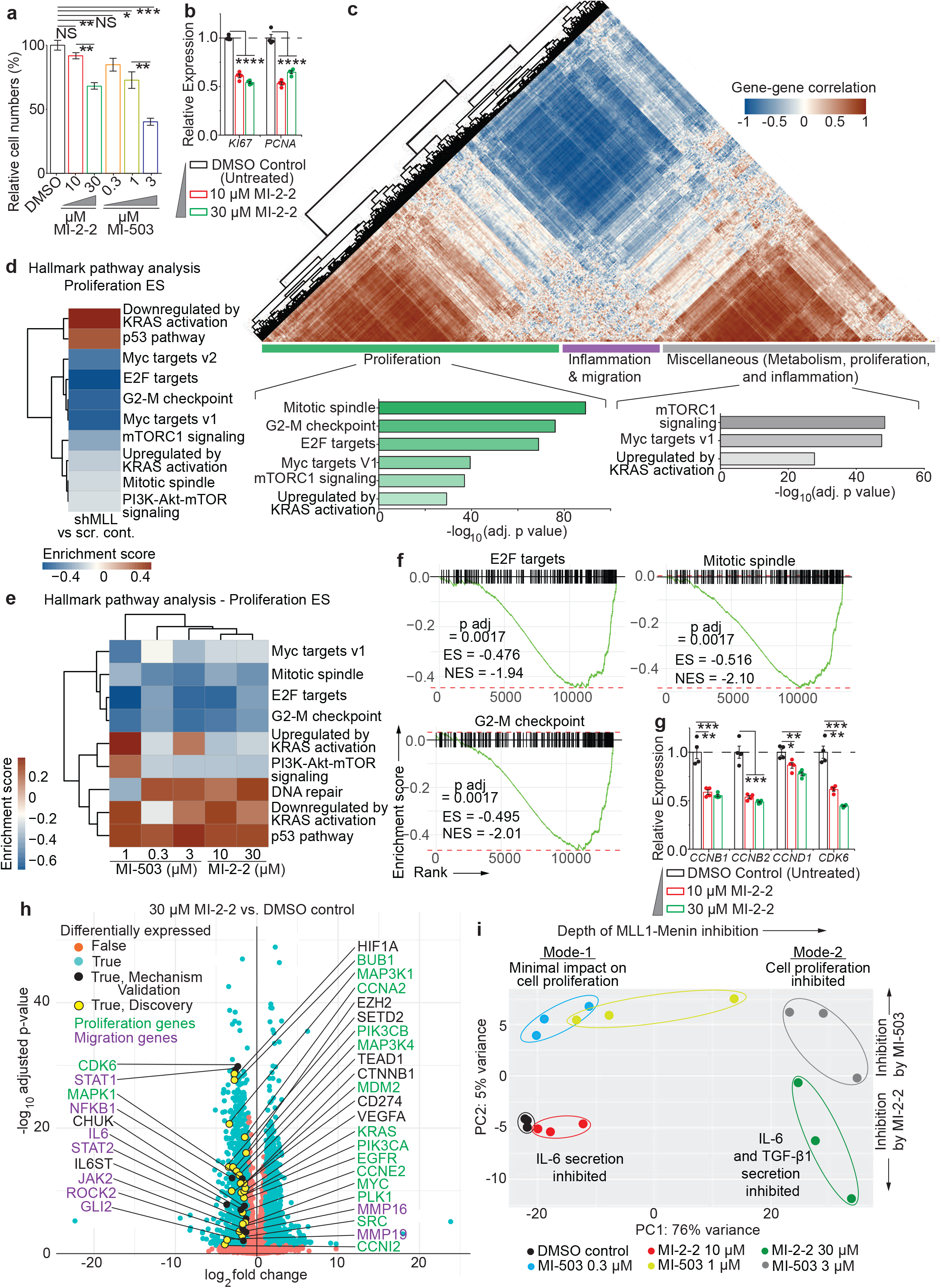
MLL1-Menin regulates cell proliferation via a multitude of proliferation and cell-cycle-related pathways. **(a)** MLL1-Menin inhibition with MI-2-2 or MI-503 results in a dose-dependent growth suppression. n(wells) = 4(each condition); one-way ANOVA (**** p < 0.0001, F (6, 21) = 39.05); two-tailed t-test (NS p(Untreat. - 10) = 0.1299, ** p(Untreat. - 30) = 0.0023, ** p(10 - 30) = 0.0026, NS p(Untreat. – 0.3) = 0.0735, * p(Untreat. - 1) = 0.0223, *** p(Untreat. - 3) = 0.0002, ** p(1 - 3) = 0.0036). **(b)** MLL1-Menin inhibition decreases RNA-level expression of markers of proliferation, *KI67* and *PCNA* (qRT-PCR). n(wells) = 4(each condition); one-way ANOVA (KI67: **** p < 0.0001, F (2, 9) = 279.8; PCNA: **** p < 0.0001, F (2, 9) = 95.95). **(c)** Gene-gene correlation analysis on the 3926 Hallmark gene set genes that were expressed revealed three distinct modules: proliferation, inflammation and migration, and a mixed module. Key cell-cycle and proliferation genes were significantly enriched in modules 1 and 3. RNA-seq analysis of **(d)** shMLL1 cells vs. scrambled control cells or of **(e)** MLL1-Menin inhibited cells shows downregulation of several key proliferation and cell-cycle related pathways as well as upregulation of anti-proliferative pathways. **(f)** GSEA enrichment plots of cell-cycle-related pathways for mode-2 MLL1 inhibited cells (30 µM MI-2-2 treatment). **(g)** Transcription of key cell cycle genes such as *CCND1, CDK6, CCNB1,* and *CCNB2* were reduced with MLL1 inhibition assessed via qRT-PCR. n(wells) = 4(each condition); one-way ANOVA (CCNB1: **** p < 0.0001, F (2, 9) = 36.77; CCNB2: **** p < 0.0001, F (2, 9) = 56.68; CCND1: *** p = 0.0009, F (2, 9) = 16.64; CDK6: **** p < 0.0001, F (2, 9) = 55.58). two-tailed t-test (CCNB1: ** p(Untreat. - 10) = 0.0011, *** p(Untreat. - 30) = 0.0005; CCNB2: *** p(Untreat. - 10) = 0.0004, *** p(Untreat. - 30) = 0.0002; CCND1: * p(Untreat. - 10) = 0.0214, ** p(Untreat. - 30) = 0.0011; CDK6: ** p(Untreat. - 10) = 0.0011, *** p(Untreat. - 30) = 0.0001). **(h)** Volcano plot of mode-2 (30 µM MI-2-2) vs. untreated (DMSO) control show nearly 4000 genes that are significantly impacted by MLL1-Menin inhibition (log2 fold change > 1 and p-value < 0.05, plotted in blue). These genes include those that have been implicated in our MLL1-Menin based regulation of cell migration and proliferation (validation, in black). Additionally, a subset of these significant genes consists of proliferation-related genes that further extend the scope of MLL1-Menin interaction in regulation of proliferation (discovery, in yellow). **(i)** PCA analysis on all conditions show that lower doses of MLL1-Menin inhibitors (mode-1) cluster closer to the untreated control than to higher drug dosages. Each oval encompasses all technical replicates for that condition. All data in this figure was generated with MDA-MB-231 cells embedded in 3D collagen gels.

To assess the impact of MLL-Menin interaction on proliferation, we performed correlation analysis on our RNA-seq data for the 3926 genes that are a part of Hallmark gene sets (MSigDB collection) and that were expressed in our samples. Proliferation genes formed the largest module (Fig. 5c), consisting mainly of mitosis and cell cycle-based gene sets. Gene sets pertaining to inflammation and cell-migration formed the second module, while the third module was a mixture of all remaining gene sets including oncogene-based proliferation gene sets. Enrichment analysis on all proliferation-related pathways for shMLL1 and MLL1-Menin inhibited cells emphasized the downregulation of key cell cycle-based pathways including E2F targets (impacting G1/S transition), G2M checkpoint, and mitotic spindle assembly (Fig. 5d-e, GSEA enrichment plots in Fig. 5f and S8a). Key proliferation pathways such as Myc targets, PI3K-Akt-Mtor signaling, and (genes upregulated by) KRAS signaling were also diminished by MLL1 inhibition. Additionally, anti-proliferative pathways such as p53 pathway and DNA repair, which induce proliferative arrest were upregulated. The extent of downregulation in these pathways was dose-dependent, but unlike migration, the identities of pathways affected were dose-independent. MLL1-Menin inhibition affected a wide variety of cell cycle regulators at the transcript level, including cyclins and CDKs (Fig. 5g). *CCND1* and *CDK6*, regulators of G1/S transition (*86*), as well as *CCNB1* and *CCNB2*, major regulators of G2/M transition (*86*), were downregulated by MLL1- Menin inhibition. Downregulation of pathways and key genes involved in both G1/S and G2/M transitions is consistent with decreased proliferation accompanied with minimal change in cell cycle distribution following MLL1-Menin inhibition. Members of the MAPK family (*MAPK1, MAPK10,* and *MAPK12)*, which mediate cell proliferation (*87*), were also downregulated with MLL1-Menin inhibition (Fig. 3d and S8b). MAPK1 was also found to be positively correlated with MLL1 in our TCGA analysis (Fig. 4m) along with G2/M checkpoint gene set (Figs. S6j and S7a). The same cell cycle factors were impacted by MLL1 depletion in shMLL1 cells (Fig. S8c). Thus, MLL1-Menin interaction controls key proliferation-related genes and pathways, including markers of proliferation, cell cycle genes, and members of the MAPK family.

Transcriptomic analysis revealed that 3978 genes were significantly down (1994) or up (1984)- regulated in mode-2 MLL1 inhibition. These genes corroborated the abovementioned mechanism of MLL1-based cell migration (Validation, key genes labelled in black on the left side) and revealed further proliferation genes and transcription factors that were regulated by the MLL1-Menin interaction (Discovery, labelled in yellow on the right side). Genes previously implicated in MLL1-menin based cell migration including IL6, JAK2, STAT1, STAT2, NFKB1, GLI2, and ROCK2 were all downregulated by MLL1-Menin inhibition (Fig. 5h). Additionally, other genes shown to play a role in cell migration (HIF1A, MMP16, and MMP19), proliferation (KRAS, EGFR, SRC, PIK3CA, PIK3CB, CTNNB1, MDM2, MAP3K1, and MAP3K4), cell cycle and mitosis (CCNA2, CCNE2, CCNI2, BUB1, and PLK1), transcription and epigenetic factors (TEAD1, EZH2, MYC, and SETD2), and other genes necessary for tumor progression (CD274 and VEGFA) were also downregulated by MLL1-Menin inhibition. This hints at a compelling role for MLL1 in regulating cancer cell proliferation and migration, as well a broader and multifaceted role in tumor progression including those in angiogenesis and immune evasion. Clustering analysis showed that mode-1 MLL1-Menin inhibition (10 µM MI-2-2 and 0.3 µM MI-503) clustered closer to untreated control than to mode-2 MLL1-Menin inhibition (30 µM MI-2-2 and 3 µM MI-503) (Fig. 5i). PC1 accounted for the ‘depth’ of MLL1-Menin inhibition (76% variance), while PC2 accounted for the differences between the two drugs, MI-2-2 and MI-503 (5% variance). In addition to migration and proliferation pathways, various metabolic and immune-related pathways were also affected by MLL1- menin inhibition (Fig. S8d-e). Further analysis of these pathways is beyond the scope of the manuscript. Analysis of MLL1-Menin inhibition by MI-503 showed the same targets and underlying mechanism (Fig. S8f).

Any crosstalk between cell migration and proliferation-based pathways was determined by assessing if a rescue in cell migration impacted proliferation or cell cycle distribution. Addition of IL-6/8 to MLL1- Menin inhibited cells did not alter their proliferation (Fig. S8g) or their cell cycle distribution (Fig. S8h, blue vs. purple bars). Additionally, supplementation with IL-6/8 and/or TGF-β1 did not increase levels of key proliferation genes, *KI67* and *PCNA,* or cell cycle genes (Fig. S8i), indicating that supplementation of cytokines can rescue cell motility, but not proliferation. Thus, the regulation of cell migration and proliferation by the MLL1-Menin interaction occurs via independent pathways. Hence, the MLL1-Menin interaction controls TNBC cell proliferation via a multitude of cell-cycle related and proliferation-based pathways. Further, this regulation of cell proliferation is uncoupled from its regulation of cell migration.

### MLL1 depletion reduces metastatic burden in vivo

As MLL1 plays a critical role in cell migration, depletion of MLL1 has a negative impact on the metastasis of cancer cells. Mice bearing MLL1-deficient tumors exhibited longer survival and lower metastatic burden (Fig. 1g-k). However, MLL1-deficient tumors grew more slowly, and reduced tumor growth rate may obscure the contribution of MLL1-regulated cell-migration on metastatic burden. To account for this, mice bearing shMLL1, scrambled control and wild-type tumors were sacrificed when the tumors reached a set threshold size (1400 mm^3^) (Fig. 6a-c). shMLL1 tumors grew for ∼50% longer time (62 days post tumor establishment) than scrambled control or wild-type tumors (41 and 43 days, respectively). Lungs from mice bearing scrambled control and wild type tumors showed more numerous and larger metastatic foci (evident as darker staining) than lungs from shMLL1 tumor bearing mice (Fig. 6d and S9a). Quantification of total lung metastatic burden showed three times lower lung metastatic burden for shMLL1 tumor bearing mice compared to mice bearing scrambled control tumors (Fig. 6e). Histogram of metastatic lesions showed that MLL1 depletion not only led to fewer metastatic lesions, but these lesions were also much smaller compared to WT or scrambled control cells (Fig. 6f). Fewer number of lesions (Fig. 6f inset) may be the result of lower cell migration or decreased survivability of cells in circulation. Given that shMLL1 mice were sacrificed ∼3 weeks after control mice, metastasized cells that have reached the lungs would have a longer time to proliferate and colonize. Hence, smaller metastases may not be due to lower proliferation of shMLL1 cells at the metastatic site. Thus, MLL1 depletion inhibits metastasis independent of its effect on proliferation at the primary tumor site.

**Figure 6.**
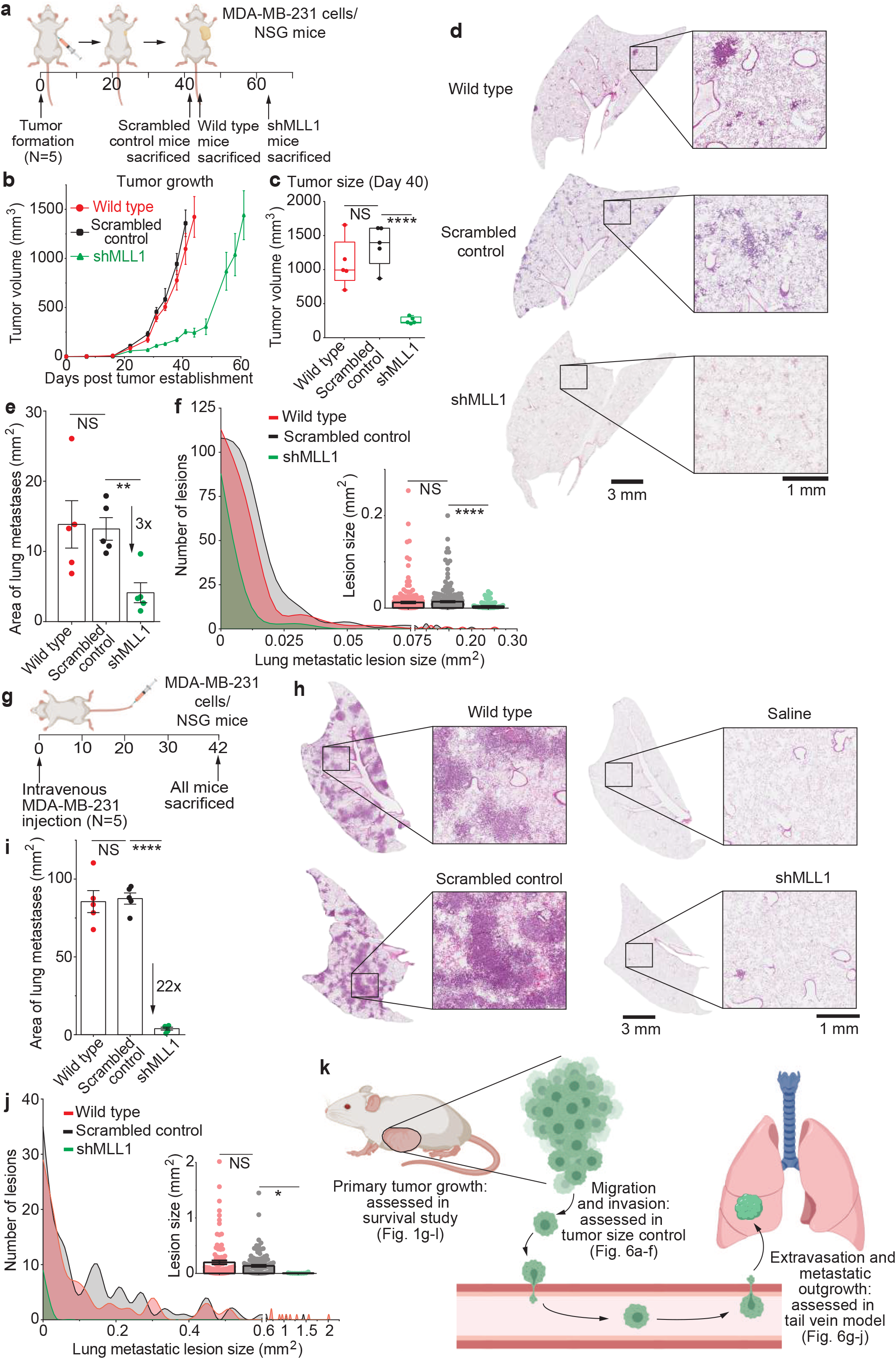
MLL1-depleted cancer cells exhibit decreased lung metastases *in vivo*. **(a)** Orthotopic breast tumors were established in NSG mice by injecting cells into mammary fat pad. Mice were sacrificed upon reaching a set threshold size (1400 mm^3^). shMLL1 mice were sacrificed ∼3 weeks after control mice. **(b)** Wild type and scrambled control tumors grew faster than shMLL1 tumors. **(c)** This difference in growth rate is also illustrated in tumor sizes at day 40. n(mice) = 5(each condition); one-way ANOVA (**** p < 0.0001, F (2, 12) = 22.91); two-tailed t-test (**** p(SC-sh) < 0.0001, NS p(SC- WT) = 0.2440). **(d)** shMLL1-tumor bearing mice showed reduced lung metastatic burden despite being sacrificed later and at the same primary tumor size as scrambled control and wild type tumors. **(e)** Quantification of total metastatic burden per lung shows a three-fold reduction in shMLL1 lungs compared to scrambled control. n(lungs) = 5(each condition, 1 lung/mouse); one-way ANOVA (* p = 0.0210, F (2, 12) = 5.e19); two-tailed t-test (** p(SC - sh) = 0.0029, * p(WT - sh) = 0.0313, NS p(WT - SC) = 0.8960). **(f)** Histogram of metastatic lesions of representative lungs show that the reduced metastatic burden in shMLL1 lungs was due to fewer metastatic lesions that were also smaller. n(lesions) = 227(WT), 270(SC), 115(sh); one-way ANOVA (** p = 0.0010, F (2, 609) = 6.984); two-tailed t-test (**** p(SC - sh) < 0.0001, NS p(WT - SC) = 0.4636). **(g)** Extravasation and metastatic outgrowth were assessed using a non-spontaneous metastatic model that was performed via tail-vein injection of wild type, scrambled control, or shMLL1 cells. Mice were sacrificed 6 weeks after injection. **(h)** Mice injected with control cells showed numerous and large metastatic lesions while shMLL1 lungs showed barely any lesions. Wild type cells and saline were the corresponding controls. **(i)** Quantification of total metastatic burden per lung shows more than a 20-fold decrease for shMLL1 lungs compared to scrambled control. n(lungs) = 5(each condition, 1 lung/mouse); one-way ANOVA (**** p < 0.0001, F (2, 12) = 107.0); two-tailed t-test (**** p(SC - sh) < 0.0001, **** p(WT - SC) < 0.0001, NS p(WT - SC) = 0.8107). **(j)** Histogram of metastatic lesions for the representative lungs shown in Fig. 6d show a dramatically reduced lesion size distribution for shMLL1 lungs compared to scrambled control or wild type. n(lesions) = 101(WT), 122(SC), 10(sh); one-way ANOVA (* p = 0.0487, F (2, 230) = 3.063); two-tailed t-test (* p(SC - sh) = 0.0444, NS p(WT - SC) = 0.0961). **(k)** Representation of the role of MLL1 in various steps of the metastatic cascade. Three different *in vivo* studies have been utilized to elucidate the contribution of MLL1-Menin interaction in driving primary tumor growth (left), cancer cell migration and invasion (center), as well as extravasation and metastatic outgrowth (right). All data in this figure was generated with MDA-MB-231 cells in NSG mice.

Next, the colonization and metastatic outgrowth of MLL1-depleted cells was assessed using a tail-vein metastasis model (Fig. 6g). Lungs from mice injected with either scrambled control or wild type cells displayed extensive metastasis with large colonies (Fig. 6h and S9c, sacrificed six weeks after injection). Lungs from mice injected with shMLL1 cells were mostly clear and looked similar to saline-injected lungs. Quantification of lung metastatic burden showed a 22-fold decrease in lung metastatic burden for mice bearing MLL1-depleted tumors compared to scrambled control or wild type control (Fig. 6i). shMLL1-lungs showed dramatically fewer and smaller metastatic colonies (Fig. 6j), indicating that MLL1-depletion plays a role in extravasation and metastatic outgrowth of lesions. Thus, MLL1 depletion reduces metastatic outgrowth of cancer cells and may also have an impact on extravasation of cancer cells. Hence, MLL1 regulates metastasis of TNBC cells in multiple models of *in vivo* metastasis and MLL1 depletion impairs multiple steps in the metastatic cascade (Fig. 6k). MLL1 regulates cell migration, a critical requirement for invasion and metastatic dissemination (*1, 88*); extravasation and metastatic outgrowth; as well as proliferation, which leads to primary tumor growth and hence, the number of cells invading into circulation.

### MLL1-Menin inhibitors combined with Paclitaxel block cell migration and proliferation

As new drugs in clinical trials are often tested in combination with existing standard of care, we assessed if MLL1-Menin inhibitors could be utilized in conjunction with current standard of care in a potential neoadjuvant or adjuvant setting. Paclitaxel is a chemotherapeutic that is routinely administered to patients with TNBC (*89, 90*), including in the metastatic setting (*91*). However, Paclitaxel suffers from toxicity-based limitations that restrict its dosage, leading to poor efficacy in clinic (*90, 91*). Doxorubicin, an anthracycline, is another chemotherapeutic routinely administered in TNBC (*89*). The impact of MLL1 depletion on diminishing tumor growth and metastasis *in vivo* hinted at the efficacy of MLL1-Menin inhibitors in a preclinical setting. The anti-proliferative effect of MLL1-Menin inhibition was additive with both Paclitaxel (Fig. 7a-d) and Doxorubicin (Fig. S10a). Lower dosage of MLL1-Menin inhibition (corresponding to Mode-1 MLL1 inhibition, which had minimal impact on proliferation) had a much smaller impact on cell numbers compared to either chemotherapeutic (Fig. S10b-c). Combination treatment of MLL1 inhibitor plus Paclitaxel was also effective in decreasing cell migration (Fig. 7e). Both MLL1 inhibition and Paclitaxel (*92*) decreased 3D cell migration, with combinations of MI-2-2 and Paclitaxel exhibiting the lowest cell velocities. These *in vitro* results indicate that the combination of paclitaxel and MI-2-2 simultaneously inhibits cell proliferation and migration, while inducing apoptosis.

The combination of MLL1 depletion in tumors and Paclitaxel treatment (Fig. 7f) led to a nearly flat growth curve (Fig. S10e) and the smallest tumors (Fig. S10f-g). Lung sections of shMLL1 tumor bearing mice treated with Paclitaxel showed the lowest metastatic burden, with virtually no darker staining visible in the lung (Fig. 7g and S10h). Administration of Paclitaxel to scrambled control tumors led to marginally lower metastatic burden than vehicle administration, which was not statistically significant (Fig. 7h). This decrease was attributable to a reduction in both size and number of metastatic lesions (Fig. 7i). Of note, treated mice did not exhibit any toxicity associated with Paclitaxel treatment either (Fig. S10i).

**Figure 7.**
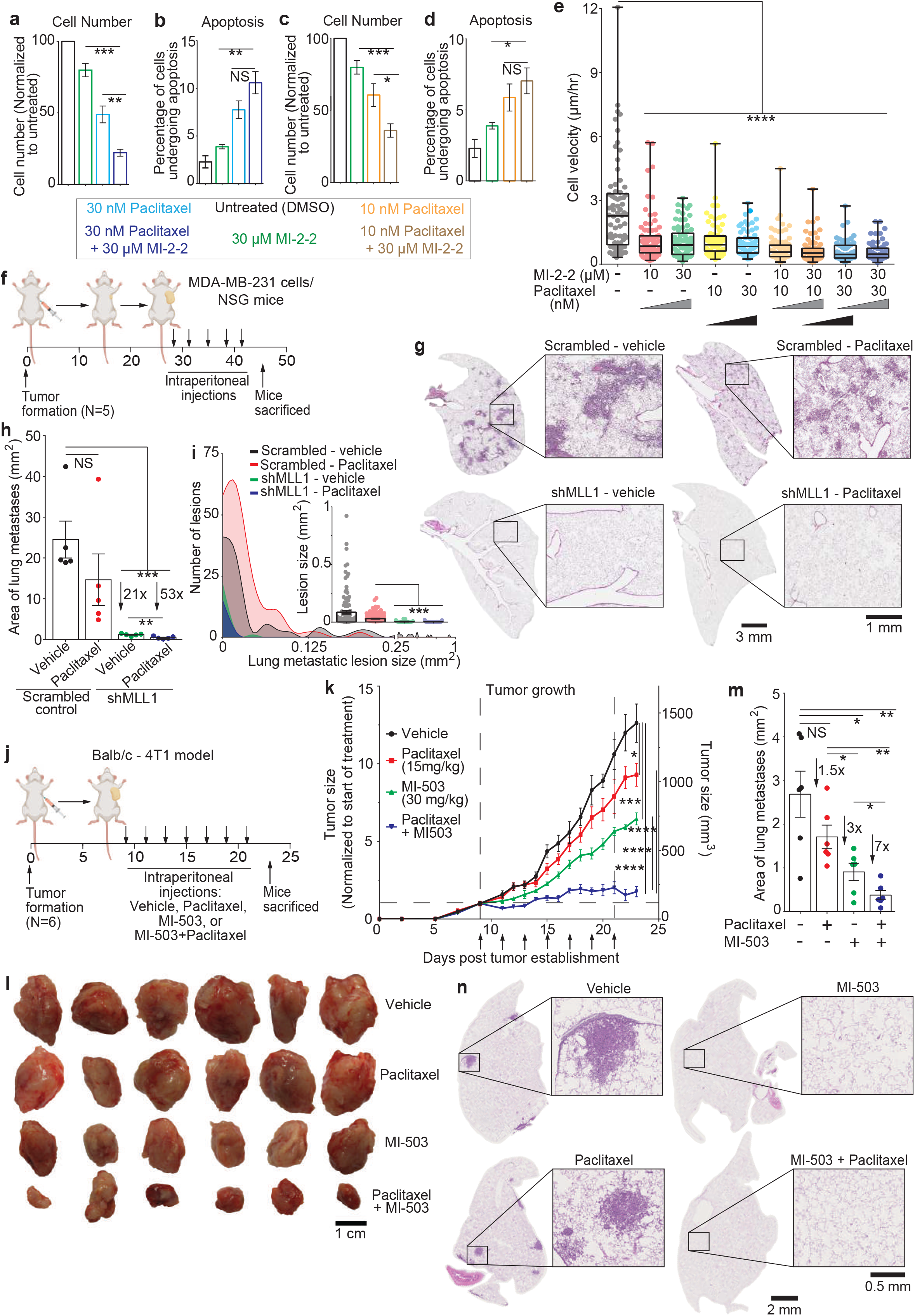
Concurrent paclitaxel and MLL1 inhibition arrests cell migration and proliferation. **(a- d)** Cytotoxicity of Paclitaxel is additive with the anti-proliferative effect of MI-2-2 *in vitro*. Combination of 30 µM MI-2-2 showed the lowest cell numbers (a and c) and the highest apoptosis (b and d) in MDA-MB-231. n(wells) = 4(each condition); one-way ANOVA (a: **** p < 0.0001, F (3, 12) = 27.57, b: ** p = 0.0032, F (3, 12) = 8.113, c: **** p < 0.0001, F (3, 12) = 75.65, d: **** p < 0.0001, F (3, 12) = 21.08). **(e)** MLL1-Menin inhibition or treatment with Paclitaxel reduces cell migration. Combination of MI-2-2 and Paclitaxel led to the lowest cell velocities. n(cells) = 90(Untreat.), 99(10 MI), 94(30 MI), 89(10 P), 84(30 P), 97(10 P+10 MI), 93(10 P+30 MI), 93(30 P+10 MI), 93(30 P+30 MI); one-way ANOVA (**** p < 0.0001, F (8, 823) = 27.87); two-tailed t-test (**** p(Untreat. – MI+P) < 0.0001, **** p(Untreat. – MI) < 0.0001, **** p(Untreat. - P) < 0.0001). **(f)** NSG mice bearing orthotopic MDA-MB-231 shMLL1 or control tumors were injected with Paclitaxel (25 mg/kg) or a vehicle intraperitoneally for five times. **(g)** Representative lung images from mice bearing control or shMLL1 tumors treated with Paclitaxel or vehicle. shMLL1 tumor bearing mice showed lower metastatic burden, and treatment with Paclitaxel also marginally reduced metastatic burden. **(h)** Quantification of metastatic burden per lung showed lowest burden for mice bearing shMLL1 tumors and treated with Paclitaxel. n(lungs) = 5(each condition, 1 lung/mouse); one-way ANOVA (** p = 0.0011, F (3, 16) = 8.832); two-tailed t-test (*** p(V,SC - P,sh) = 0.0007, *** p(V,SC – V,sh) = 0.0009, ** p(V,sh – P,sh) = 0.0058, NS p(V,SC – P,SC) = 0.2409). **(i)** Histogram of metastatic lesions in representative lungs show a reduction in both size and number of lesions with Paclitaxel treatment and/or MLL1-depletion. n(lesions) = 112(V,SC), 182(P,SC), 25(V,sh), 17(P,sh); one-way ANOVA (**** p < 0.0001, F (3, 332) = 10.79); two-tailed t-test (* p(V,SC - P,sh) = 0.0360, * p(V,SC – V,sh) = 0.0129, *** p(P,SC - V,sh) = 0.0002, *** p(P,SC – P,sh) = 0.0010). **(j)** Balb/c mice bearing syngeneic orthotopic 4T1 tumors were injected with MI-503 (30 mg/kg), Paclitaxel (15 mg/kg), a combination of the two drugs, or a vehicle intraperitoneally every alternate day for seven times. **(k)** The combination of MI-503 and Paclitaxel was effective in reducing tumor growth, with the growth of combination-treated tumors being effectively arrested. n(mice) = 6(each condition); one-way ANOVA (**** p < 0.0001, F (3, 20) = 36.50); two-tailed t-test (**** p(V – P+M) < 0.0001, **** p(P – P+M) < 0.0001, **** p(M – P+M) < 0.0001, *** p(V - M) = 0.0007, * p(V – P) = 0.0427). **(l)** The efficacy of combination treatment was reflected in final tumor sizes, with Paclitaxel + MI-503 treated tumors being the smallest. **(m)** Quantification of metastatic burden per lung showed that MI-503 treatment significantly reduced metastatic burden, while administration of Paclitaxel alone did not produce a significant decrease. Combination treatment (MI-503+Paclitaxel) led to a seven-fold decrease in metastatic burden compared to vehicle control. n(lungs) = 6(each condition, 1 lung/mouse); one-way ANOVA (*** p = 0.0003, F (3, 20) = 10.10); two-tailed t-test (** p(V – P+M) = 0.0016, *** p(P – P+M) = 0.0010, * p(M – P+M) = 0.0395, * p(V - M) = 0.0102, ** p(P – M) = 0.0375, NS p(V – P) = 0.1299). **(n)** Sample lungs from each condition show large but limited metastases in vehicle and Paclitaxel treated mice. MI-503 treated mice – alone or in conjunction with Paclitaxel – showed no major metastatic foci in their lungs. Data in this figure was generated with MDA-MB-231 cells *in vitro* (a-e), MDA-MB-231 cells *in vivo* in NSG mice (f-i), or with 4T1 cells *in vivo* in Balb/c mice (j-m).

Finally, as cytokines play a critical role in immune responses, we assessed the combination of MLL1- Menin inhibition with Paclitaxel in an immunocompetent model (*93, 94*). Balb/c mice bearing syngeneic orthotopic 4T1 tumors were subjected to MI-503+Paclitaxel treatment, or single drug controls, or vehicle (Fig. 7j). MI-503 was the MLL1-Menin inhibitor used as MI-2-2 is not suitable for *in vivo* administration due to poor metabolic stability (*59*). Both single drug controls slowed down tumor growth, with MI-503 being more efficient than Paclitaxel (Fig. 7k). Importantly, the combination of MI-503 and Paclitaxel nearly arrested tumor growth. The superior efficacy of MI-503+Paclitaxel was also evident in the final tumor sizes (Fig. 7l) and weights (Fig. S11b). Paclitaxel treatment did not reduce metastatic burden, but MLL1-Menin inhibition (either alone or in conjunction with Paclitaxel) led to a sharp decrease in lung metastasis (Fig. S11c), with combination treated lungs exhibiting virtually no metastatic lesions (evident in purple staining in Fig. S11d, metastatic burden quantified in Fig. 7m). High resolution sample lungs (Fig. 7n) showed that vehicle treated mice had large metastatic lesions (but fewer in number compared to MDA-MB-231 tumors) that was eliminated by MLL1-Menin inhibition (quantification and lesion size distribution in Fig. S11e). Crucially, the administration of MI-503 by itself or concurrently with Paclitaxel did not lead to significant toxicity, as evident by a steady mouse body weight (Fig. S11f).

In sum, our *in vitro* and *in vivo* results provide a pre-clinical rationale for the use of MLL1 inhibitors as anti-metastatic agents, and it is compatible for administration in conjunction with current TNBC chemotherapeutics.

## Discussion

### Biology of 3D cytokine-driven cell migration

Cytokines produced by cancer cells drive their migration, but the epigenetic regulation of cytokine production (*95–102*) and cell migration (*103, 104, 113, 114, 105–112*) are relatively understudied areas of cancer cell biology. Our work identifies MLL1 as an epigenetic regulator of cytokine-driven cell migration and metastasis. While MLL1 has been extensively studied in the context of MLL1-fusion driven leukemias, their biology is profoundly different from the proposed mechanism of cell migration in breast cancer cells ((*115*) and Fig. S3a-d). The metastatic cascade for liquid tumors is also different from that of solid tumors, particularly in the early stages, where migration plays an important role (*116*). MLL1-Menin interaction mediates cell motility in part via IL-6-pSTAT3-Arp3-based protrusion generation in cancer cells. IL-6 has been linked to breast cancer progression and metastasis (*117–124*), including in triple-negative cancer (TNBC) cells (*125*). Serum levels of IL-6 have been investigated as a potential prognostic marker in breast cancer (*126, 127*), but, until the current work, the upstream epigenetic regulators of this cytokine was unknown. The identification of these upstream proteins could provide additional avenues of therapy for cancers in which IL-6 plays an important role in progression.

In addition to establishing MLL1 as an important regulator of actomyosin machinery, our study also highlights Gli2 as the downstream effector of MLL1-mediated myosin contractility. The Gli family of transcription factors lie downstream of hedgehog signaling and essential for normal embryogenesis and development (*128*). Gli proteins have been reported to be upregulated and drive tumor progression in a variety of cancers including basal-cell carcinomas, medulloblastomas, and gliomas (*129*). Gli2 has been well-studied as a member of the TGF-β (*81, 82*) and Hedgehog (*130–132*) family. Gli2 promotes the establishment of bone metastasis in breast cancer via PTHrP-mediated osteolysis (*82, 131*). However, Gli2 has not been reported to be essential for cell migration or play a role in other steps of the metastatic cascade in breast cancer. Our work connects Gli2 to the well-established ROCK-pMLC2 axis in myosin-based cell contractility. Our work also highlights the potential of anti-Gli2 therapies in blocking metastasis of breast cancer. Gli2 inhibitors would not only block the spread of cancer cells from the breast, but also hamper the osteolysis-induced proliferation of cancer cells that colonize the bone.

### Potential clinical implications

Despite the fact that metastasis causes the vast majority of cancer-related deaths, there are currently no FDA-approved therapies in clinic that target metastasis directly (*133, 134*). Rather, they target metastasis indirectly as a byproduct of primary tumor shrinkage or attempt to shrink pre-existing metastatic lesions. Validation of MLL1-Menin interaction as a regulator of tumor metastasis in our preclinical models could add a new therapeutic target in TNBC, the subtype of breast cancer with the most limited treatment options. TNBC was chosen as the key focus for this study due to its relatively high metastatic rate. However, as noted, these observations could also be extended to other types of cancer. As this study investigates cell migration, a key step in metastatic dissemination, patients with Stage II and early Stage III TNBC, who are at the highest risk for metastatic spread, were the targeted demographic of this study. Epigenetic drugs have garnered interest as anti-cancer drugs and many are currently in clinical trials (*135*). However, the vast majority of the epigenetic drugs target families of epigenetic proteins rather than a specific protein (*136*). Our combination of MLL1-Menin inhibitor plus chemotherapeutic (Paclitaxel or Doxorubicin) would reduce metastasis in a two-pronged manner: By inhibiting key steps in the metastatic cascade – cell migration as well as extravasation and metastatic outgrowth - (via MLL1-Menin inhibitor), as well as by reducing primary tumor growth (reducing number of disseminating cancer cells; via MLL1-Menin inhibitor and chemotherapeutics). Inhibition of cell migration and proliferation by MLL1-Menin inhibitors was also readily reversible. Cell speed and proliferation were restored two days after withdrawal of drug treatment. This is consistent with MLL1 being an epigenetic protein and hence, the effect on migration and proliferation being plastic and reversible (*137*). Additionally, supplementation of three cytokines - IL-6, IL-8, and TGF- β1 - fully rescued motility, indicating that motility lost by MLL1-Menin inhibition was not due to poor viability/high stress in the cells following drug treatment. MLL1 is expressed in breast cancers and higher levels of MLL1 could be indicative of invasiveness and poor prognosis if the cancer has not spread. However, the lack of MLL1 expression being a clear prognostic indicator hints at a more multifaceted role for MLL1 in tumor progression than what is illustrated here.

We have utilized two chemotherapeutics with different modes of action: a taxane (Paclitaxel, Figs. 7 and S10-S11) and an anthracycline (Doxorubicin, Fig. S10). Paclitaxel and Doxorubicin are currently used in standard-of-care regimens in TNBC (*34, 138, 139*), and the ability of MLL1 inhibitor to work with these drugs indicate a potential for MLL1 inhibitors to be incorporated into the standard-of-care. MLL1 inhibitors are currently in Phase 1/2 clinical trials in relapsed or refractory acute myeloid leukemia (NCT04067336, NCT04065399, and NCT04811560), an indication of their capability to be utilized in clinical settings. Most clinical regimens are administered as combinations of drugs and/or antibodies, with combination therapies being better than mono-therapies (*140–142*). MLL1-Menin inhibitors could be incorporated into neoadjuvant chemotherapy regimens (including those containing taxanes and anthracyclines (*143*)), preventing the establishment of metastatic lesions concurrently - and independently - of primary tumor management. Once undergone breast surgery, MLL1-Menin inhibitors could be administered as a part of an adjuvant chemotherapy regimen such as sequential anthracycline-taxane regimens (*144*). Along with management of residual disease, MLL1-Menin inhibitors could also target the outgrowth of metastatic lesions. The usage of immunocompetent syngeneic mouse models was necessitated due to the central role of cytokines in MLL1-Menin’s regulation of cell migration and metastasis. However, since there is no appreciable difference in the magnitude of our observations between immunocompromised (which sets a baseline for the impact of MLL1 on metastasis) and immunocompetent (which factors in the baseline effect plus the impact of the immune system), it could be unlikely that the immune system is majorly responsible in mediating the reduced metastatic propensity caused by the impairment of MLL1-Menin interaction. Finally, the mouse models used in this manuscript primarily metastasizes to the lung, leading to assessment of lung metastatic burden as the primary indicator of metastasis. However, since cell migration is one of the first steps in the metastatic cascade (*145*), we expect MLL1 to play a vital role in the metastasis to other organs as well.

## Methods

### Reagent list

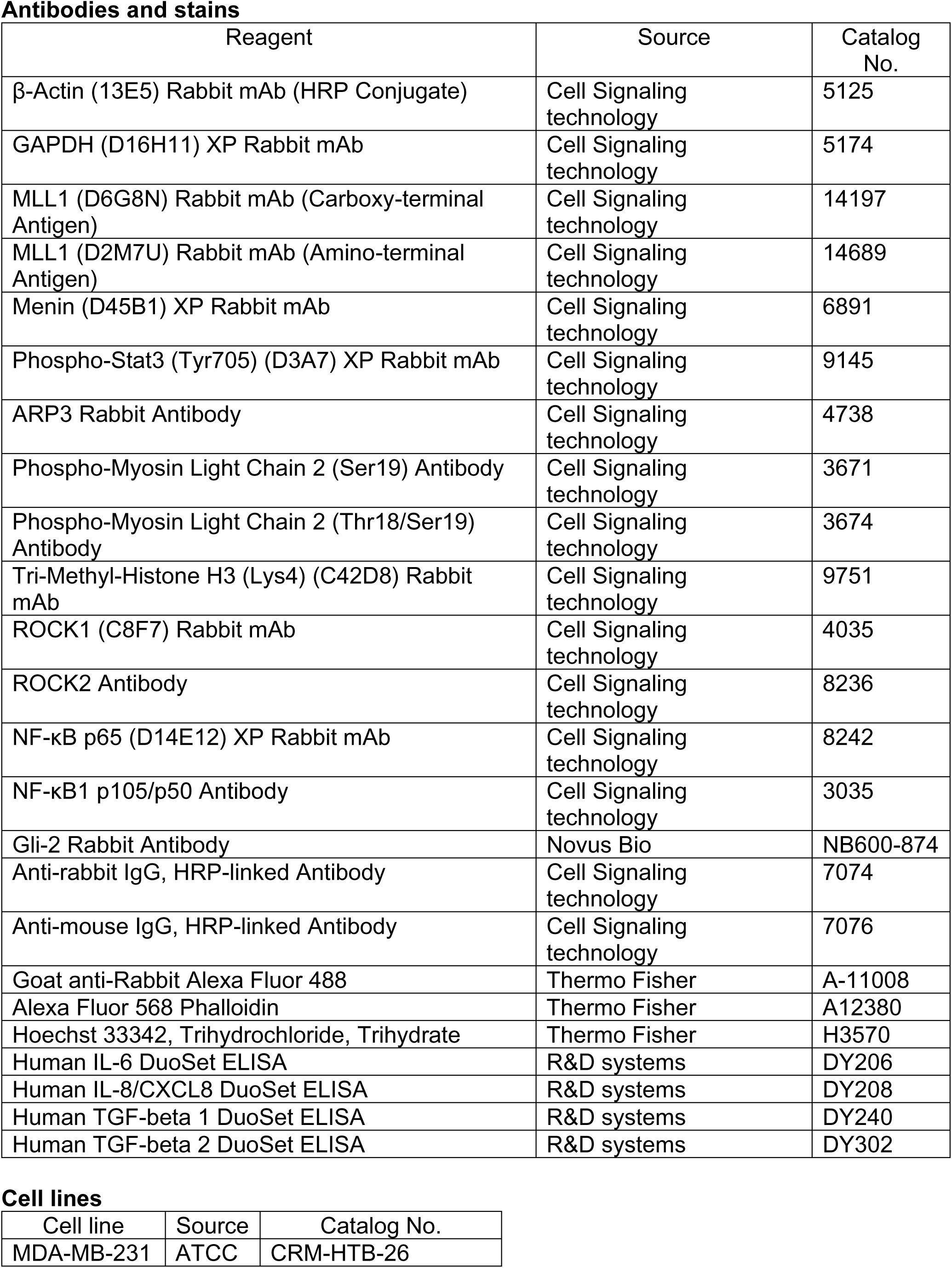
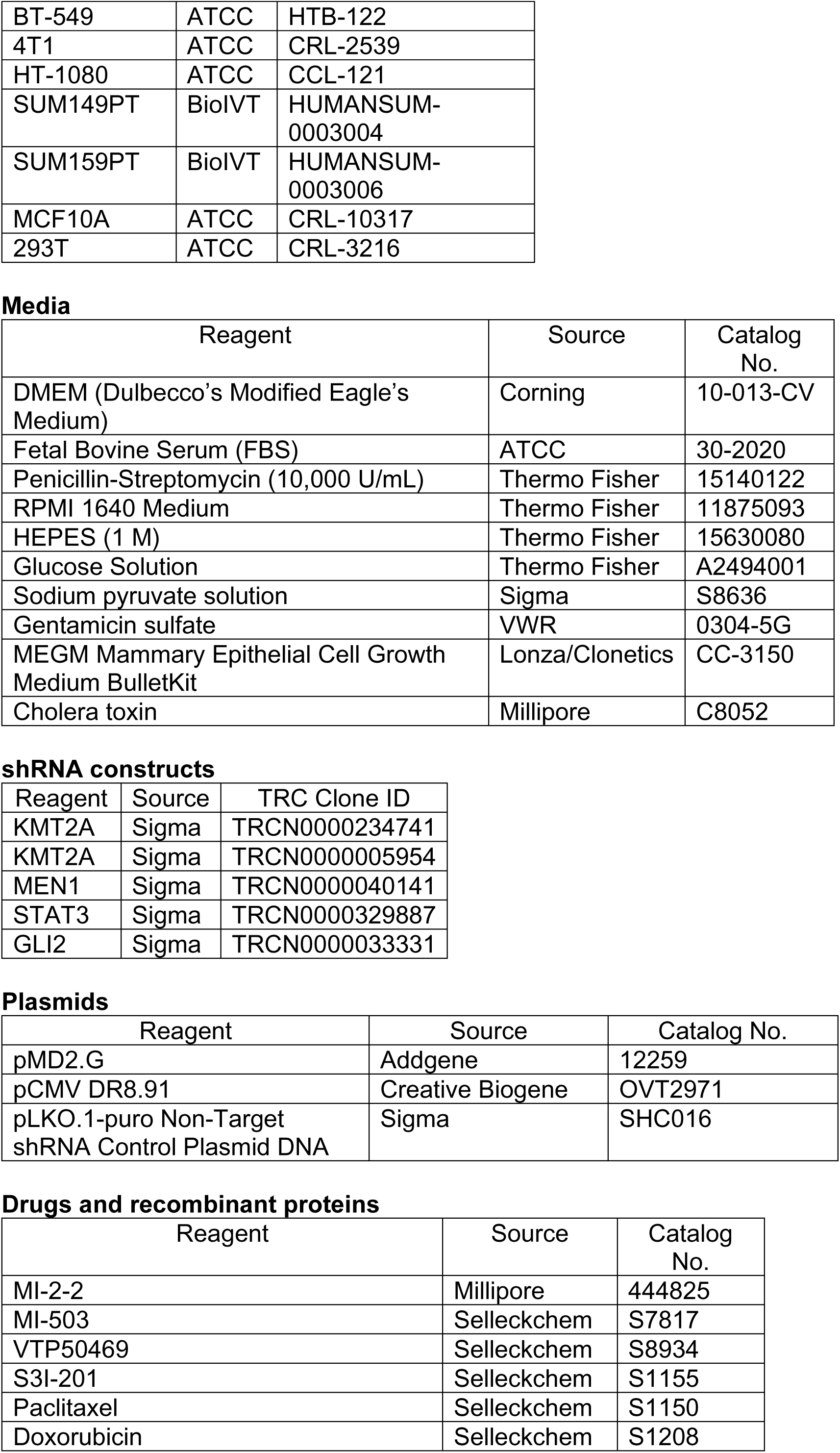
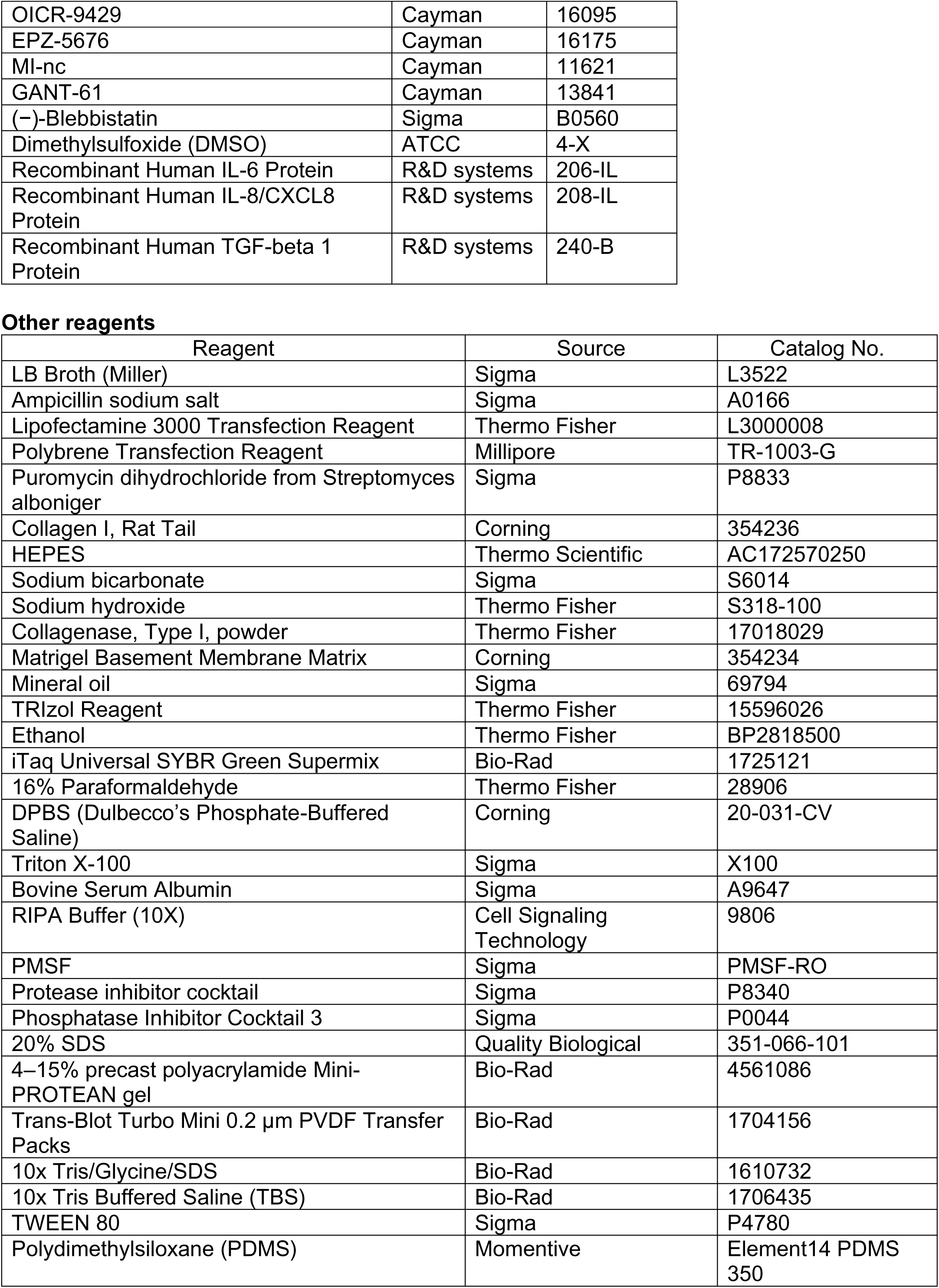
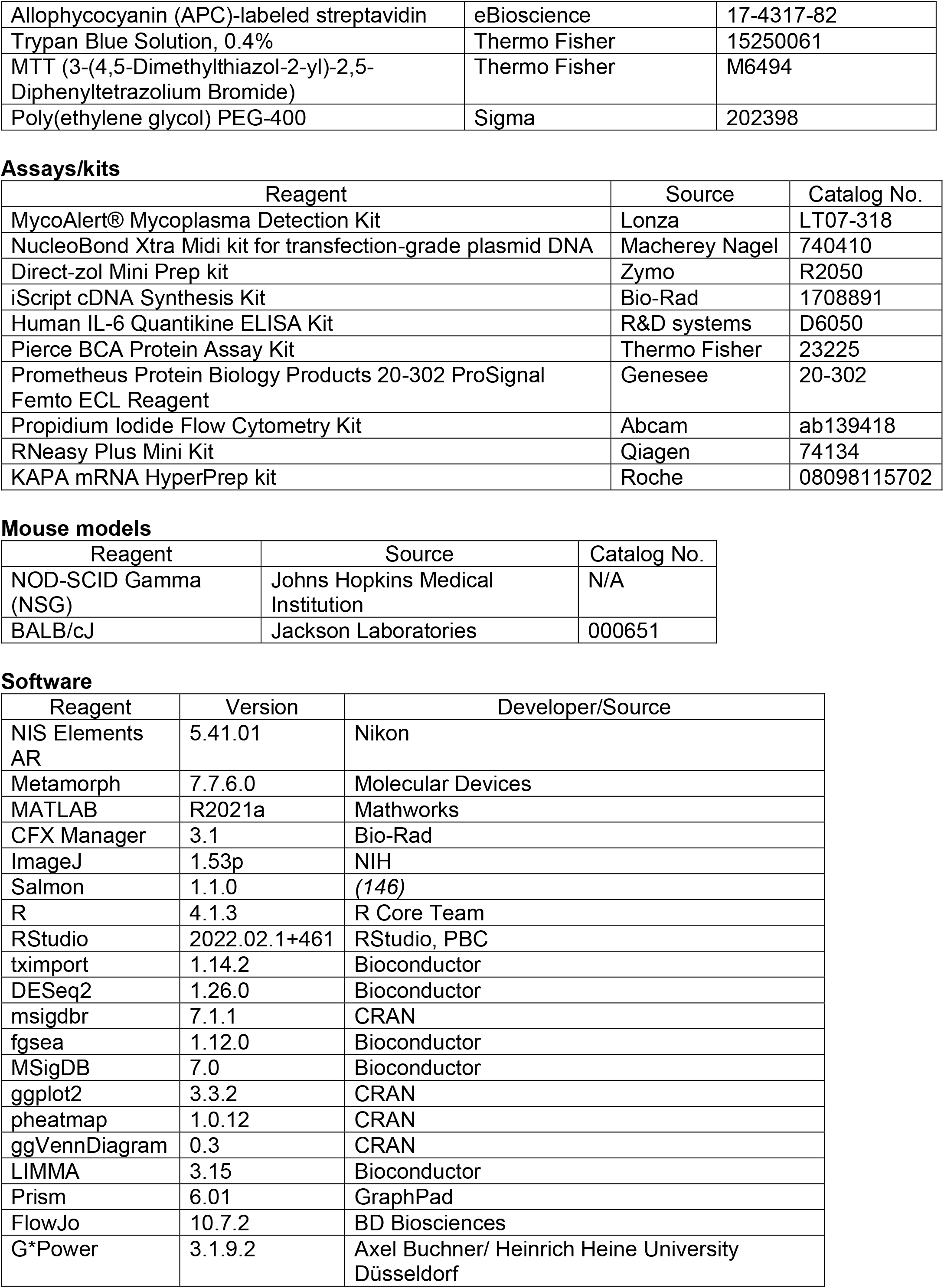
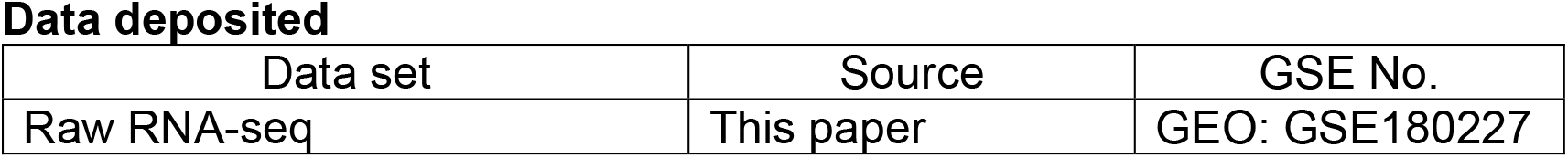

### Resource availability

Further information and requests for resources and reagents should be directed to and will be fulfilled by the lead contact, Denis Wirtz.

### Materials availability

This study did not generate new unique reagents.

### Cell culture

MDA-MB-231, and BT549 were purchased from ATCC and cultured with DMEM (Corning, with 4.5 g/L glucose, L-glutamine, and sodium pyruvate) supplemented with 10% fetal bovine serum (ATCC) and 1% penicillin streptomycin (Gibco). 4T1 cells were purchased from ATCC and cultured with RPMI 1640 (Gibco) supplemented with 10% fetal bovine serum (ATCC), 1% penicillin streptomycin (Gibco), 1% HEPES (Gibco), 1% glucose (Gibco), and 1% sodium pyruvate (Sigma). HT-1080 cells were purchased from ATCC and cultured with DMEM supplemented with 10% fetal bovine serum and 0.1% Gentamycin (VWR). SUM149 (SUM149PT) and SUM159 (SUM159PT) cells (BioIVT) were cultured with DMEM supplemented with 10% fetal bovine serum and 1% penicillin streptomycin. MCF10A cells were purchased from ATCC and cultured with MEGM™ Mammary Epithelial Cell Growth Medium BulletKit™ (Lonza/Clonetics) supplemented with not supplemented with 100 ng/ml cholera toxin (Millipore), and no GA-1000. These cells were cultured at 37°C and 5% CO_2_ for passage numbers less than 25 (15 for HT-1080) and tested for mycoplasma contamination every 6 months.

### MLL1, MEN1, STAT3, and GLI2 depletion by shRNA

MLL1 was depleted using two separate shRNA constructs (Millipore Sigma):

1) Gene: KMT2A; TRC Number: TRCN0000234741; Target Sequence: GATTATGACCCTCCAATTAAA (KD1 and KD2)

2) Gene: KMT2A; TRC Number: TRCN0000005954; Target Sequence: GCACTGTTAAACATTCCACTT (KD3)

MEN1 was depleted using the shRNA construct (Millipore Sigma): Gene: MEN1; TRC Number: TRCN0000040141; Target Sequence: GCTGCGATTCTACGACGGCAT

STAT3 was depleted using the shRNA construct (Millipore Sigma): Gene: STAT3; TRC Number: TRCN0000329887; Target Sequence: GTTCCTGAACATGATGACCTA

GLI2 was depleted using the shRNA construct (Millipore Sigma): Gene: GLI2; TRC Number: TRCN0000033331; Target Sequence: GCACAATCTACGAAGAATCAA

The bacterial glycerol stocks were grown overnight in sterile LB broth (Sigma) with 100 µg/ml Ampicillin (Sigma). Plasmid was extracted from the bacteria using a midi prep kit (Macherey Nagel) per the manufacturer’s instructions. The shRNA construct/plasmid, along with pMD2.G (pMD.G VSV-G) and pCMVDR8.91 (encoding Gag, Pol, Tat, and Rev), was co-transfected using Lipofectamine 3000. 293T cells were transfected with 10 µg of shRNA, 5 µg of pCMVDR8.91, and 1 µg of pMD.G VSV-G. The viral harvest was collected 48 hours after transfection and filtered through a 0.45 μm filter. MDA-MB-231 cells were incubated for 30 minutes with DMEM supplemented with polybrene at a final concentration of 8 µg/ml. After incubation, the media was replaced with fresh DMEM-polybrene, and 1 ml of filtered lentivirus was added (two separate replicates with one shMLL1 construct each). Cells were incubated with the lentivirus for 24 hours. Cells were selected with 2 µg/ml puromycin 72 hours after transduction. Knockdowns were verified via western blotting.

### 3D collagen gels and overnight live-cell tracking

Cells were trypsinized, counted, and resuspended in a 1:1 mixture of fresh media and reconstitution buffer [2.2% w/v NaHCO3 (Sigma Aldrich) and 4.8% w/v HEPES (Acros) in DI water] at pre-calculated cell concentrations. High concentration rat tail Collagen I (Corning) was added and was neutralized immediately with 1N NaOH. The solution was mixed thoroughly but gently (to avoid inducing air bubbles) and plated into well pre-heated at 37°C. Collagen gels were allowed to polymerize in the incubator for 1 hour, following which fresh media was added on top of the gels. The final cell density was 100 cells/mm^3^ and the collagen concentration in the gel was 2 mg/ml, unless specified otherwise. For low density (LD), cells were embedded at 10 cells/mm^3^.

Cell migration was assessed by overnight live cell tracking. As proliferation can influence “apparent” cell migration *in vitro,* traditional cell-migration assays (including transwell and wound-healing) preclude a direct ability to dissociate the contributions of proliferation from measured invasion. Our live-cell microscopy platform (*46, 147*) enables us to track the migration of individual cells, without the potential confounding effect of cell proliferation. Additionally, we have utilized new 3D techniques for more accurate representations of the microenvironment compared to flat culture dishes (*148–150*). Cells were maintained at 37 °C and 5% CO_2_ using a live cell box (Pathology Devices, Inc.) and imaged using a Cascade 1K CCD camera (Roper Scientific) mounted on a Nikon Eclipse TE200-E. Phase contrast images were recorded with a 10x objective every 10 minutes for a time frame of 20 hours. Cells were tracked using Metamorph imaging software (Molecular devices). A minimum of 100 cells were tracked (60 cells for LD) per condition. Migration parameters such as diffusivity, persistence time, and persistence speed were computed from measured cell trajectories using custom MATLAB codes (*46*). Cell velocity was calculated to obtain a more practical measure of motility and was calculated as follows:

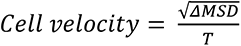

where ΔMSD is the absolute value of the difference in MSD between a time interval of T. In order to avoid errors introduced due to a non-steady state system a time scale was chosen that was longer than the persistence time but shorter than the total experiment time. T was set as half of the time length of experiment. For a total of N time points imaged, N/2 velocities were obtained (one each for the velocity between time points t_i_ and t_N/2+I_; I ranging from 1 to N/2) which were then averaged to attain the reported velocities.

### Generation of double-layered tumor spheroids

Tumor spheroids were prepared by immersing a core of cells in Matrigel into 2 mg/ml collagen gel as described in (*151*). Briefly, 50,000 cells were suspended in 1 µl Matrigel (Corning), pipetted into mineral oil (Sigma Aldrich) to form a sphere, and incubated at 37°C for 1 hour to polymerize the Matrigel. An outer layer of collagen was added by dipping the Matrigel cores in cold liquid collagen mixture (described in previous section) and repeating the abovementioned steps again. This architecture of a Matrigel core and an external collagen I matrix better mimics physiological conditions experienced by cancer cells and models two critical steps in metastasis, i.e., the invasion of cancer cells through the basement membrane followed by migration in the collagen-rich stromal matrix. These double-layered spheroids were added to 96-wells with 100 µl of fresh media. Scrambled control and shMLL1 spheroids were imaged at 4x magnification using phase-contrast microscopy.

### Drug treatments and cytokine supplementations

MI-2-2 was purchased from Millipore; MI-503, VTP50469, S3I-201, Paclitaxel, and Doxorubicin were purchased from Selleckchem; OICR-9429, EPZ-5676, MI-nc, and GANT-61 were purchased from Cayman Chemical; and Blebbistatin was purchased from Sigma-Aldrich. These drugs were dissolved in DMSO (except doxorubicin in water) and used at the concentrations shown in Supplementary Table 1.

Untreated (control) cells were treated with an amount of DMSO corresponding to the highest concentration of drug that the cells were treated with (and hence, the greatest volume of DMSO).

IL-6, IL-8, and TGF-β1 were purchased from R&D systems and reconstituted according to manufacturer instructions. IL-6 and IL-8 were supplemented at 375 pg/ml and 150 pg/ml, respectively. These correspond to the levels that these cytokines are expressed when embedded in 3D collagen gels at 100 cells per mm^3^ (*3*). IL-8 levels were only marginally reduced with MLL1-Menin inhibition, but it was also supplemented to avoid being a bottleneck following IL-6 supplementation. For the 3x IL-6/8 rescue experiments, IL-6 was supplemented at 1125 pg/ml and IL-8 at 450 pg/ml. TGF-β1 was supplemented at 10 ng/ml. The concentration reported in literature for cells on 2D matrices is 5 ng/ml (*157, 158*). However, TGF-β1 has been reported to adsorb strongly to collagen matrices with over 6 ng/ml (of the initial 10 ng/ml) getting adsorbed (*159*). Hence 10 ng/ml was supplemented in order to have ∼4 ng/ml available to cells.

Once cells were embedded in 3D collagen gels, cells were allowed to polarize and acclimate to the gel for 12 hours. After this time, media was exchanged for fresh media containing the appropriate drugs and/or cytokines at the concentrations indicated above. Cells were incubated with the drugs for ∼18 hours before being imaged overnight with live cell tracking. Total drug treatment/cytokine supplementation time for RNA or protein level analysis was two days unless stated otherwise.

### Assessing any potential off-target effects of drug treatment

The usage drugs as a primary method to elucidate the mechanism of MLL1-Menin mediated regulation of cell migration and proliferation necessitates the confirmation that observed effects and delineated mechanisms are not partly or entirely due to off-target effects of the drug. The various steps taken to ensure the fidelity and robustness of these mechanisms include:

1. <u>Choosing drug dosages that are consistent with that reported in literature</u>. Drug dosages for the key drugs (MLL1-Menin inhibitors MI-2-2 and MI-503) were chosen in-line with those reported in top journals (∼10-20 references each). The top reference for choosing the dosage of each drug is presented in the previous section.
2. <u>Validation of key proteins with knockdowns</u>. MLL1 (Fig. S2d), Menin (Fig. S2h), and its key downstream effectors STAT3 (Fig. S5d) and GLI2 (Fig. S6i) were depleted using shRNA and these cells were subjected to migration and mechanism-related studies.
3. Testing the effect of MLL1-Menin inhibition on a <u>normal non-cancerous breast epithelial cell</u> <u>line, MCF10A</u> (Fig. S4b-d). These cells were less susceptible to inhibition of proliferation and migration following MLL1-Menin inhibition.
4. <u>Using MI-nc as a negative control</u> (Fig. S4a). MI-nc has a similar structure to MI-2-2, but binds much more weakly to MLL1-Menin and does not inhibit MLL1-Menin interaction. MI-nc treatment at concentrations identical to MI-2-2 had no effect on migration.
5. <u>Using three well characterized and highly cited MLL-Menin inhibitors</u>. We have used MI-2-2, MI-503, and VTP50469 (Figs. 1e, S3j, and S3l); which span three generations, are synthesized by different groups, and are highly cited. All three MLL1-Menin inhibitors lead to identical cellular responses in migration, proliferation, apoptosis, and gene expression,
6. <u>Confirming full reversibility of pharmacological inhibition of MLL1</u> on proliferation and migration (Fig. S4h and S7e-f). Withdrawal of drugs lead to the recommencement of cell migration and proliferation within two days, the same period of time needed for these drugs to repress migration and proliferation.
7. <u>The absence of drug-induced apoptosis</u> (Fig. S4e-f). Reduced cell proliferation by MLL1-Menin was not accompanied by an increase in cellular apoptosis.

This data in addition to the extensive catalog of literature (see point 1) makes it highly unlikely that the observed effects are due to off-target effects.

Additionally, it is important to note that <u>drug availability is much more limited in 3D collagen matrices</u> <u>compared to a regular 2D system.</u> All the reference articles used a guide to choosing the optimal concentration of drugs have been performed in 2D systems in which the concentration of drugs apparent to the cells is the same as the concentration of drugs at which it was added. However, for greater physiological relevance, we embed our cells in a 3D Collagen I gel. Collagen is reported to be hydrophobic, leading to adsorption of hydrophobic molecules in the gel including hydrophobic drugs and proteins (see determination of TGF-β1 concentrations for supplementation in previous section). A similar effect would be in play regarding the drugs used in this manuscript, especially MLL1 inhibitors (which is a highly hydrophobic and insoluble in water), wherein the <u>apparent concentration of the</u> <u>drugs felt by the cells would be lower than the reported concentration</u> (it would be proportional to the amount of drugs added – amount adsorbed by gels).

### qRT-PCR

Cells were obtained by digesting the 3D collagen gels using collagenase. Gels were chopped into smaller pieces to increase their surface area and incubated with 1:1 ratio of collagen to type I collagenase (Gibco) dissolved in PBS at 2 mg/ml. After 1 hour, the solution was strained through a 40 µm strainer to remove any undigested collagen.

RNA was extracted using TRIzol (Thermo Fisher) and ethanol (Thermo Fisher) using a Direct-zol Mini Prep kit (Zymo). RNA concentration was measured using a NanoDrop spectrophotometer and 1 µg of RNA was reverse transcribed using iScript cDNA Synthesis Kit (Bio-Rad). Primers sequences were obtained from Harvard primerbank (sequences are listed in Supplementary Table 2) and synthesized by Integrated DNA Technologies. SYBR Green Master Mix (Bio-Rad) was used for fluorescence and the reaction mixture was amplified (40 cycles) using a CFX384 Touch Real-Time PCR Detection System (Bio-Rad). Gene expression was analyzed using Bio-Rad CFX Manager v3.1.

### Immunofluorescence

Immunofluorescence was carried out as described in (*154*). Briefly, cells were seeded in 6-well plates at a density of 10,000 cells/cm^2^. After drug treatment, cells were fixed using 4% paraformaldehyde (Thermo Fisher, diluted from 16%) for 10 minutes followed by three PBS (Corning) washes for 5 minutes each. Cells were then permeabilized with 0.5% Triton X-100 (Fisher) for 10 minutes followed by three PBS washes. Blocking was performed for 30 minutes using 5% bovine serum albumin (Sigma) followed by overnight incubation with primary antibody (Phospho-Myosin Light Chain 2 (Ser19) #3671, Cell Signaling technology, 1:100) at 4 °C. The next day, cells were washed three times with PBS and incubated with the secondary antibody (Goat anti-Rabbit Alexa Fluor 488, Thermo Fisher, 1:1000) and Alexa Fluor 568 Phalloidin (Thermo Fisher, for visualizing actin, 1:100) for 2 hours at room temperature. The solution was aspirated, and the cells were washed three times with PBS. DNA was stained using Hoechst 33342 (final concentration 10 µg/ml, Thermo Fisher) for 10 minutes, followed by three PBS washes. The stained cells were visualized with a Nikon A1 confocal microscope using a Plan Fluor 100x oil objective.

### Western blotting

For assessment at the protein level, gels were digested, and lysates were obtained from the cell pellet using RIPA buffer (Cell Signaling Technology) supplemented with 1mM PMSF (Roche), protease inhibitor cocktail (Sigma), phosphatase Inhibitor Cocktail 3 (Sigma), and 4% SDS (Quality Biological, diluted from 20%). The lysate was sonicated (5 short pulses at 30% power using Branson Sonifier 250), centrifuged at 10,000 rcf, and heated at 100 °C for 5 minutes. Protein concentration was measured using Pierce BCA Protein Assay Kit (Thermo Fisher) and 30 µg of protein was loaded per well/condition (unless stated otherwise). Lysates were loaded onto a 4–15% Mini-PROTEAN gel (Bio-Rad) and transferred onto a PVDF membranes using a Trans-Blot Turbo transfer system (Bio-Rad). Membranes were blocked with 5% BSA (Sigma Aldrich) in 0.1% Tween-TBS (Bio-Rad) and incubated with primary antibodies. Primary antibodies were purchased from Cell Signaling technology (β-Actin-HRP #5125, GAPDH #5174, MLL1 Carboxy-terminal Antigen #14197, Menin #6891, Phospho-Stat3 (Tyr705) #9145, ARP3 #4738, Phospho-Myosin Light Chain 2 (Ser19) #3671, Phospho-Myosin Light Chain 2 (Thr18/Ser19) #3674, Tri-Methyl-Histone H3 (Lys4) #9751, ROCK1 #4035, ROCK2 #8236, NF-κB p65 #8242, and NF-κB1 p105/p50 #3035), or Novus Bio (Gli-2, #NB600) and used at manufacturer’s recommended concentration for western blotting. After overnight incubation in primary antibodies, membranes were washed with Tween-TBS and incubated in secondary antibodies (Anti-rabbit IgG, HRP-linked Antibody #7074 or Anti-mouse IgG, HRP-linked Antibody #7076; both Cell Signaling technology) for 1 hour. Bound antibodies were detected with ECL western blotting substrate (Promega) and imaged using a ChemiDoc imaging system (Bio-Rad). Images were analyzed in ImageJ (NIH, (*160*)).

### ELISA assay

IL-6 secretion was measured using a Human IL-6 Quantikine ELISA Kit (R&D systems, #D6050) per manufacturer’s instructions.

### Protrusion analysis

#### 1. Cell body and protrusion segmentation method

The method to extract morphological measures from the phase contrast time-lapse videos consists of three steps: (1) cell body and protrusion segmentation, (2) cell tracking and (3) merge of the tracks and the segmentations to calculate the morphological measures for each cell along the time dimension.

##### 1.1 Cell body and cell protrusions segmentation

To segment each of the frames in the phase contrast microscopy videos, we trained a U-Net shaped convolutional neural network (CNN) (*161*) with manually annotated videos. To build the training set, we extracted 25 short videos of 15 continuous temporal points from the MDA-MB-31 dataset. Then, we annotated those videos with a label of 1 for the cell body and a label of 2 for the protrusions (Fig. S11f- h). To fit the input size of the network, we cropped patches from each of the annotated frames where there was a cell either focused or non-focused. Overall, we obtained a set with 5159 image patches from which 4642 and 516 were used as training and validation sets.

Our U-Net has four levels in the encoding path. Each level is composed by two 2D-convolutional layers and a 2D max-pooling. As described in (*161*), the convolutional layers on each of the four levels has a size of 32-64-128-256 convolutional filters, respectively. These are then followed by a bottleneck with two convolutional layers of 512 convolutional filters. The decoding path is analog to the encoding path with 2D upsamplings instead of max-poolings. Each level of the encoding path has a skip connection to the corresponding level in the decoding path. All the convolutional layers have a kernel of size 3 x 3 and ReLU as activation function. The last convolutional layer of the U-Net is followed by a convolutional layer with a kernel of size 1 x 1, 1 convolutional filter and the sigmoid activation function. In our case, the input to the U-Net was a series of the temporal frames (t – 1, t, t + 1) to output the binary mask of the cell body and protrusions at the time point t. Therefore, the size of the images entering the U-Net was 320 pixels x 320 pixels x 3 pixels and the output 320 pixels x 320 pixels x 2 pixels. The training was conducted over a pre-trained network. The pretrained weights were obtained from processing phase contrast microscopy images of fibroblasts. We fine-tuned our network for 200 epochs, with a batch-size of 10 and a decreasing learning rate starting from 5e^-6^. The categorical cross-entropy was chosen as the loss-function. At the end, we obtained a constant loss value of 4.1166 e^-04^ and an accuracy of 0.9880 for the training set, and a loss value 0.007 and an accuracy of 0.9878 for the validation set. We used the trained network to process the MDA-MB-231 image datasets. Each channel of the U-Net’s output was binarized with the threshold 0.5. Then, we ensured that all the pixels segmented as cell bodies were not segmented as protrusions and that all protrusions corresponded to a cell body. Those cells with a size smaller than 127 µm^2^ and all the protrusions with a size smaller than 6 µm^2^ were discarded. These values were estimated from the manual annotations.

To get the temporal morpho-dynamics of the cells, cells were tracked using MetaMorph software (Molecular Devices, Sunnyvale, CA, USA). The obtained tracks were merged with the previous masks by filtering out all those segmentations that did not have a track. We revised all the final masks and tracks to correct for those spurious segmentations and reduce the noise in the results.

We computed the following morpho-dynamics metrics for each cell and time-point *t* in the video: mean squared displacements (MSD) (µm), number of protrusions, and the length of the longest protrusion. MSD was computed as the average of the squared displacements made in a given time-window (*i*):

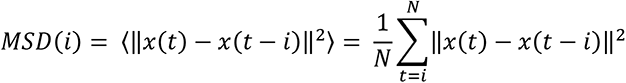

where x(t) represents the centroid of the cell body in Cartesian coordinates at the time-point *t*. The MSD value is expected to increase with *i* when the cells express a persistent behavior or travel long distances. For each cell, we calculated the number of protrusions using the mask of the protrusions assigned to that cell. The number of protrusions (P(t)) was calculated as the number of connected components in the protrusions mask. The cumulative number of protrusions at each time point was calculated as

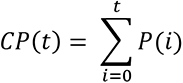

The length of each protrusion - each connected component - was given by the length of its skeleton. Then, the maximum length of each protrusion at each time point (L(t)) was calculated. Finally, the cumulative maximum protrusion length is given as

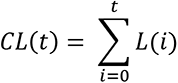

As cells enter and exit the plane of focus, their segmentation can be missing for certain time points. In those cases, we averaged each variable considering their value in the former and the latter time points.

### Multiplex cytokine assay

To perform profiling of secreted cytokines in the conditioned medium, multiplexed antibody barcode microarray chip was prepared as described previously (*162*). In brief, a polydimethylsiloxane (PDMS, Momentive, USA) template was combined onto poly-L-lysine coated glass slide and capture antibodies (Supplementary Table S3) were flowed into guiding channels on the PDMS template for overnight at room temperature. After detaching the PDMS template, 3% bovine serum albumin (BSA, Milipore Sigma, USA) in PBS solution was introduced onto surface of capture antibody coated glass slide, and then glass slides were kept in -20 °C fridge.

For running assay, the glass slide was overlaid with PDMS slab containing 24 sample loading chambers, which were prepared by biopsy punch (3 mm in diameter, Ted Pella, Sweden). 20 μl of conditioned media was added into the loading chambers followed by incubation for 12 hours at 4 °C. A cocktail of biotinylated detection antibodies (Supplementary Table S3) was introduced for 45 minutes at room temperature after detaching PDMS slab and rinsing with 3% BSA 3 times. Then, 5 μg/ml allophycocyanin (APC)-labeled streptavidin (eBioscience, USA) was added. After 20 minutes incubation at room temperature, the glass slide was again rinsed with 3% BSA 3 times, followed by blocking with 3% BSA for 30 minutes at room temperature. The glass slide was dried using nitrogen gas after dipping into 100 % DPBS, 50% DPBS, and DI water sequentially.

Scanning fluorescence images were obtained by using Genepix 4200A scanner (Molecular Devices, USA) with FITC (488 nm) and APC (635 nm) channels, and Genepix Pro software (Molecular Device, USA) was used to analyze and export fluorescence intensities. Antibody list for multiplex cytokine assay:

### Cell number measurement and proliferation assay

Cell numbers were counted by digesting the 3D collagen gels (described in previous section) and counting the number of cells using Trypan Blue (Thermo Fisher) stain. Cell proliferation (for kinetics) was determined using an MTT assay (Thermo Fisher) as described in (*154, 155*). 5000 cells were plated per well in 96-well plates. Once the cells had attached to the bottom, the existing media was exchanged for fresh media containing drugs. At set time points (0, 1, 2, and 3 days), media was aspirated, and cells were incubated with a 1:1 mix of fresh media and 5 mg/ml MTT solution in PBS. After 3 hours of incubation, media was aspirated, DMSO was added to solubilize the crystals, and absorbance was read at 550 nm. Apoptosis was measured using a propidium iodide kit (Abcam) per manufacturer’s instructions.

### RNA-seq: sample preparation, sequencing, and analysis

RNA isolation, library preparation, and sequencing were performed by Quick Biology. Total RNA was isolated and DNase I treated using Qiagen RNeasy Plus Mini Kit (Qiagen, Valencia, CA). The RNA integrity was checked by Agilent Bioanalyzer 2100 (Agilent, Santa Clara, CA) and samples with clean rRNA peaks (RIN>6) were used for library preparation. RNA-Seq library was prepared according to KAPA mRNA HyperPrep kit with 200-300 bp insert size (Roche, Wilmington, MA) using 250 ng of total RNAs as input. Final library quality and quantity were analyzed by Agilent Bioanalyzer 2100 and Life Technologies Qubit 3.0 Fluorometer (Thermo Fisher Scientific Inc, Waltham, MA). 150 bp PE (paired-end) reads were sequenced on Illumina HiSeq 4000 sequencer (Illumnia Inc., San Diego, CA).

Transcript level counts were quantified from reads with salmon version 1.1.0 (*146*) using human genome build h38. Quantification files generated by salmon were loaded into R/Bioconductor software (*163*). Data analysis was performed using with custom routines as well as standard packages. We perform paired differential expression analysis that retain cell-line pairing using a multi-factor design using the tximport (v 1.14.2) (*164*) and DESeq2 (v 1.26.0) (*165*) packages. The only protein coding genes were used. Genes with FDR adjusted p-values below 0.05 and log fold change greater than 1 were called statistically significant. Pathway analysis was performed by the msigdbr (v 7.1.1) and fgsea (v 1.12.0) (*166*) packages using hallmark pathways from the Molecular Signatures Database version 7.0 (MSigDB) (*167, 168*). Volcano plot were created using the ggplot2 package (v 3.3.2) (*169*), and heat maps were created using the pheatmap R package (v 1.0.12). Venn diagrams were created by ggVennDiagram (v 0.3). Identification of significant Hallmark gene sets following RNA-seq-based gene clustering was done by EnrichR (https://maayanlab.cloud/Enrichr/). Relevant gene sets were sorted based on their adjusted p-value.

### TCGA analysis

RNA-seq V2 data for basal breast cancer tumors from the Pan-Cancer Atlas (PanCanAtlas (*170*)) initiative of The Cancer Genome Atlas (TCGA) were used for analyses. Gene set statistics were computed with hallmark and curated gene sets from MSigDB (*171, 172*) and MLL1 expression using a one-sided Wilcoxon gene set test in LIMMA (*173*). Gene sets with FDR adjusted p-values below 0.05 were considered significantly enriched.

### ChIP-Atlas analysis

ChIP-Atlas (*76*) was used to acquire ChIP-seq data and viewed in Integrative Genomics Viewer (*174*) (IGV, ver. 2.10.0.06). ChIP atlas peak browser was used with the following parameters for hg38 data: Antigen class as TFs and others, Cell type class as all cell types, Threshold for significance as 50, and Antigen as KMT2A (MLL1), MEN1 (Menin), or WDR5. Cell type class was set as ‘all cell types’ to ensure sufficient KMT2A/MEN1/WDR5 studies with breast samples alone (< 5 each). Binding to a region of 2 kilobases (2 kb) flanking the TSS of each gene was considered as relevant.

### In vivo mouse modeling

All *in vivo* studies were performed in accordance with protocols approved by the Johns Hopkins University Animal Care and Use committee.

<u>MLL1 knockdown tumors:</u> Five-to-seven week-old female NOD-SCID Gamma (NSG) mice were purchased from Johns Hopkins Medical Institution and housed under a 12-h dark/light cycle at 25°C. 1 million MDA-MB-231 cells were suspended in a 1:1 mixture of PBS and Matrigel at 10^7^ cells/ml. 100 µl of cell suspension was injected into the second nipple of the left mammary fat pad to form orthotopic breast tumors. Tumor volume (measured with calipers) and body weight was monitored every two to three days. For survival study, mice were sacrificed when they reached the maximum tumor size permitted in ACUC protocol.

<u>Set tumor size threshold endpoint studies:</u> For tumor size control study, NSG mice were sacrificed when their primary tumors reached 1,400 mm^3^. Scrambled control and wild-type tumors grew faster and at an identical pace, and these mice were sacrificed on days 41 and 43, respectively. shMLL1 tumors grew slower and shMLL1-tumor bearing mice were sacrificed 62 days after tumor establishment. The lungs were excised, formalin fixed, paraffin embedded, sectioned, and stained with H&E (Hematoxylin and eosin). Lungs were imaged by the Johns Hopkins Oncology Tissue Services core (SKCCC). Lesion size and number were quantified using ImageJ (NIH).

<u>Tail vein injection of to assess extravasation and colonization:</u> MDA-MB-231 shMLL1, scrambled control, or wild type cells by were injected via the tail vein (500,000 cells/injection) in NSG mice as per (*175*). 100 µl cell suspension in PBS was injected into the tail-vein of a mouse using a 30-gauge syringe. Saline with no cells was the negative control. Mice were sacrificed six weeks after tumor establishment, the lungs were excised, fixed, H&E stained, and imaged.

<u>MLL1 knockdown plus Paclitaxel treatment:</u> shMLL1 or scrambled control tumors were established in NSG mice as described above. Four weeks after tumor establishment, mice bearing both tumor types were randomly assigned to Paclitaxel or vehicle treatment. Paclitaxel was administered at 25 mg/kg (*176*) by solubilizing in a vehicle [50% saline, 25% PEG-400 (Sigma Aldrich), and 25% Paclitaxel in DMSO; (*177*)]. Mice were injected intraperitoneally every 3 days for five times. All mice were sacrificed four days after the last injection and their tumors were excised and weighed. The lungs were excised, formalin-fixed, H&E stained, and fixed.

<u>MI-503 plus Paclitaxel treatment in syngeneic model:</u> Six week-old female Balb/c mice were purchased from Jackson Laboratories and housed under a 12-h dark/light cycle at 25°C. Mice were allowed to acclimate to the facility for a week before establishment of orthotopic breast tumors via the injection of 4T1 mouse TNBC cells (7000 cells/ 100 µl injection, suspended in a 1:1 mixture of PBS: Matrigel at 7x10^4^ cells/ml) as described above. Tumor volume (measured with calipers) and body weight was monitored every day. Nine days after tumor establishment, the mice were randomized and assigned to one of the treatment groups: vehicle (formulated as described above), Paclitaxel (15 mg/kg), MLL1- Menin inhibitor MI-503 (30 mg/kg), or a combination of Paclitaxel and MI-503. Half of the previously reported MTDs were chosen for Paclitaxel and MI-503 in order to avoid toxicity in combination treated mice. Mice were injected intraperitoneally seven times every alternate day. Mice were sacrificed 23 days after tumor establishment, when the tumors in the control (vehicle) group reached the maximum permitted tumor size of 1500 mm^3^. Tumor and organs were harvested from the mice as described above and subjected to the same analysis.

### Statistics

All *in vitro* experiments were repeated three times (n=3) as biological repeats. Measurement for each replicate was performed on distinct samples, and representative experiments are shown in figures. Each data point is represented by an individual dot where possible. Barplots represent mean ± s.e.m. For box-and-whisker plots, the upper limit of the box represents the 75th percentile, lower limit represents the 25th percentile, and the center line represents the median. The top and bottom whiskers stretch to the 99th and 1st percentile, respectively. Experiments were analyzed using one-way ANOVA (wherever number of conditions ≥ 3, following output parameters in figure captions – F(DFn, DFd) and p value). After confirming significance by ANOVA, significance between select key conditions were also analyzed by two-tailed t-test (results depicted in figures, output parameters in figure captions). p-values <0.05 was considered significant (*p<0.05, **p<0.01, ***p<0.001, ****p<0.0001, NS: Not Significant). Paired t-test was used where suitable (e.g. in rescues like Fig. 2m). All figure plotting and statistical calculations were performed using GraphPad Prism.

All migration and proliferation experiments were performed in technical replicates of 4 (N=4) per condition. For migration, a minimum of 80 cells were tracked (40 cells for LD) per condition. The exact number of cells/wells are noted in figure captions. Spheroid experiments were performed with a minimum of N=6 per condition and n=3. qRT-PCR was performed with N ≥ 4 per gene.

All *in vivo* experimental conditions were performed with a minimum of 5 mice bearing one tumor each (N=5). This N value provides a 73% power to detect a difference in mean tumor size between control and treated/shMLL1 groups. *A priori* power analysis was performed using t-tests in G*Power 3.1.9.2 software using the following parameters: alpha as 0.05, power of 0.65, allocation ratio (N2/N1) as 1, effect size (ρ) as large, and analysis with one-tailed t-test. For *in vivo* drug treatments, mice were randomized prior to initiation of treatment. For survival study, significance, hazard ratios, and 95% confidence intervals were calculated via logrank (Mantel–Cox) test.

## Data availability

RNA-seq dataset generated in this study will be available online upon publication in the GEO database under accession number GSE180227.

## Author’s Contributions

P.R.N. and D.W. conceived the project. P.R.N. and D.W. designed and performed the *in vitro* and *in vivo* experiments. L.D. and E.F. analyzed the RNA-seq data and did the TCGA analysis. E.G.M. and A.M.B. performed protrusion analysis. D.K. and R.F. performed the secretomic analysis. All authors contributed to the writing and editing of the manuscript.

## Acknowledgements

This work was supported by grants from the National Cancer Institute (U54CA143868) and the National Institute on Aging (U01AG060903) to D.W.; E.G.M. and A.M.B are grateful to Ministerio de Ciencia, Innovacion y Universidades, Agencia Estatal de Investigacion, under Grants TEC2015-73064-EXP and PID2019-109820RB-I00, MINECO/FEDER, UE, co-financed by European Regional Development Fund (ERDF), “A way of making Europe” (AMB); BBVA Foundation under a 2017 Leonardo Grant for Researchers and Cultural Creators (AMB); They also acknowledge the support of NVIDIA Corporation with the donation of the Titan X (Pascal) GPU used for this research.

We are grateful to Prof. Hari Easwaran, Dr. Ines Godet, Prof. Vito Rebecca, and Dr. Pei-Hsun Wu for carefully reading and editing the manuscript. Schematic illustrations were created using BioRender.

## Acronyms

CM: Conditioned medium
EMT: Epithelial to mesenchymal transition
GSEA: Gene Set Enrichment Analysis
H3K4: Histone 3 Lysine (K) 4
HD: High density (100 cells/mm3)
HR: Hazard Ratio
IL: Interleukin
LD: Low density (10 cells/mm3)
MLC: Myosin Light Chain
ND: Not detected
pMLC2: phospho-Myosin Light Chain 2
MLL1: Mixed Lineage Leukemia
MLL i: MLL1-Menin interaction inhibitor/inhibition
MSD: Mean squared displacement
SC: (Non-targeting) Scrambled shRNA Control
TCGA: The Cancer Genome Atlas
TGF: Transforming Growth Factor
TCGA: The Cancer Genome Atlas
TNBC: Triple negative breast cancer
TSS: Transcription start site
WT: Wild-type

Mode-1: The cell state/gene expression following low dose MLL1-Menin inhibition (10 µM MI-2-2 or 0.3-1 µM MI-503)

Mode-2: The cell state/gene expression resulting from a high dose MLL1-Menin inhibition (30 µM MI- 2-2 or 3 µM MI-503)

---

**Supplementary Figure 1.**
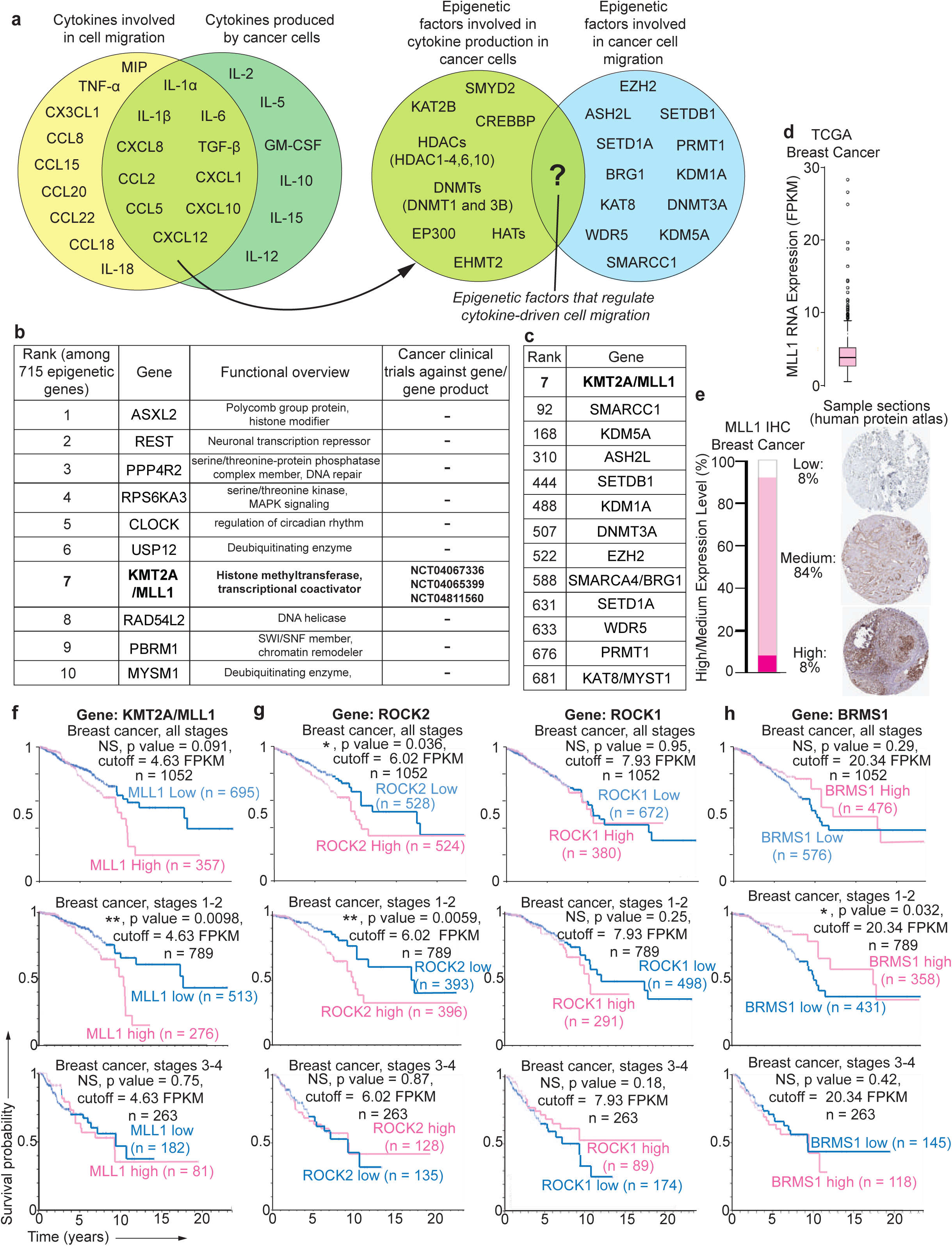
Analysis of TCGA and Human Protein Atlas suggests that MLL1 is a regulator of cancer-cell migration expressed in breast cancer. (a) Cell migration can be regulated in an autonomous manner by cancer cells via the secretion of cytokines. There is no overlap between the epigenetic factors that control the production of these cytokines and epigenetic proteins that have been reported to regulate cell migration. (b) Functional overview and clinical trial information for the top ten ranked epigenetic genes correlating best with migration-based genes in TCGA basal breast cancer dataset. Only MLL1 has been the subject of therapeutic interventions in cancer clinical trials, leading to a higher translational potential. (c) MLL1 was ranked much higher than other reported epigenetic drivers of migration shown in Fig. S1a. (d and e) MLL1 is expressed in breast cancers at the RNA and protein level (Source: Human Protein Atlas). (f) MLL1 was not prognostic across all breast cancers. The survival of MLL1 high tumors was not statistically significant than MLL1 low (p-value = 0.068, Human protein atlas KMT2A gene, n =1075, cutoff = 4.64 FPKM). Stratification of tumors into earlier stages (stages 1-2) and later stages (stages 3-4) shows that MLL1 is prognostic in earlier stages (which are yet to invade locally or metastasized) but not in later stages of breast cancer (which might have already metastasized). MLL1 might provide an advantage in metastasis, but once metastasized, the status of MLL1 might not be consequential. (g) ROCK1 and ROCK2 show similar prognostic trends when split into earlier and later stage breast cancer. (h) BRMS1, a breast cancer metastasis suppressor gene, also showed similar survival trend but with the exception that higher levels of BRMS1 correlated with better prognosis. All data in this figure was generated from TCGA (b-c) or human protein atlas (d-h).

**Supplementary Figure 2.**
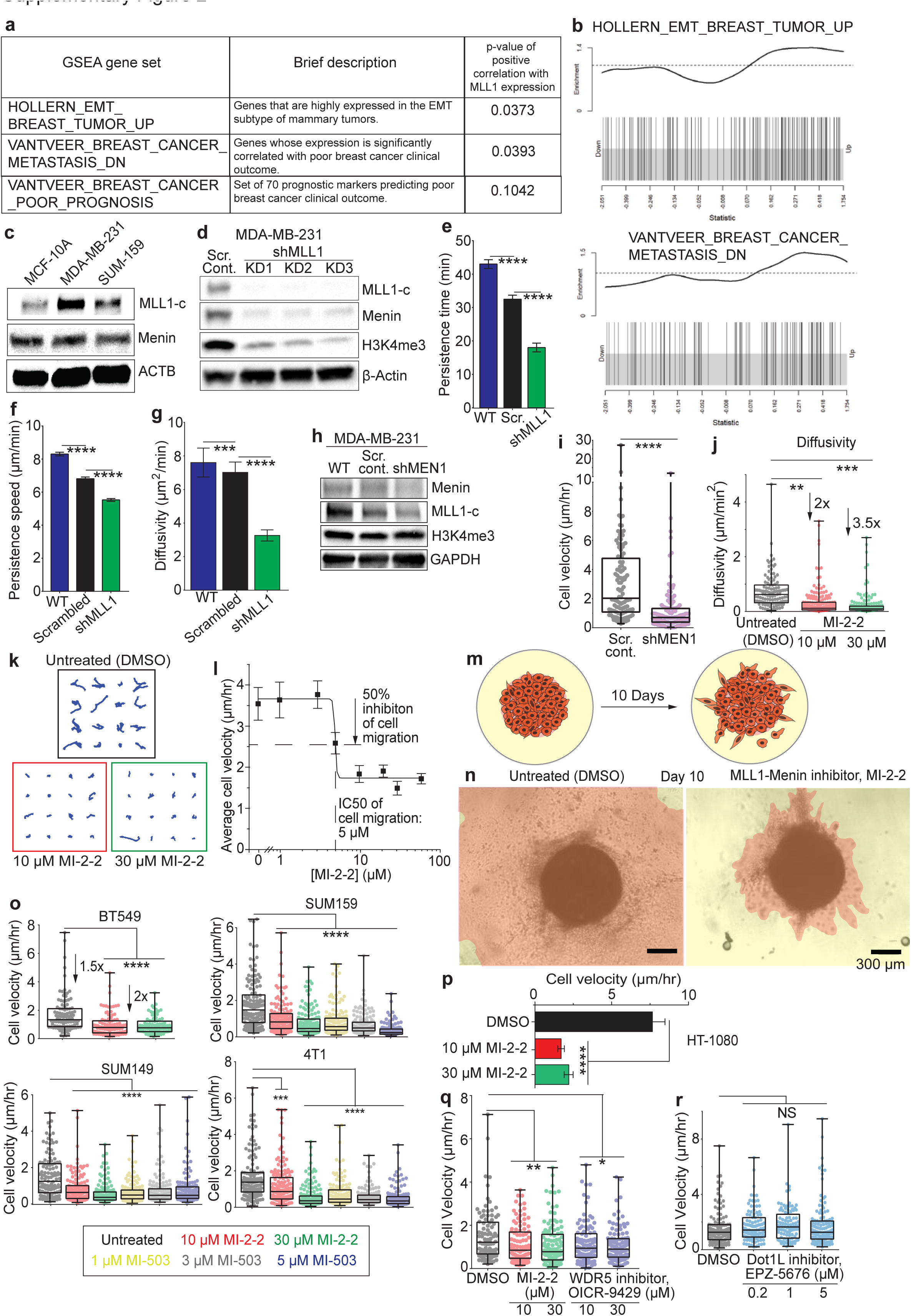
MLL1-Menin interaction regulates cell migration in tumor spheroids and in a panel of TNBC cell lines. (a) Two metastasis-associated GSEA gene sets were significantly and positively correlated with MLL1 expression. (b) Enrichment plots for the two significant gene sets in (a). (c) Levels of MLL1 protein are higher in metastatic TNBC cell lines (MDA-MB-231 and SUM-159) than breast non-tumorigenic epithelial cell line (MCF-10A). (d) MLL1 was depleted (>93%) from MDA- MB-231 cells using shRNA, leading to lower levels of H3K4me3. Actin was the loading control. Parameters associated with cell migration other than cell velocity, such as persistence time (e), persistence speed (f), and diffusivity (g) were also reduced by MLL1 knockdown. (h) Menin (MEN1) was depleted in MDA-MB-231 cells via shRNA. (i) Menin depletion reduces cell migration. (j) Cell diffusivities and (k) trajectories were reduced after MLL1-Menin inhibition by MI-2-2 at doses of 10 (red) or 30 µM (green). (l) Dose response curve for cell migration (measured as cell velocity) versus MI-2-2 dosage. The half-maximal inhibitory concentration (IC50) of cell migration was 5 µM for MLL1-Menin inhibition via MI-2-2. (m) MDA-MB-231 tumor spheroids were created by collagen-coating a core of cells embedded in Matrigel. With time, cells migrate and invade from the Matrigel core to the collagen outer layer. (n) The original spheroid core can be seen in the center as a black sphere and areas traced in orange indicate areas of collagen invaded by cells from the core. shMLL1 spheroids had a much smaller invaded area compared to untreated (DMSO control) tumor spheroids, indicated lower invasiveness of MLL1-inhibited cells. Scale bars 200 µm. (o) MLL1-menin inhibition suppresses cell migration in a wide range of TNBC cell lines. Inhibition of MLL1-menin in other TNBC cell lines: 4T1 (mouse), BT549, SUM149 and SUM159 (human) using the inhibitors MI-2-2 and MI-503 led to a significant reduction in velocity in all cell lines. (p) Outside of TNBC, MLL1-Menin inhibition also reduced cell migration in a human fibrosarcoma cell line, HT-1080. (q) MDA-MB-231 cells treated with a second- generation MLL1-menin inhibitor, MI-503, produced the same suppression in velocity as MI-2-2. (r) Dose response curve for cell velocity versus MI-503 dosage. The IC50 of cell migration was 40 nM for MI-503. Data in this figure was generated with MDA-MB-231 cells except panels a-b (GSEA gene sets), o (BT549, SUM-149, SUM-159, and 4T1), and p (HT-1080).

**Supplementary Figure 3.**
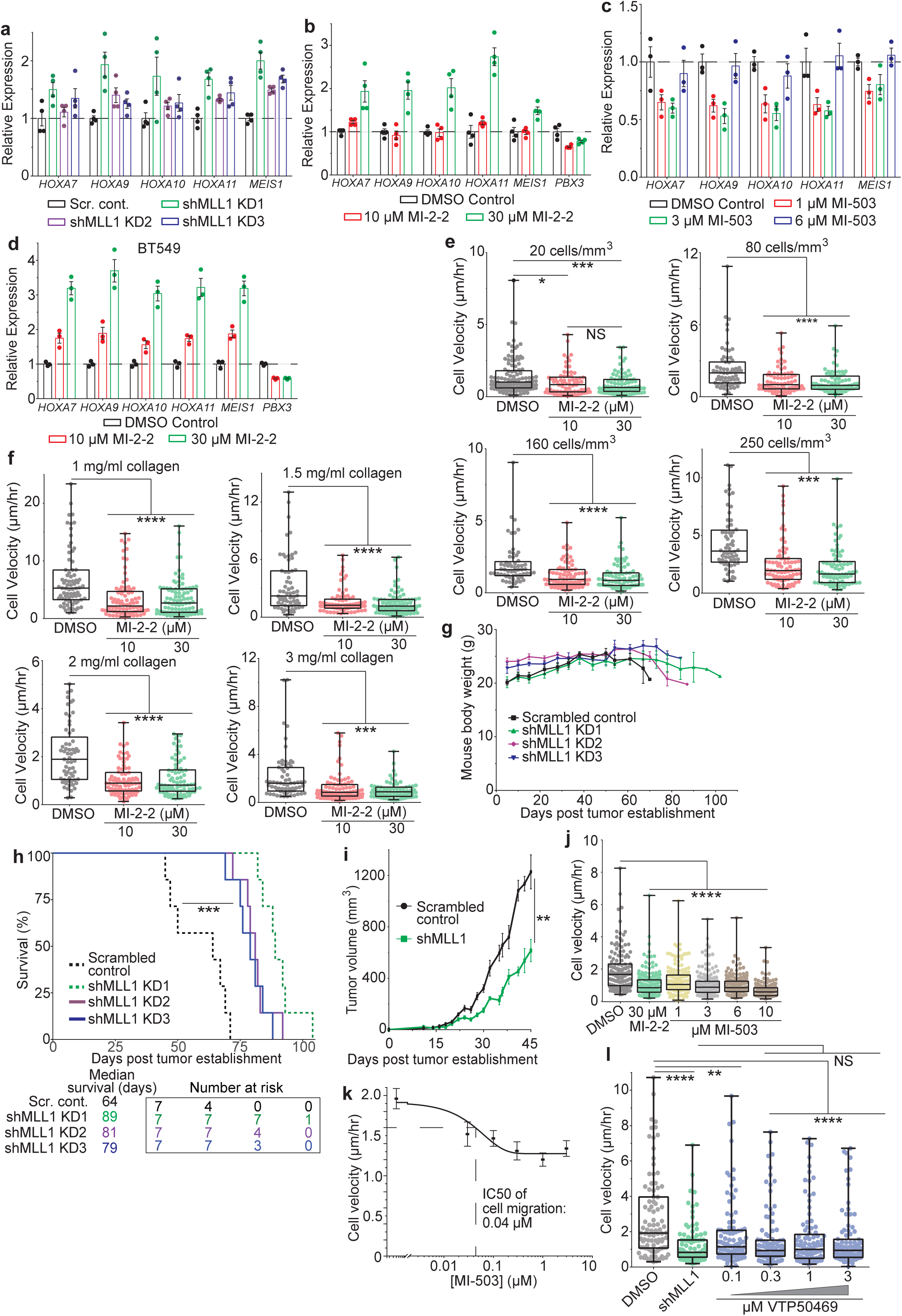
MLL1-Menin inhibition has a diminished impact on non-cancerous breast epithelial cells. (a) Another MLL1-Menin inhibitor, VTP50469, also inhibits cell migration. (b) MI-nc, a negative control for MLL1-Menin inhibition, did not alter cell migration. (c) Cells were treated with MI-2-2 prior to plating (pre-treatment) and subject to further treatment (post-treatment). (d) Recovery of cell migration after withdrawal of MI-2-2 in MDA-MB-231 cells and HT-1080 fibrosarcoma cells. (e) Migration of MCF10A (non-cancerous breast epithelial) cells was impacted to a lesser extent by MLL1-Menin inhibition compared to TNBC cells. (f) Concurrent inhibition of MLL1-Menin interaction plus MLL1 depletion does not reduce cell motility any further compared to either approach of disrupting the MLL1-Menin interaction alone. (g) MI-2-2 treatment did not appreciably increase apoptosis levels in MDA-MB-231 or (h) BT549 cells. (i-j) MLL1-Menin inhibition has diminished impact on proliferation of MCF10A cells compared to TNBC cells. (k) Inhibition of WDR5-MLL1 interaction also reduces cell migration to levels seen with MLL1-Menin inhibitors. (l) Inhibition of Dot1L, necessary for maintenance of MLL1-fusion leukemias, did not alter cell migration. (m-p) MLL1-Menin inhibition did not downregulate the expression of genes reported to be downstream of MLL1 in MLL-fusion leukemias such as Hox genes, *MEIS1*, and *PBX3*. Data in this figure was generated with MDA-MB-231 cells embedded in 3D collagen gels except panels d (HT-1080), e, i, j (MCF10A), and h, p (BT549).

**Supplementary Figure 4.**
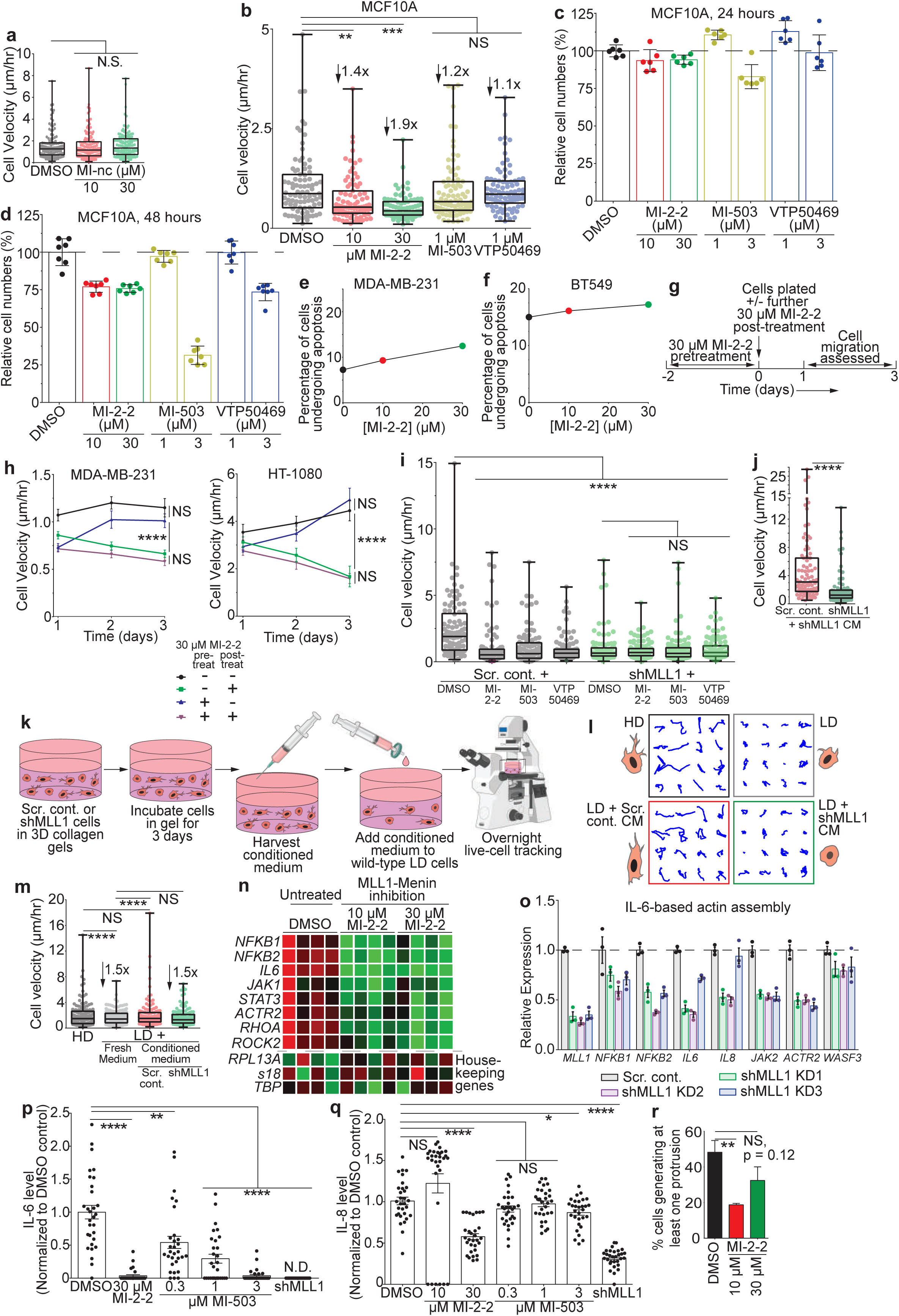
Conditioned medium from MLL1-depleted cells fails to rescue motility in low density cells. (a) Collagen density and (b) cell density did not affect the ability of MLL1-Menin inhibition to alter cell migration. (c) Mouse body weight for in vivo survival study in Fig. 1g-h. (d) NSG mice bearing other MDA-MB-231 shMLL1 clone tumors also survive longer than mice bearing scrambled control tumors. (e) Tumor growth curve for time-point study of scrambled control tumors vs shMLL1 tumors. (f) Addition of conditioned medium collected from shMLL1 cells to scrambled control or shMLL1 cells does not increase cell velocity compared to fresh medium in Fig. 2c. (g) Conditioned medium was collected from shMLL1 or control cells and added to WT cells at low density (LD). Cells with conditioned medium were then tracked overnight using live cell microscopy. (h and i) LD cells incubated with conditioned medium from scrambled control cells showed an increase in migration, while addition of conditioned medium from shMLL1 cells did not change their motility. (j) PCR heatmap of MI- 2-2 treated cells show downregulation of several key genes associated with cell motility and protrusion generation such as IL6, STAT3, ACTR2, and RHOA. RPL13A, s18, and TBP were the housekeeping genes. (k) shMLL1 cells showed downregulation of the same IL-6-STAT3 regulated actin assembly pathway, as was shown by MLL1-Menin inhibition. (l) Assessment of IL-6 production using multiplex cytokine assay shows that cytokine levels are reduced with MI-2-2 and MI-503 treatment, as well as with MLL1 knockdown. (m) In contrast to IL-6 production, MLL1 exerted only a weak effect on IL-8 production. Higher drug dosages or MLL1 knockdown were the only conditions that significantly altered IL-8 levels. (n) Percentage of cells producing at least one protrusion was reduced with 10 µM MI-2-2 treatment. Data in this figure was generated with MDA-MB-231 cells *in vivo* (c-e) or with MDA-MB-231 cells embedded in 3D collagen gels (a-b and f-n).

**Supplementary Figure 5.**
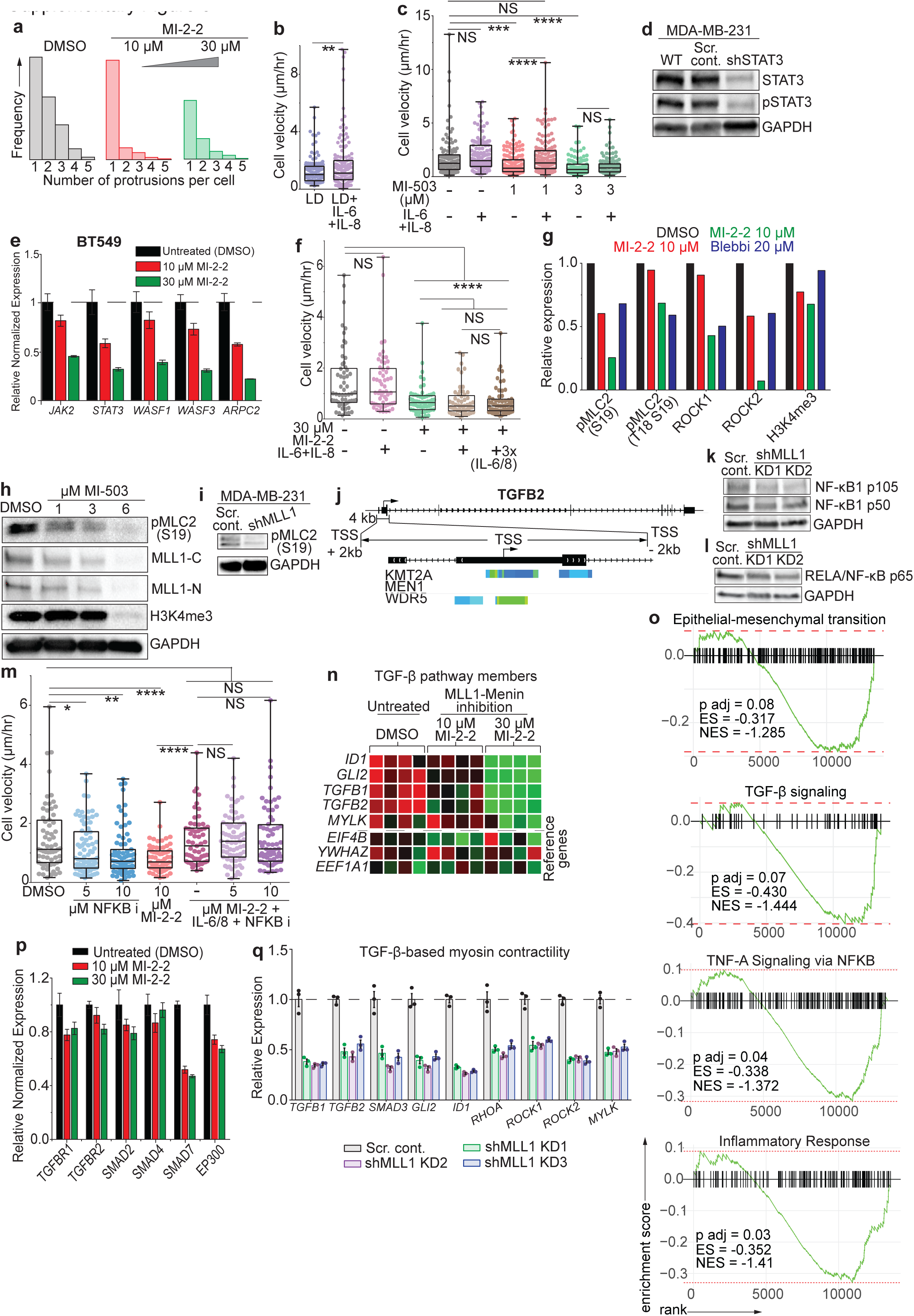
MLL1 depletion reduces expression of NF-κB1 and RELA. (a) Histogram of number of protrusions generated per cell show a sharp decrease in number of cells generating multiple protrusions. (b) Addition of IL-6+IL-8 to cells at LD increases their motility. (c) Migration lost after low dose (1 µM) MI-503 treatment was rescued by IL-6/8 supplementation. IL-6/8 supplementation did not rescue migration after a higher dose (3 µM) MI-503 treatment. (d) STAT3 was depleted in MDA- MB-231 cells via shRNA. (e) The same IL-6-STAT3 based actin assembly pathway was downregulated with loss of cell motility in BT549 cells. (f) Supplementation of IL-6/8 at three times the concentration did not rescue motility in cells treated with 30 µM MI-2-2. (g) Quantification of western blots after MI-2-2 treatment shows that pMLC2, ROCK1, and ROCK2 were downregulated after 30 µM MI-2-2 treatment. (h) pMLC2 was also reduced with MI-503 treatment. (i) shMLL1 cells showed reduced pMLC2 levels, indicating lower myosin contractility. (j) MLL1 and WRD5 bind to the promoter region of TGFB2. MLL1 depletion reduces expression of (k) NF-κB1 (p50 and p105) and (l) p65 RELA. (m) Inhibition of NF-κB decreases cell migration, but not in cells supplemented with IL-6/8. This is consistent with NF-κB lying upstream of IL-6/8 and controlling IL-6/8 production. (n) PCR heatmap of TGF-β pathway members show genes that are differentially downregulated with mode-2 (30 µM MI-2-2) but not with mode-1 (10 µM MI-2-2) MLL1 inhibition. Additionally, MYLK, involved in regulating myosin contractility, was also differentially downregulated. (o) GSEA plots for other key migration-related pathways. (p) Various members of the TGF-β family were downregulated after MI-2-2 treatment, including *TGFB1, TGFB2, SMAD3, SMAD7* and *EP300*, as assessed by qRT-PCR. (q) MLL1 knockdown/depletion reduces expression of genes central to TGF-β signaling and myosin contractility. Data in this figure was generated with MDA-MB-231 cells embedded in 3D collagen gels except panel e (BT549 cells).

**Supplementary Figure 6.**
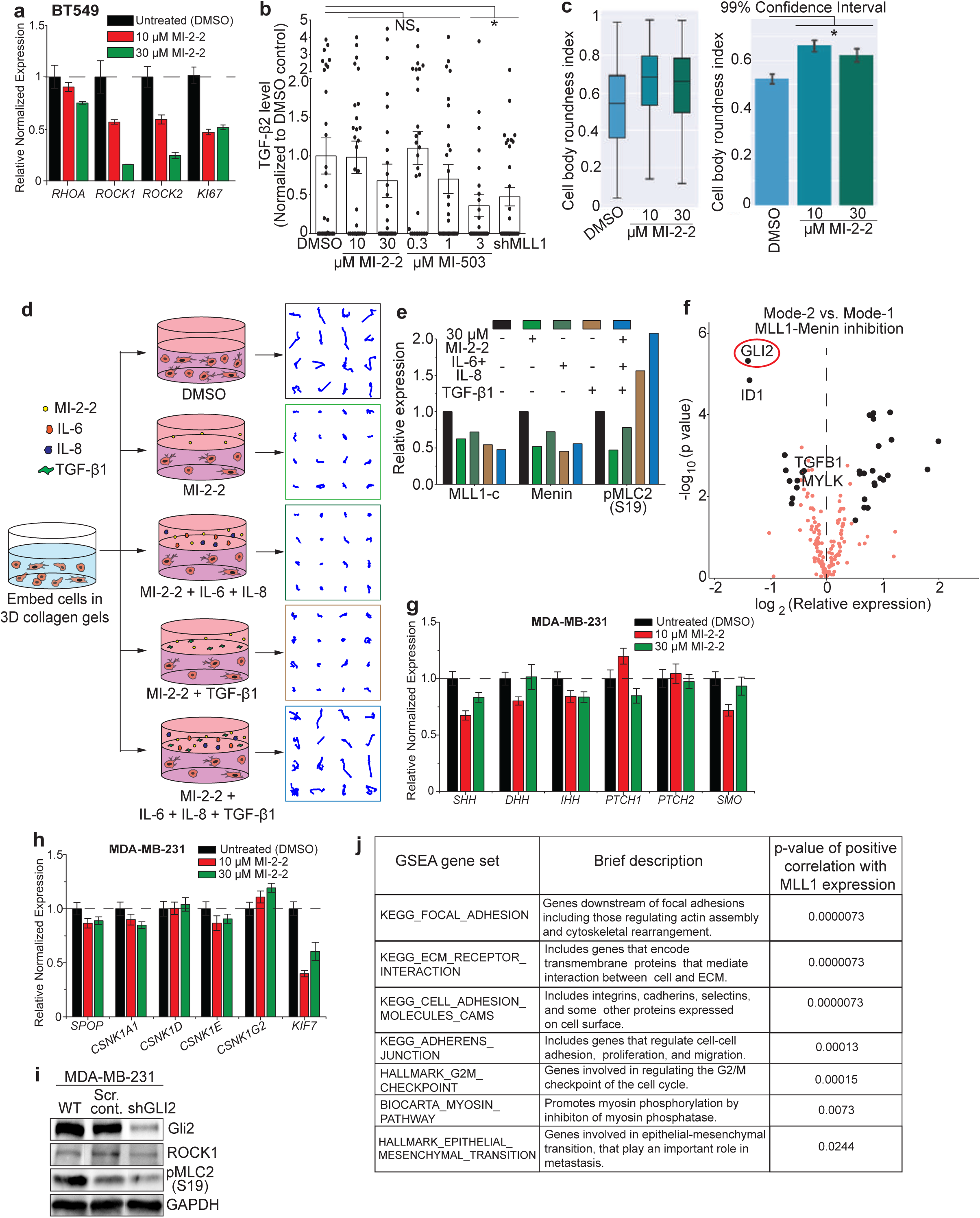
TGF-β1 supplementation rescues myosin contractility in cells depleted or inhibited for MLL1. (a) MLL1-Menin inhibition reduces expression of genes that regulate myosin contractility in BT549 TNBC cells. (b) Levels of TGF-β2 were largely unaffected by MLL1-Menin inhibition. Lower drug dosages produced no change in TGF-β2 levels, with high dose of MI-503 (3 µM) and MLL1 knockdown the only conditions to change TGF-β2 production. (c) Treatment with MI-2-2 increases cell roundness, analyzed by machine learning algorithm used for protrusion analysis. (d) Individual cell trajectories for cells treated with 30 µM MI-2-2 and rescued with TGF-β1+IL-6/8. (e) Quantification of the rescue western blot shown in Fig. 4a. pMLC2 levels were increased along with rescuing cell motility after TGF-β1+IL-6/8 supplementation. Levels of MLL1-c or Menin were not rescued by the supplementation. (f) A volcano plot of PCR-based expression showed *GLI2* as the most significant and most downregulated gene between mode-2 (30 µM MI-2-2) vs. mode-1 (10 µM MI-2-2) of MLL1-Menin inhibition. (g-h) MLL1-Menin inhibition did not appreciable alter the expression of ligands, receptors, and intermediates in the hedgehog signaling pathway, one of the pathways that lie upstream of the Gli proteins. (i) Gli2 was depleted in MDA-MB-231 cells using shRNA. (j) GSEA migration and proliferation-gene sets that were significantly correlated with MLL1 expression. Data in this figure was generated with MDA-MB-231 cells embedded in 3D collagen gels except panel a (BT549) and j (TCGA).

**Supplementary Figure 7.**
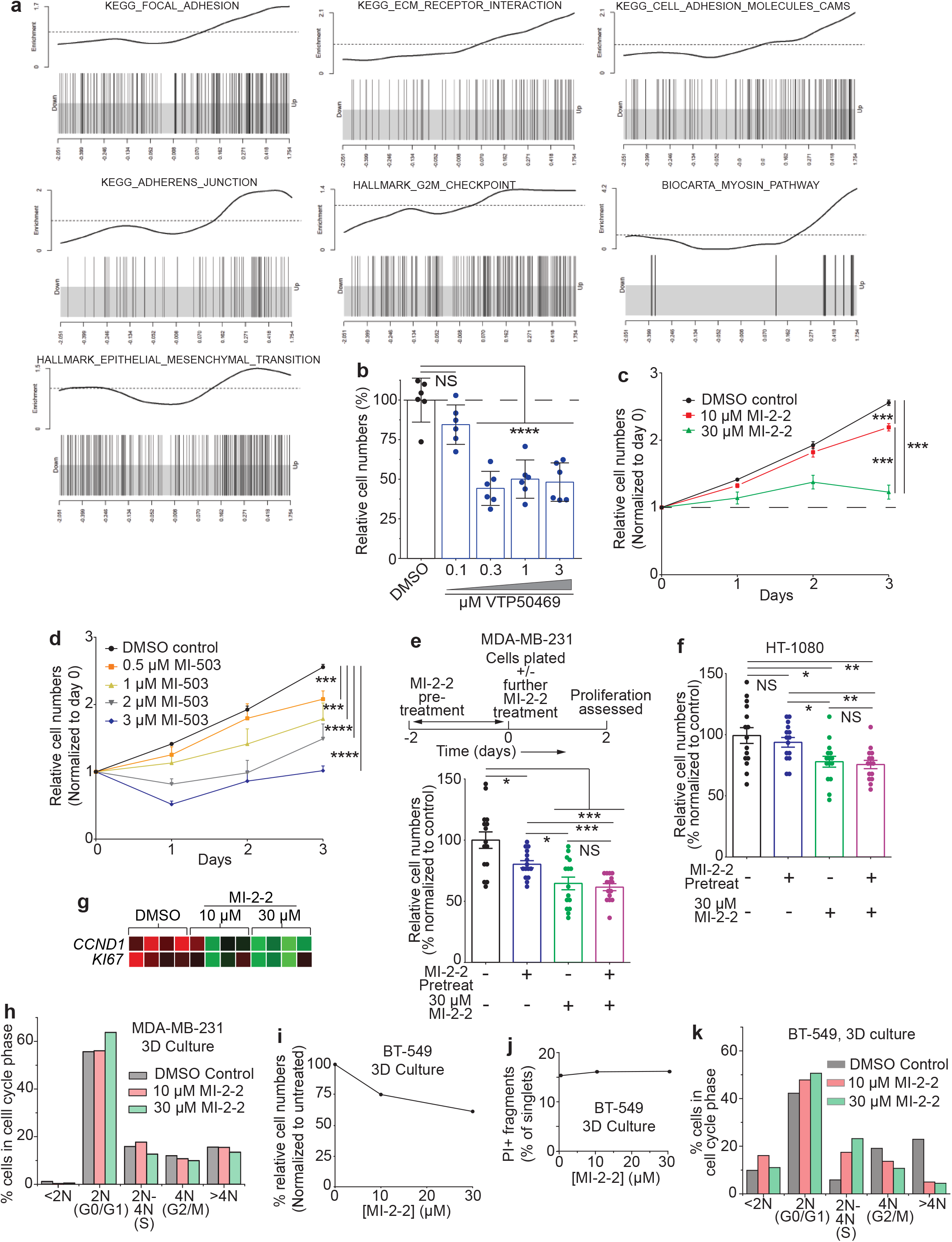
MLL1-Menin inhibition affects cancer cell proliferation. (a) Enrichment plots for the GSEA migration and proliferation-gene sets that were significantly correlated with MLL1 expression in Fig. S6j. (b) The newer MLL1-Menin inhibitor, VTP50469, also inhibits cell proliferation. (c) Growth kinetics of MDA-MB-231 cells shows that growth inhibition by MI-2-2 is evident within one day and continues henceforth. (d) Cell number kinetics of MDA-MB-231 cells treated with MI-503. MI- 503 has a dose dependent effect on proliferation, with 3 µM MI-503 halting cell proliferation. (e) Recovery of cell proliferation following termination of MI-2-2 treatment in MDA-MB-231 cells and (f) HT- 1080 fibrosarcoma cells. (g) Heatmap of qRT-PCR on *KI67* and *PCNA*, two proliferation related genes, show dose-dependent downregulation with MI-2-2 treatment. (h) Despite the growth inhibitory effect of MLL1-Menin inhibition, cell cycle distribution of MLL1 inhibited cells (10 or 30 µM MI-2-2 treated) was identical to untreated (DMSO) control. (i) MI-2-2 also reduced cell proliferation in BT549 TNBC cells. (j) MLL1-Menin inhibition by MI-2-2 does not increase apoptosis appreciably in BT549 cells. Hence, reduced cell numbers were due to reduced proliferation. (k) In spite of reduced proliferation, MI-2-2 did not alter cell cycle distribution in BT549 cells. Data in this figure was generated with MDA-MB-231 cells embedded in 3D collagen gels except panels a (TCGA), f (HT-1080), and i-k (BT549).

**Supplementary Figure 8.**
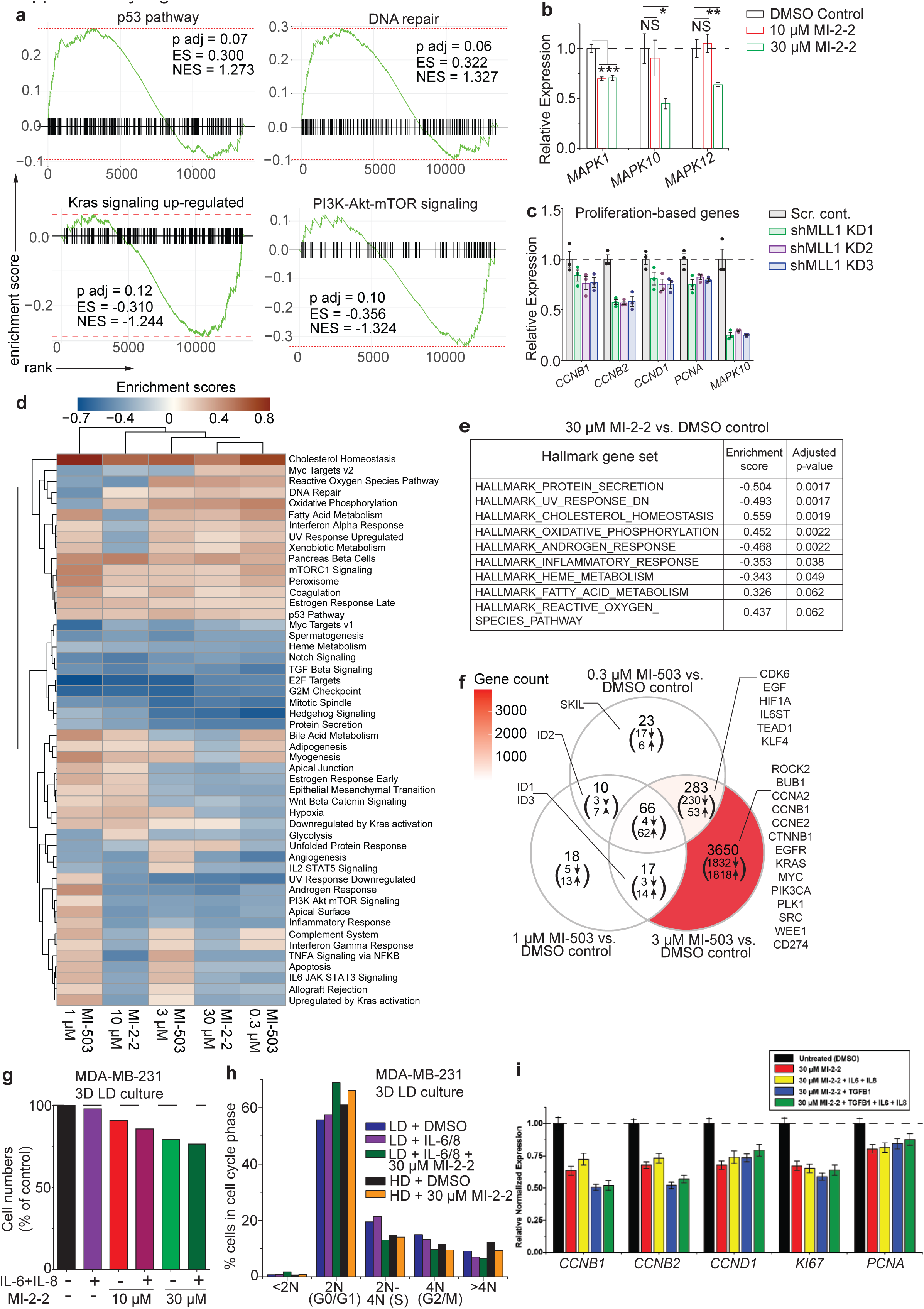
RNA-seq analysis of MLL1-Menin inhibition. (a) GSEA plots for other key proliferation-related pathways. (b) Key proliferation genes in the MAPK family: *MAPK1, MAPK10*, and *MAPK12*, were reduced at RNA level after MLL1 inhibition. (c) Similar trends for cell cycle related genes were observed with MLL1 knockdown. (d) Enrichment score heatmap on all Hallmark gene sets show an alteration in several gene sets. (e) Hallmark gene sets that were significantly up or down-regulated after 30 µM MI-2-2 treatment. (f) Venn diagram showing the number and names of select significantly altered genes with each MI-503 treated condition and the overlap between these conditions. (g) Unlike cell migration, reduced cell proliferation after MI-2-2 is not rescued by supplementation of IL-6/8. (h) Supplementation of IL-6/8 does not alter cell cycle distribution either. Control cells, cells treated with MI-2-2, and cells treated with MI-2-2 and supplemented with IL-6/8 showed similar cell cycle distributions. (i) Supplementation with TGF-β1 and/or IL-6/8 did not change expression of key proliferation marker genes or cell cycle related genes. All data in this figure was generated with MDA-MB-231 cells embedded in 3D collagen gels.

**Supplementary Figure 9.**
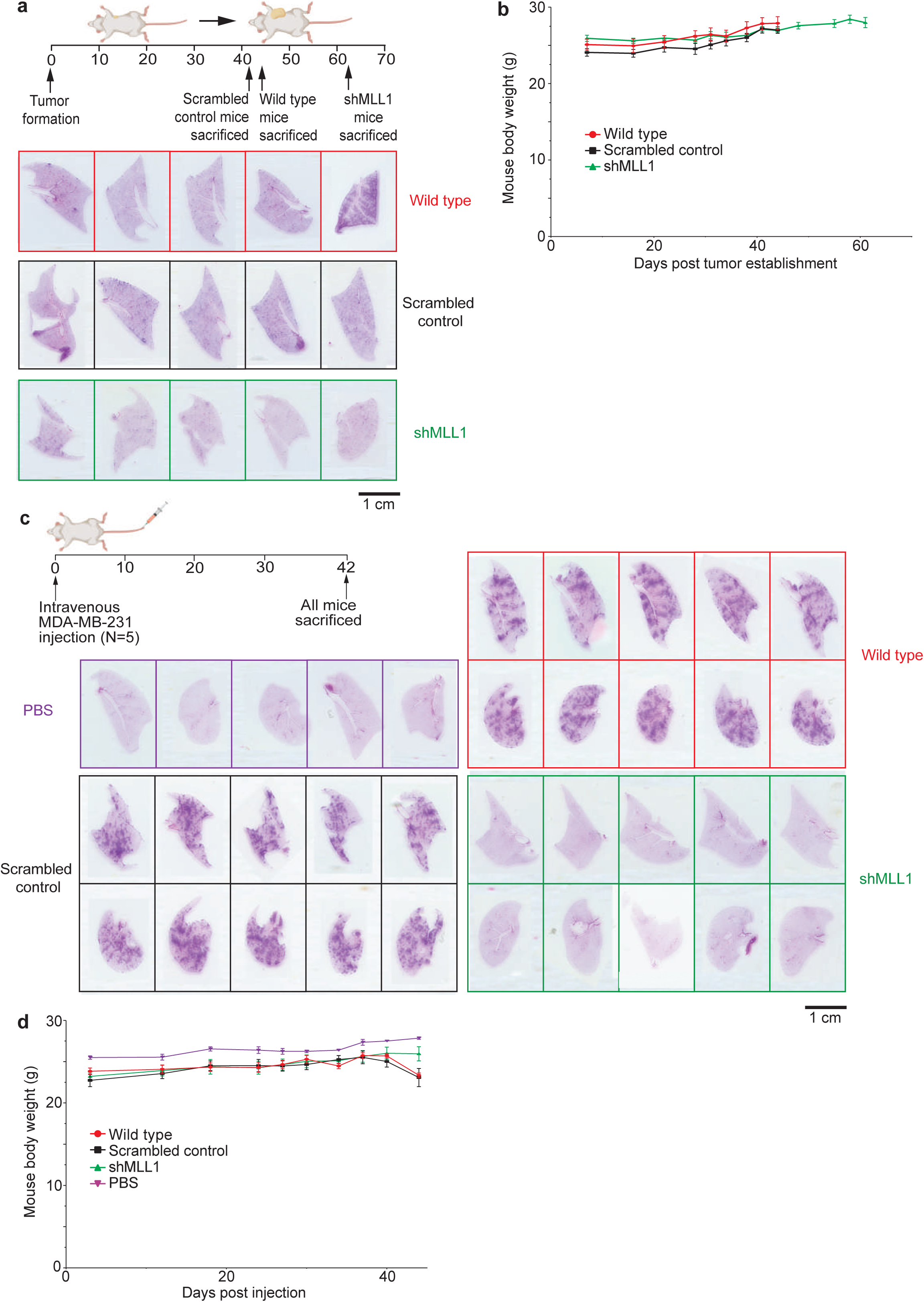
MLL1 depletion reduces lung metastatic burden in tumor bearing mice. (a) Full panel of lungs from mice bearing wild type, scrambled control, and shMLL1 tumors. Mice were sacrificed when they reached a set tumor size. (b) Mice body weights for tumor size control study. (c) Full panel of lungs from mice that received tail-vein injections of wild type, scrambled control, and shMLL1 cells. Lungs from mice injected with saline (PBS) was the negative control. (d) Mice body weights for tail vein metastasis model. All data in this figure was generated with wild type, scrambled control, or shMLL1 MDA-MB-231 cells *in vivo*.

**Supplementary Figure 10.**
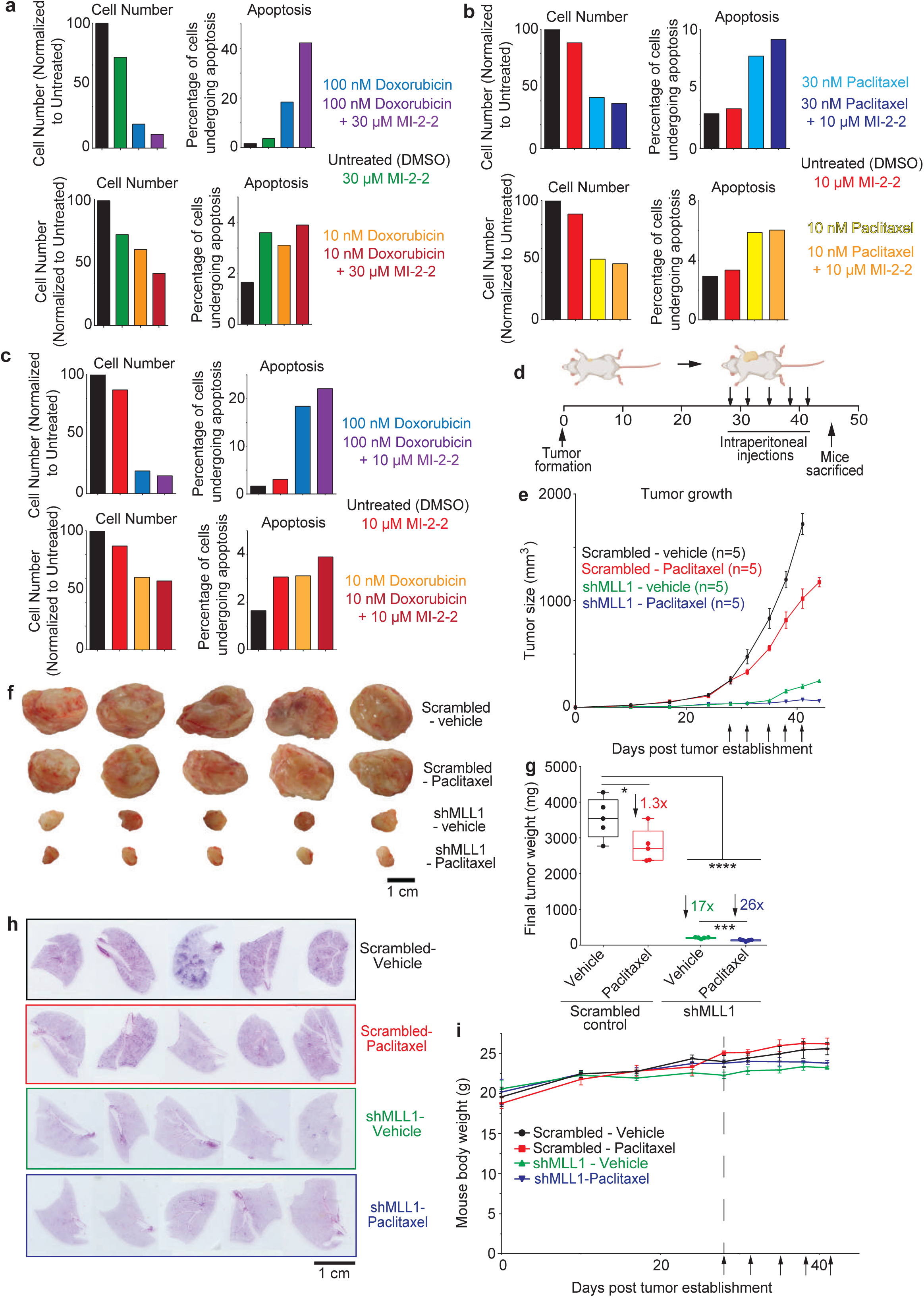
MLL1-Menin inhibition is additive with TNBC chemotherapeutics *in vitro* and *in vivo*. (a) MI-2-2 is additive with Doxorubicin, with 100 nM Doxorubicin and 30 µM MI-2-2 leading to the lowest cell numbers and inducing the most apoptosis. (b) Combination of 10 µM MI-2-2 with Paclitaxel. (c) Combination of 10 µM MI-2-2 with Doxorubicin. (d) NSG mice bearing orthotopic MDA-MB-231 shMLL1 or control tumors were injected with Paclitaxel (25 mg/kg) or a vehicle intraperitoneally for five times. (e) shMLL1 tumor bearing mice treated with paclitaxel showed a near- arrest of tumor growth, (f) showed the smallest tumor sizes (scale bar is 1 cm), and (g) had the lightest tumors on excision. (h) Full panel of lungs from mice bearing either scrambled control or shMLL1 cells and treated with Paclitaxel or vehicle. H&E staining of lungs show darker staining in scrambled tumors treated with Paclitaxel compared to lungs from mice bearing shMLL1 tumors treated with Paclitaxel. (i) Mouse body weight for in vivo Paclitaxel-shMLL1 study. All data in this figure was generated with MDA-MB-231 cells *in vivo* or embedded in 3D collagen gels.

**Supplementary Figure 11.**
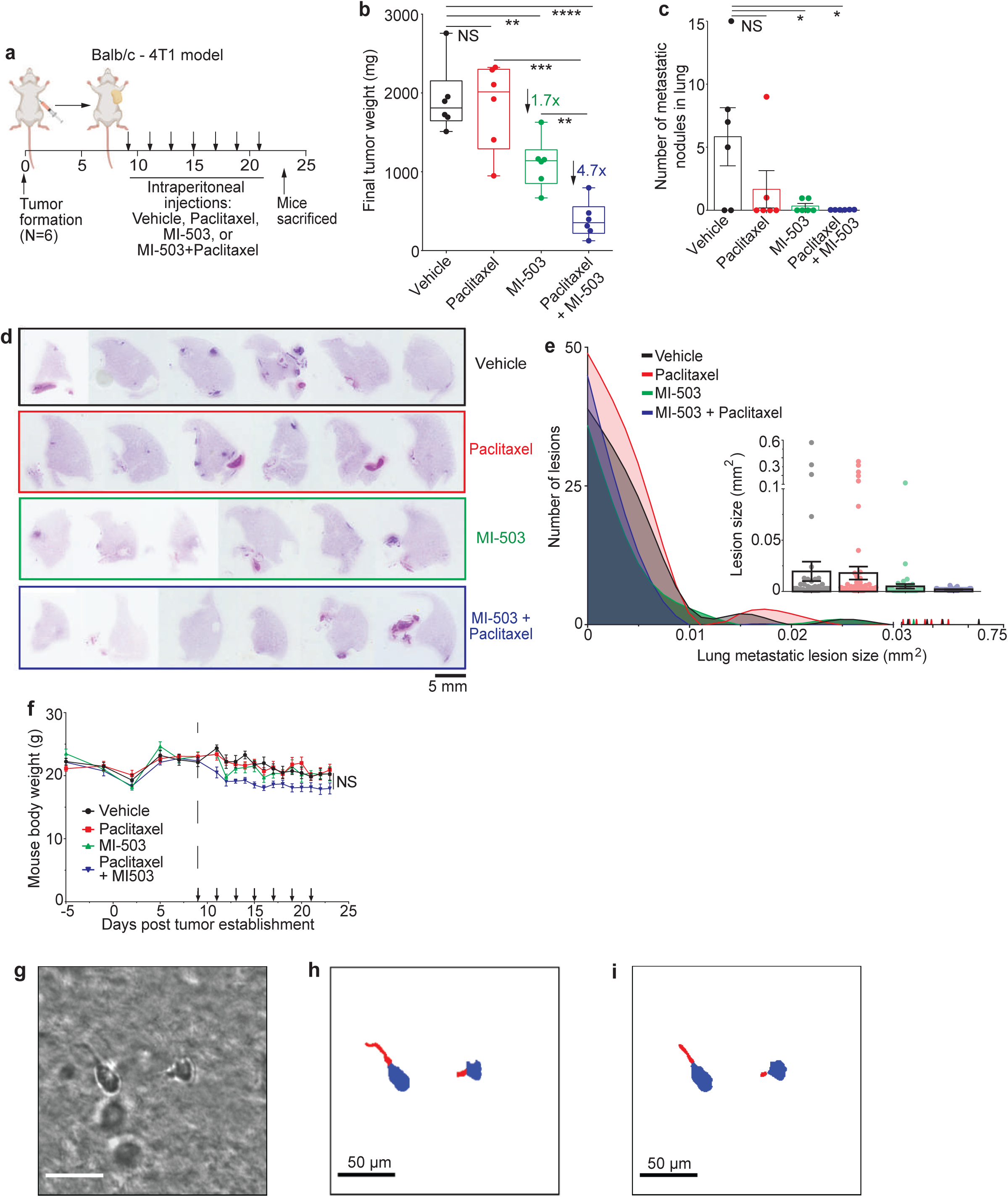
MLL1-Menin inhibition is additive with Paclitaxel in a syngeneic orthotopic model of TNBC. (a) Balb/c mice bearing syngeneic orthotopic 4T1 tumors were injected with MI-503, Paclitaxel, a combination of the two drugs, or a vehicle intraperitoneally every alternate day for seven times. (b) MI-503 treatment reduced tumor size and final tumor weights. Tumors treated with MI-503 + Paclitaxel were the lightest. (c) The number of lung metastatic nodules were counted upon excision. MLL1-Menin inhibition led to a dramatic decrease (over 10-fold) in metastatic burden, when administered by itself or in conjunction with Paclitaxel. (d) Full panel of lungs from 4T1-bearing mice. H&E staining of lungs show dark purple staining in mice that were administered vehicle. Administration of the MLL1-Menin inhibitor, MI-503, either by itself or in conjunction with Paclitaxel led to a decrease in lung metastases. (e) Histogram of metastatic lesions in the sample lungs shown in Fig. 7n. (f) Administration of MI-503, Paclitaxel, or a combination of the two did not lead to adverse toxicity as evident by a lack of significant decline in mouse body weight. (g) Input image, (h) the corresponding manual annotation and (i) the result of the segmentation. Cell body is in blue and cell protrusions are in red (scale bar 50 µm). Data in this figure was generated with 4T1 tumors in Balb/c mice (a-f) or MDA-MB-231 cells in 3D collagen gels (g-i).

**Supplementary Table 1:**
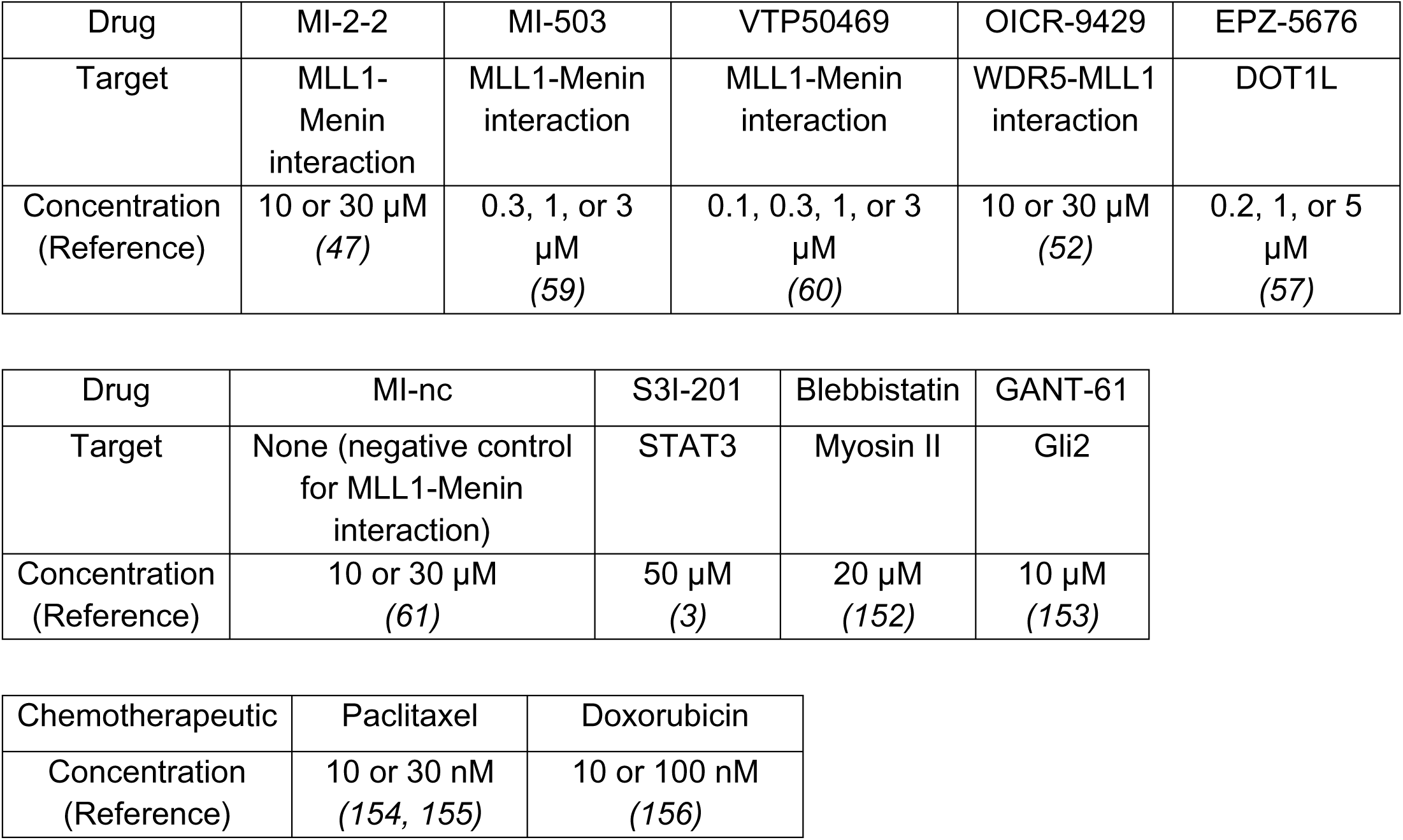
Concentrations for drugs used in this manuscript.

**Supplementary Table 2.**
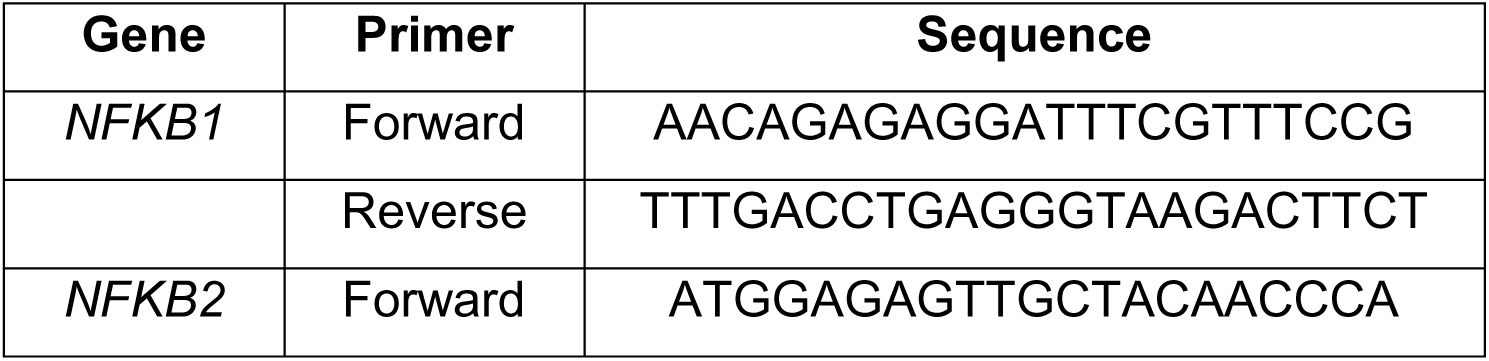
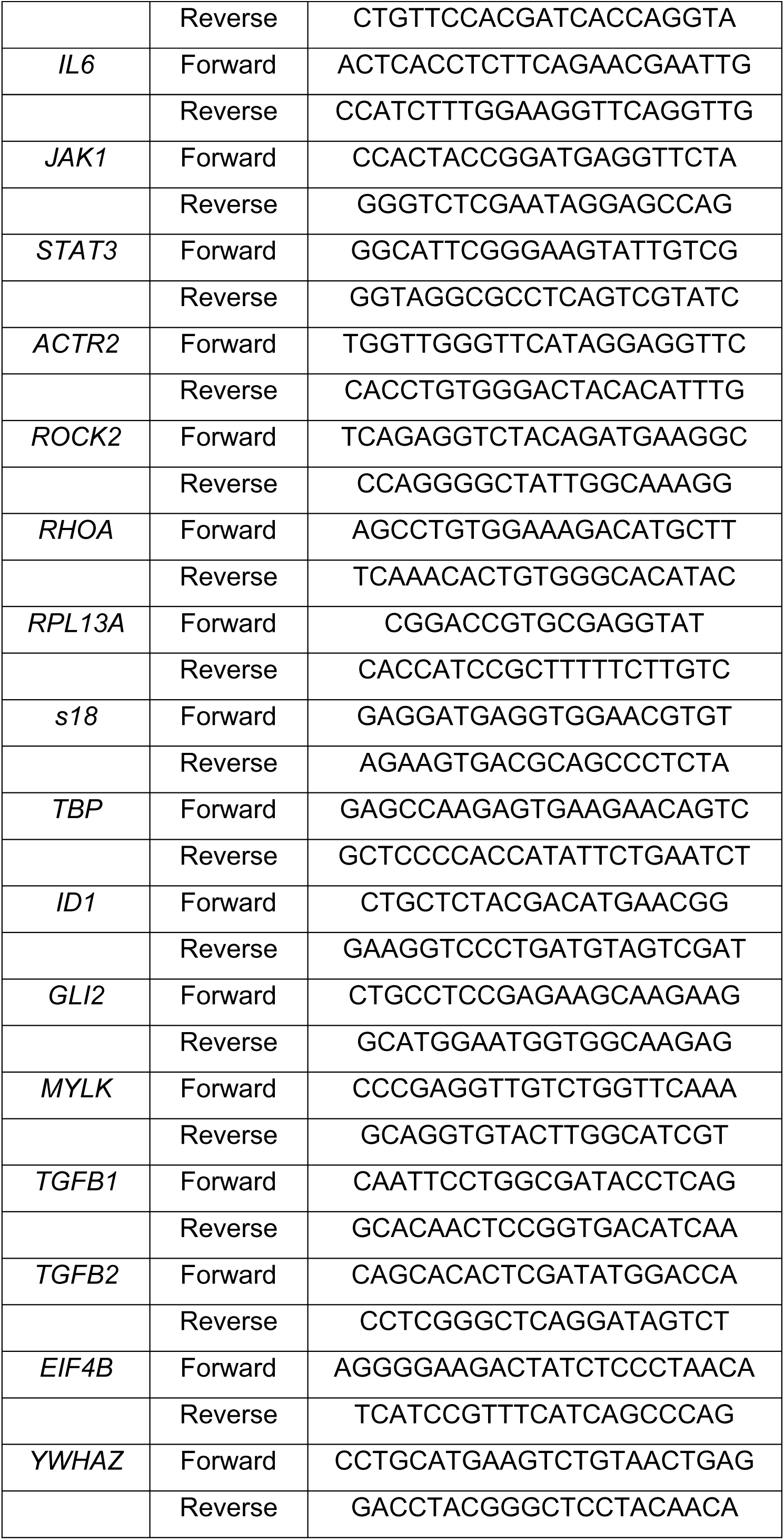
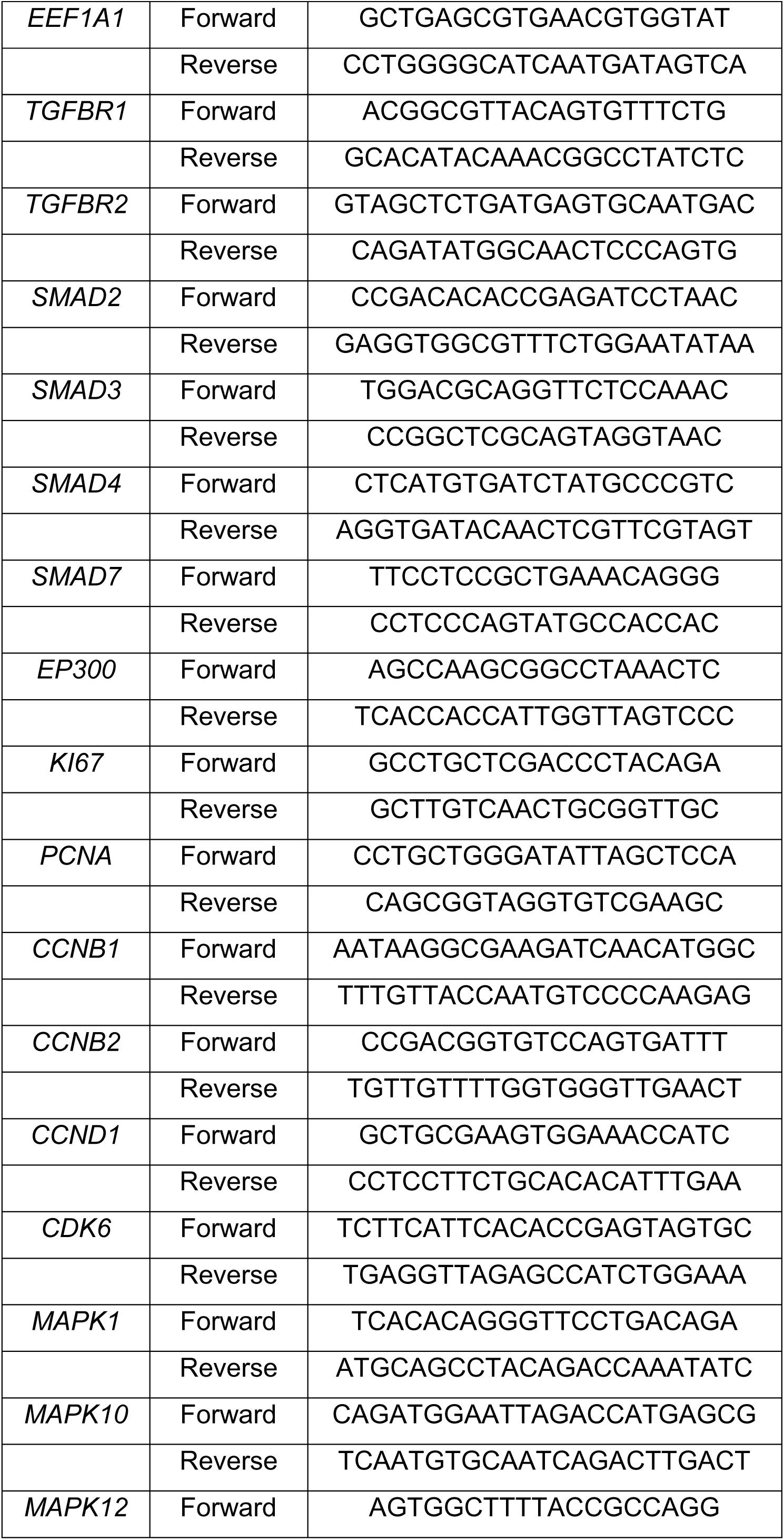
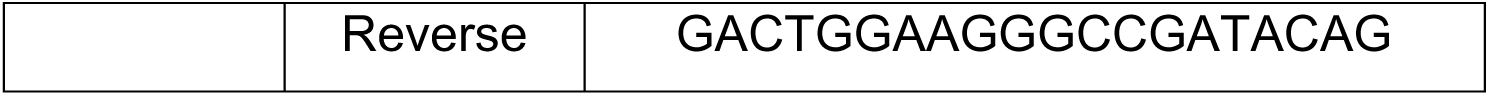
Sequences for qRT-PCR primers used.

**Supplementary Table 3.** Antibodies used for multiplex cytokine assay.

| Target Antigen | Vendor | Catalog Number |
| --- | --- | --- |
| IL-6 | R&D | DY206 |
| IL-8 | R&D | DY208 |
| TGF-β1 | R&D | DY240 |
| TGF-β2 | R&D | DY302 |

